# Evolutionary innovation within conserved gene regulatory networks underlying biomineralized skeletons in Bilateria

**DOI:** 10.1101/2025.08.27.672524

**Authors:** Yitian Bai, Yue Min, Shikai Liu, Yiming Hu, Shulei Jin, Hong Yu, Lingfeng Kong, Daniel J. Macqueen, Shaojun Du, Qi Li

## Abstract

Biomineralized skeletons have evolved convergently across animals, yet exhibit remarkable diversity in structure and development. However, the evolutionary origins of gene regulatory networks underlying biomineralized skeletons remains elusive. Here, we report comprehensive developmental profiling of transcriptomic and chromatin dynamics in a bivalve mollusc, *Crassostrea nippona*. We provide evidence for a biphasic regulatory program orchestrating larval and adult shell formation, involving the coordinated activity of ancient transcription factors and dynamic chromatin remodeling. Comparative analyses suggest a conserved developmental toolkit was co-opted for larval exoskeleton formation in the common lophotrochozoan ancestor. In contrast, limited regulatory conservation was observed between lophotrochozoans and echinoderms with regards to the formation of biomineralized skeletons, despite both relying on a heterochronic activation of ancestral regulators. Together, our findings support a hierarchical model where dynamic chromatin decouples rapidly evolving effectors from deeply conserved regulators, allowing modular innovations within conserved gene regulatory networks. This study highlights how epigenetic dynamics bridge evolutionary conservation and novelty, offering a framework for understanding the independent evolution of biomineralization across Bilateria through combinatorial regulatory evolution.

## Introduction

Many animals produce biomineralized skeletons for protection and physical support. This process represents a classic example of functional convergence, with distinct metazoan phyla independently evolving biomineralized skeletons multiple times since the early Cambrian^1^. Despite their divergent origins, biomineralized systems share strikingly similar strategies for utilizing organic matrices and amorphous precursors to construct diverse skeletal architectures^2–4^, including exoskeletons (e.g., coral skeletons, molluscan and brachiopod shells) and endoskeletons (e.g., sponge and echinoderm spicules, vertebrate bones) (Fig. 1a). Such convergence is supposedly derived from a conserved “biomineralization toolkit”, comprising a set of ancestral genes and regulatory elements co-opted across lineages^4^. However, the origins and homology of these toolkits and their associated gene regulatory networks (GRNs), both across and within phyla, remain poorly understood. Resolving these questions has implications for understanding the molecular mechanisms underlying the evolution of biological complexity and diversity.

**Fig. 1.**
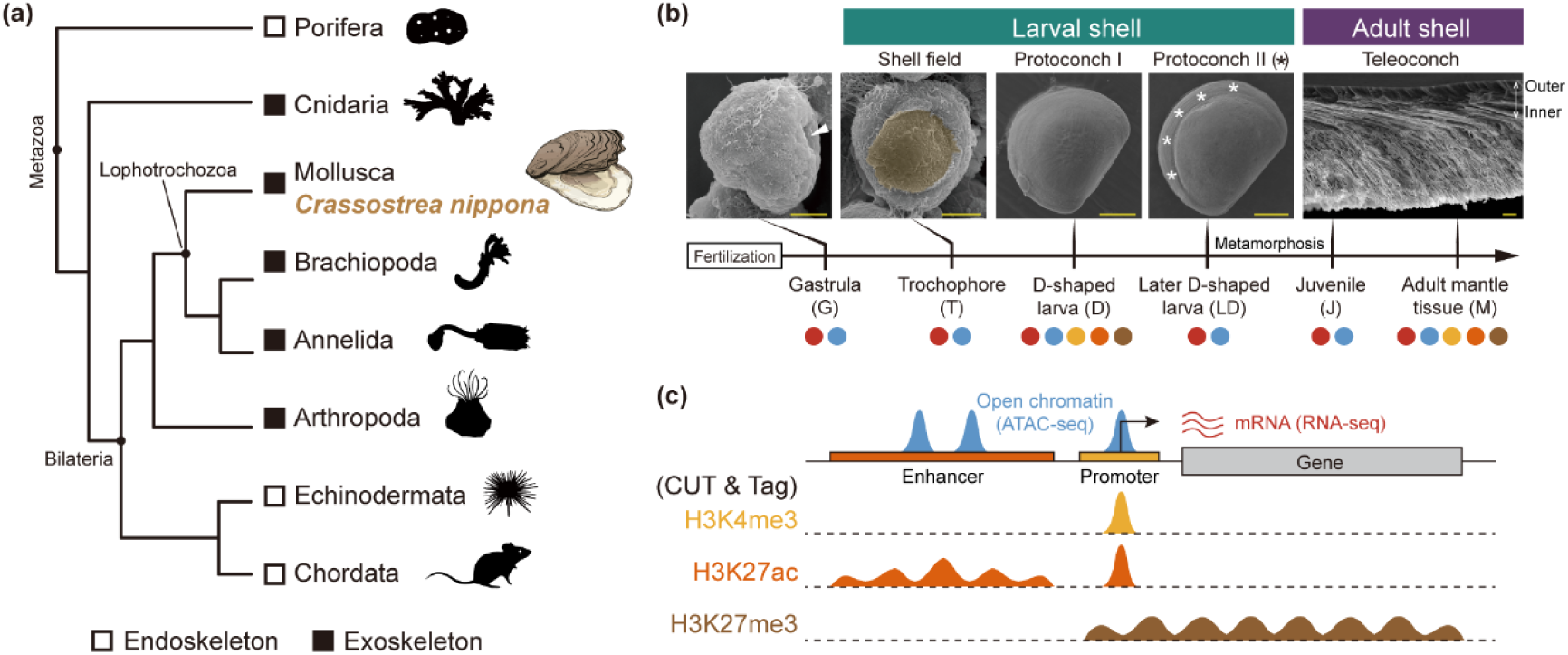
Diversity of biomineralized skeletons in Metazoans across phylogenetic lineages and developmental stages. **(a)** Schematic phylogeny illustrating the distribution of biomineralized skeletons among representative metazoan lineages. **(b)** Experimental design. Top: scanning electron micrographs (SEMs) of *C. nippona* shells at representative developmental stages. The blastopore (arrow) formed at the gastrula stage (8 hours post fertilization, hpf). At the trochophore stage (14 hpf), the shell field (yellow) covers the dorsal region. The uniform calcified shell surface of protoconch I was observed at the D-shaped stage (22 hpf). Growth striations (asterisks) of protoconch II appeared on the shell surface at 2 days post fertilization (dpf). The adult shell comprises three shell layers: outer prismatic layer, and inner foliated and chalky layers (Supplementary Fig. 1). Scale bar = 20 μm. Detailed observations of larval and adult shells are shown in Supplementary Fig. 1. Bottom: samples for RNA-seq (red), ATAC-seq (blue), and each histone modification (yellow for H3K4me3; orange for H3K27ac; brown for H3K27me3) are represented by color-coded dots. **(c)** A schematic illustrating representative peak signals of accessible chromatin and three histone modifications. The orange and yellow boxes indicate enhancer and promoter regions, respectively, while the grey box indicates the gene body.

Skeletal formation commonly initiates during early development (embryonic or larval stages) and exhibits ontogenetic plasticity, transitioning from transient larval frameworks to complex adult biomineralized architectures^5–9^. This dynamic remodeling reflects adaptive responses to ecological pressures and physiological demands throughout the life cycle^1^. Most studies have focused on echinoderms and chordates^10–13^, revealing the independent co-option of ancestral developmental GRNs for skeletogenesis across different deuterostome clades^7, 14–16^. Skeletogenic programs of deuterostomes appear to have evolved through the activation of phylum-specific regulatory and differentiation genes from modifications and rewiring of ancestral GRNs^16–18^. Yet, biomineralization GRNs remain largely underrepresented across non-deuterostome lineages and their life stages.

Mollusca is the second most diverse animal phylum, and its extraordinary morphological diversity is largely attributed to the remarkable variety of biomineralized shells^19^. A large number of rapidly evolving and co-opted genes, such as shell matrix proteins (SMPs), biomineralization enzymes, and transporters, have been identified as downstream effectors involved in shaping the diversity of adult shell structures^20–23^. Larval shells are more conserved than adult shells in terms of morphology, crystal polymorph and microstructure across lineages^5, 6, 24^. This phenotypic conservation likely results from deeply conserved upstream GRNs governing early shell development^25^. Supporting this hypothesis, recent work demonstrated an ancestral co-option of Hox genes in larval shell development programs across molluscan clades^26^. Intriguingly, proteomic comparisons revealed almost entirely distinct SMPs repertoires between larval and adult stages^24, 27, 28^, indicating that distinct sets of downstream effectors are deployed at different life stages, even within the same species. Nonetheless, it remains plausible that shared upstream GRNs, typically transcription factors (TFs), orchestrate skeletal formation across ontogeny^15, 16, 29^, with stage-specific modifications directing divergent downstream targets. Even though there have been efforts to predict GRNs for adult shell formation in molluscs such as *Laternula elliptica*^30^, the current lack of 1) epigenomic profiling of *cis*-regulatory elements and 2) shell formation GRNs spanning both larval and adult stages, limits understanding of the dynamics underlying molluscan biomineralization strategies across ontogeny and evolution.

To address these questions comprehensively, we profiled transcriptomic and genome-wide chromatin dynamics across multiple developmental stages of a bivalve mollusc, *Crassostrea nippona*, with a focus on early larval and adult shell formation (Fig. 1b, c and Supplementary Fig. 1). We discovered a novel biphasic regulatory program wherein stage-specific chromatin landscape transitions enable conserved upstream TFs to regulate distinct sets of downstream biomineralization genes in larvae and adults. By integrating transcriptomic, chromatin accessibility and TF footprinting analyses, we identify stage-specific GRNs, revealing an overlapping group of TFs that orchestrate this biphasic regulatory program. These TFs likely constitute a conserved molecular toolkit for exoskeleton formation in lophotrochozoan larvae, with some independently co-opted for early skeletogenesis in echinoderms. Despite both systems relying on the heterochronic activation of ancestral regulators, the formation of biomineralized skeleton in lophotrochozoans and echinoderms shows limited regulatory conservation.

## Results

### Transcriptomic dynamics during larval and adult shell formation

The oyster goes through several morphologically distinct developmental stages, with shell formation exhibiting major phenotypic differences: the shell field and protoconch I, II during early development (collectively referred to as larval shells in this study), and teleoconch after metamorphosis (referred to as the adult shell) (Fig. 1b and Supplementary Fig. 1). To obtain a reliable gene annotation across oyster development, we generated a new full-length transcriptome from mixed-stage mRNA samples covering six representative shell-forming stages in *C. nippona* using Iso-seq, resulting in the identification of 29,577 gene models (Supplementary Fig. 2 and 3, Supplementary Note 1, Supplementary Data 1). To investigate the global dynamics of gene expression during larval and adult shell formation, we performed RNA-seq at these stages (Fig. 1b, Supplementary Fig. 4a, b and Supplementary Data 1). We identified 25,465 genes expressed during at least one stage (transcripts per million, TPM, > 1), which were divided into ten clusters using *K*-means clustering (Fig. 2a). Gene ontology (GO) enrichment analysis of these clusters highlighted major developmental processes associated with shell formation (Supplementary Fig. 5, Supplementary Note 2, Supplementary Data 3). We found GO terms related to calcium ion transport and the extracellular matrix during the D-shape larva and adult stages (clusters C5, C6, C7, and C9), as well as peptidase inhibitor activity at the adult stage (cluster C9), suggesting the functional importance of these stages in shell formation^22, 31^.

**Fig. 2.**
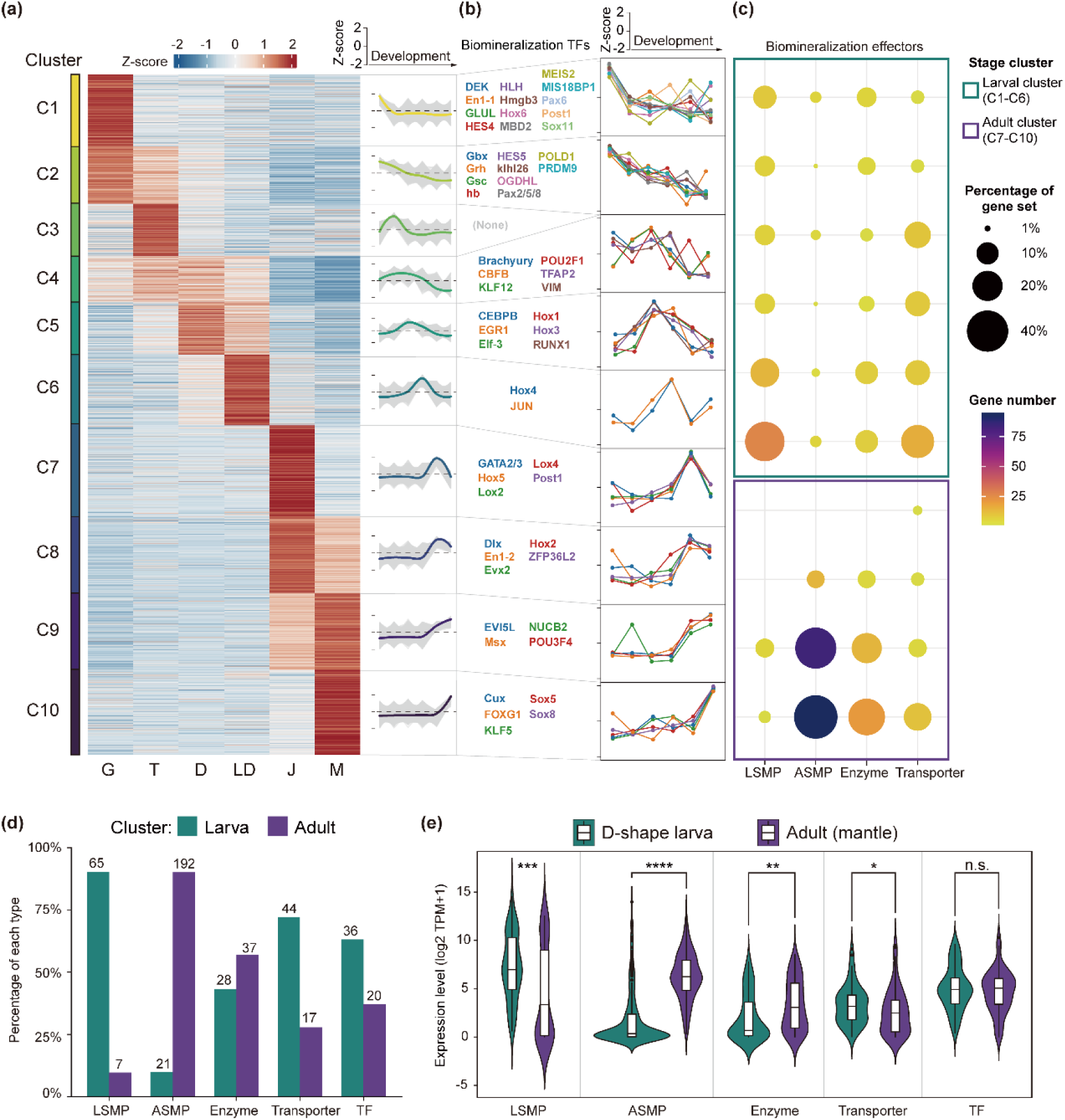
Transcriptomic atlas of biomineralization across *C. nippona* ontogeny. **(a)** K-means clustering of stage-specific highly expressed genes. On the right of heatmap, gene-wise expression dynamics (grey lines) and locally estimated scatterplot smoothing (colored lines) are shown for each cluster. Abbreviations: G: gastrula; T: trochophore; D: D-shaped larva; LD: later D-shaped larva; J: juvenile; M: adult mantle. **(b)** Expression patterns (right) of previously reported biomineralization TF genes involved in molluscan shell formation (left) in each cluster. Y axis represents Z-score normalized gene expression. **(c)** Distribution of highly expressed downstream biomineralization effector genes across clusters. Dot size represents the percentage of downstream effector genes relative to the total number of genes in each effector category. Color indicates the gene number. Green and purple boxes highlight larval (clusters C1-C6) and adult clusters (clusters C7-C10), respectively. Abbreviations: LSMP: larval shell matrix protein; ASMP: adult shell matrix protein. **(d)** Number and percentage of each category of biomineralization effector genes within larval and adult clusters. The numbers on each bar indicate gene count. Abbreviations as for part **a**. **(e)** Comparison of expression levels for each effector gene category between D-shape larva and adult stages. Boxplots include the median with quartiles and outliers above the top whisker. A two-sided Wilcoxon rank-sum test was used to assess significance (\**P* < 0.05; \*\**P* < 0.01; \*\*\**P* < 0.001; \*\*\*\**P* < 0.0001; n.s., not significant). Abbreviations as for part **a**.

To explore the transcriptional programs governing the expression of biomineralization genes at larval and adult stages, we identified 56 biomineralization TF genes implicated in molluscan shell formation (Supplementary Data 4), and 382 biomineralization effector genes (Supplementary Data 5), including larval shell matrix proteins (LSMPs), adult shell matrix proteins (ASMPs), biomineralization enzymes, and transporters (Methods). Biomineralization TF genes generally exhibited ubiquitous expression across stages (Fig. 2b and Supplementary Fig. 4c). In contrast, effector genes showed stage-specific expression during larval and adult shell formation (Fig. 2c, d, Supplementary Fig. 6, 7 and Supplementary Data 5). Most LSMP genes (90.3%) were enriched in larval clusters and highly expressed during larval shell formation (Fig. 2c, d and Supplementary Fig. 6a), while ASMP genes (90.1%) were predominantly grouped into adult clusters and showed high expression level during adult shell formation (Fig. 2c, d and Supplementary Fig. 6b). Similarly, genes encoding biomineralization enzymes and transporters showed distinct expression between larval and adult stages (Fig. 2c, d and Supplementary Fig. 7a).

Thus, transcriptional programs regulating biomineralization effector genes involved in shell formation are temporally regulated at distinct life stages^24, 27^, resulting in the morphological divergence between larval and adult shells. Strikingly, despite the significant stage-specific expression differences of effector genes, particularly between the D-shape larva and adult stages, TF genes maintain relatively stable expression levels with no significant variation across developmental stages (Fig. 2e, Supplementary Fig. 7b, c and Supplementary Data 6). Therefore, we hypothesized that these TFs play a foundational role in controlling two major transcriptional programs that separately drive larval and adult shell formation.

### A biphasic regulatory program for larval and adult shell formation

To delve into the transcriptional regulation of larval and adult shell formation, we applied the assay for transposase-accessible chromatin sequencing (ATAC-seq) to the six stages used for RNA-seq (Fig. 1b, c and Supplementary Data 7). Larval and adult stages showed a distinct genome-wide chromatin accessibility landscape (Supplementary Fig. 8). 165,946 open chromatin regions were detected across all sampled stages, with on average 12.9%, 51.5% and 29.7% overlapping promoter (- 2kb + 0.5kb transcription start site; TSS), genic (+ 0.5kb TSS to transcription end site; TES) and intergenic regions, respectively (Fig. 3a), consistent with previous studies in the oyster^32^. There was, however, an increase in promoter peaks in adults compared to larvae (Fig. 3b). Moreover, we observed that adult mantles exhibited more differentially accessible regions relative to the D-shape larvae, than to other larval stages, suggesting the existence of distinct *cis*-regulation programs between these two stages (Fig. 3c).

**Fig. 3.**
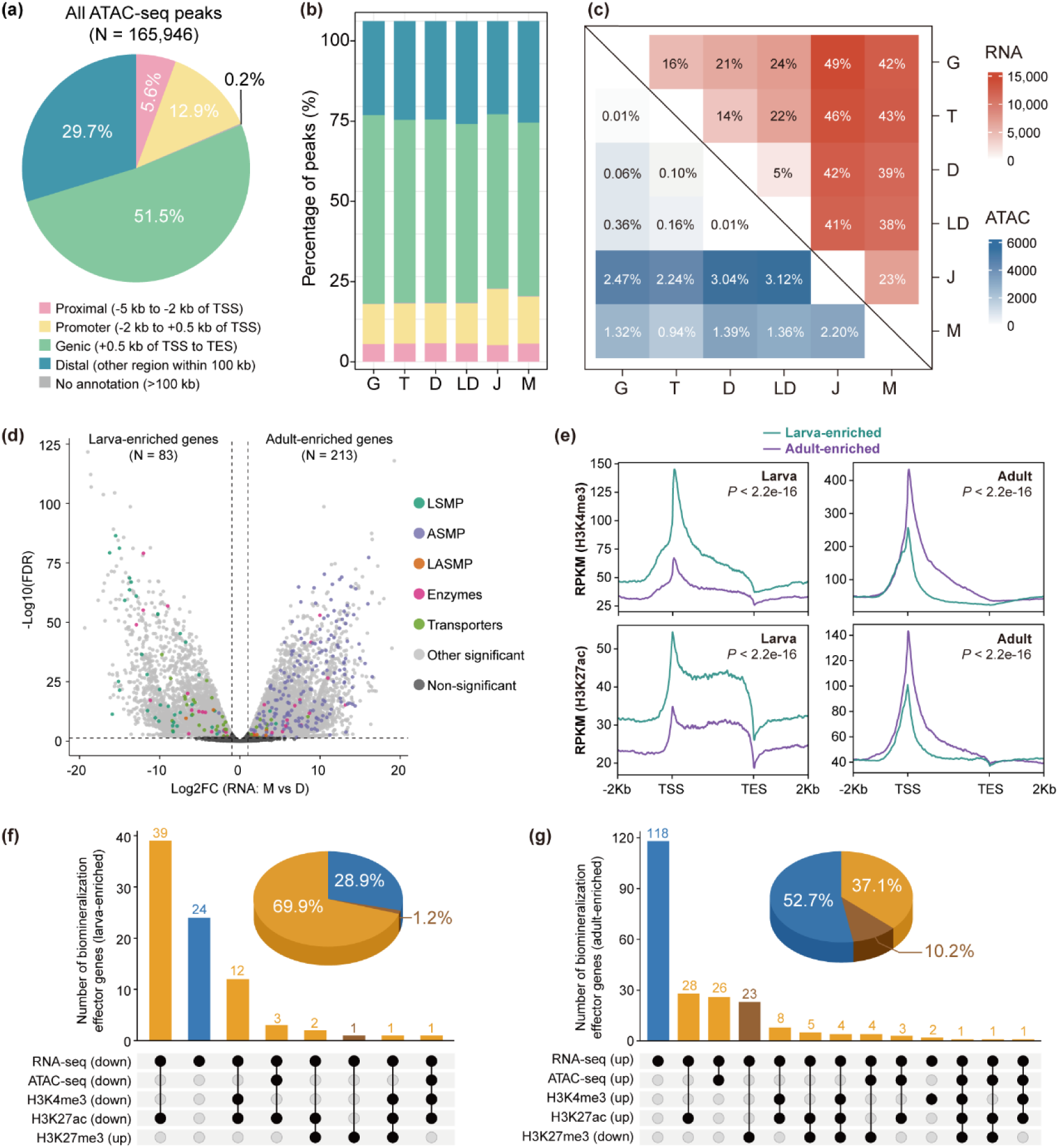
Dynamic chromatin changes underpin the expression of biomineralization effector genes during larval and adult shell formation. **(a)** Genomic feature annotation of the consensus ATAC-seq peaks. **(b)** Stacked bar plots showing the proportion of called peaks per developmental stage classified by genomic feature. **(c)** Number and proportion of differentially expressed genes (DEGs) and differentially accessible peaks. **(d)** Differential gene expression between D-shape larvae and adult mantles, with biomineralization effectors highlighted dots in color. Genes significantly up-regulated in adult mantles were defined as adult-enriched, while those down-regulated were considered as larva-enriched. Detailed information is shown in Supplementary Data 5. **(e)** H3K4me3 and H3K27ac levels in genomic regions containing larva-enriched or adult-enriched genes, shown for larval (left) and adult (right) stages. Statistical significance of histone modification levels around larva-enriched or adult-enriched genes (from 5 kb upstream to the TES) was assessed using a two-sided Wilcoxon rank-sum test. **(f)** Upset plots showing the overlap between larval-enriched biomineralization effector genes and corresponding chromatin features (from 5 kb upstream to the TES), characterized by significantly increased chromatin accessibility and active histone modifications (H3K4me3 and H3K27ac) (yellow) in the adult, compared to larvae, accompanied by a decrease in the repressive mark H3K27me3 (brown). **(g)** Upset plots showing the overlap between adult-enriched biomineralization effector genes and corresponding chromatin features (from 5 kb upstream to the TES) that exhibit significantly decreased chromatin accessibility and active histone modifications (H3K4me3 and H3K27ac) (yellow) in the adult, relative to those of larvae, along with repressive mark H3K27me3 increasing (brown). Pie charts in **f** and **g** show the proportion of larval-enriched **(f)** and adult-enriched **(g)** biomineralization-related effector genes associated with differential chromatin features (yellow for open chromatin, H3K4me3 and H3K27ac; brown for H3K27me3) between larvae and adults, compared to those without differential chromatin changes (blue). Detailed information is shown in Supplementary Data 8.

Based on these observations, we selected D-shape larvae and adult mantles respectively as representative samples for larval and adult shell formation, and performed cleavage under targets and tagmentation (CUT&Tag) profiling for three types of histone modification H3K4me3, H3K27ac and H3K27me3) (Fig. 1b, c and Supplementary Data 8). As expected, H3K4me3 peaks were highly enriched near gene TSSs and most abundant within promoter regions, whereas H3K27ac and H3K27me3 marks were more broadly located across genes, with enrichment towards the TSS (Supplementary Fig. 9a, b). We also found that genes with relatively high expression levels were on average marked by more H3K27ac and H3K4me3, and less H3K27me3, except for H3K27me3 in D-shape larvae (Supplementary Fig. 9c, d). The positive correlation between H3K27me3 and gene expression in larvae may reflect high cellular heterogeneity or unresolved long-range regulatory interactions in the larval samples. Differential gene expression between D-shape larvae and adult mantles was significantly and positively correlated with changes in H3K4me3 and H3K27ac levels (Pearson correlation coefficient R = 0.724 and 0.422, respectively; *P* < 2.2e-16) (Supplementary Fig. 9e-g).

We next identified adult-enriched (highly expressed in adult mantles) and larva-enriched (highly expressed in D-shape larvae) genes based on differential expression analysis (|log_2_foldchange| > 1, false discovery rate corrected *P*-value (FDR) < 0.05) (Fig. 3d). Both larva- and adult-enriched genes followed similar trends of H3K4me3 and H3K27ac enrichment at TSS, consistent with their roles as canonical markers of transcriptional activation. Notably, these modifications showed stage-specific patterns, with larva- and adult-enriched genes associated with high levels of both H3K4me3 and H3K27ac during the larval and adult stages, respectively (*P* < 2.2e-16) (Fig. 3e). In total, 83 and 224 biomineralization effector genes were identified as larva and adult-enriched genes, respectively (Fig. 3d). Among them, 59 (71.1%) larva-enriched and 106 (47.3%) adult-enriched biomineralization effector genes correlated with significant changes in either chromatin accessibility or at least one histone mark (Fig. 3f, g and Supplementary Data 9). Importantly, the majority of these genes exhibited regulatory changes linked to open chromatin regions, H3K4me3 or H3K27ac, accounting for 69.9% of larva-enriched genes and 37.1% of adult-enriched genes. These findings imply H3K4me3 and H3K27ac as critical histone marks for transcriptional activation during larval and adult shell formation.

The chromatin states of larvae and adults were systematically characterized using ChromHMM^33^, which integrated combinatorial patterns of multiple epigenomic marks (Fig. 4a and Supplementary Fig. 10a). For simplicity, the eight identified chromatin states were consolidated into five major categories used throughout this study (Fig. 4a), according to previous studies^34, 35^. Genomic regions marked by accessible chromatin, H3K4me3 and H3K27ac were classified as active promoters, typically showing biased distribution around the TSS (Fig. 4b), whereas those enriched for open chromatin and H3K27ac, but lacking H3K4me3, were categorized as active enhancers. Together, these regions were defined as activate elements (Act). Regions characterized by high H3K27me3 levels and low levels of chromatin accessibility were classified as repressed elements (Rep), while those marked by both H3K4me3 and H3K27me3 features were defined as poised/bivalent states (Pois). Bulk-level profiling may capture overlapping signals from heterogeneous samples and cell populations; thus, co-occurrence of H3K27ac and H3K27me3 does not imply bivalency within individual cells. Regions marked exclusively by open chromatin were classified as accessible islands (ATAC), and those with no epigenomic signals were quiescent states (Qui). As expected, poised and active elements (Pois and Act) showed high gene expression levels (Fig. 4c). In contrast, repressed (Rep) and the low signal regions (Qui) had relatively low gene expression levels (Fig. 4c). Furthermore, biomineralization effector genes linked to poised and active states (Pois and Act) exhibited significantly higher expression levels compared to those remaining in other states (Rep, ATAC and Qui) in both larval and adult stages (Fig. 4d). Effector genes associated with active promoters and enhancers (Act) showed the most significant differences (*P* = 2.7e-16) in expression levels between larvae and adults, indicating a regulatory role of these elements in larval and adult shell formation. We next investigated epigenomic transitions from larval to adult stages, and found widespread switching among chromatin states across the genome (Supplementary Fig. 10b, c).

**Fig. 4.**
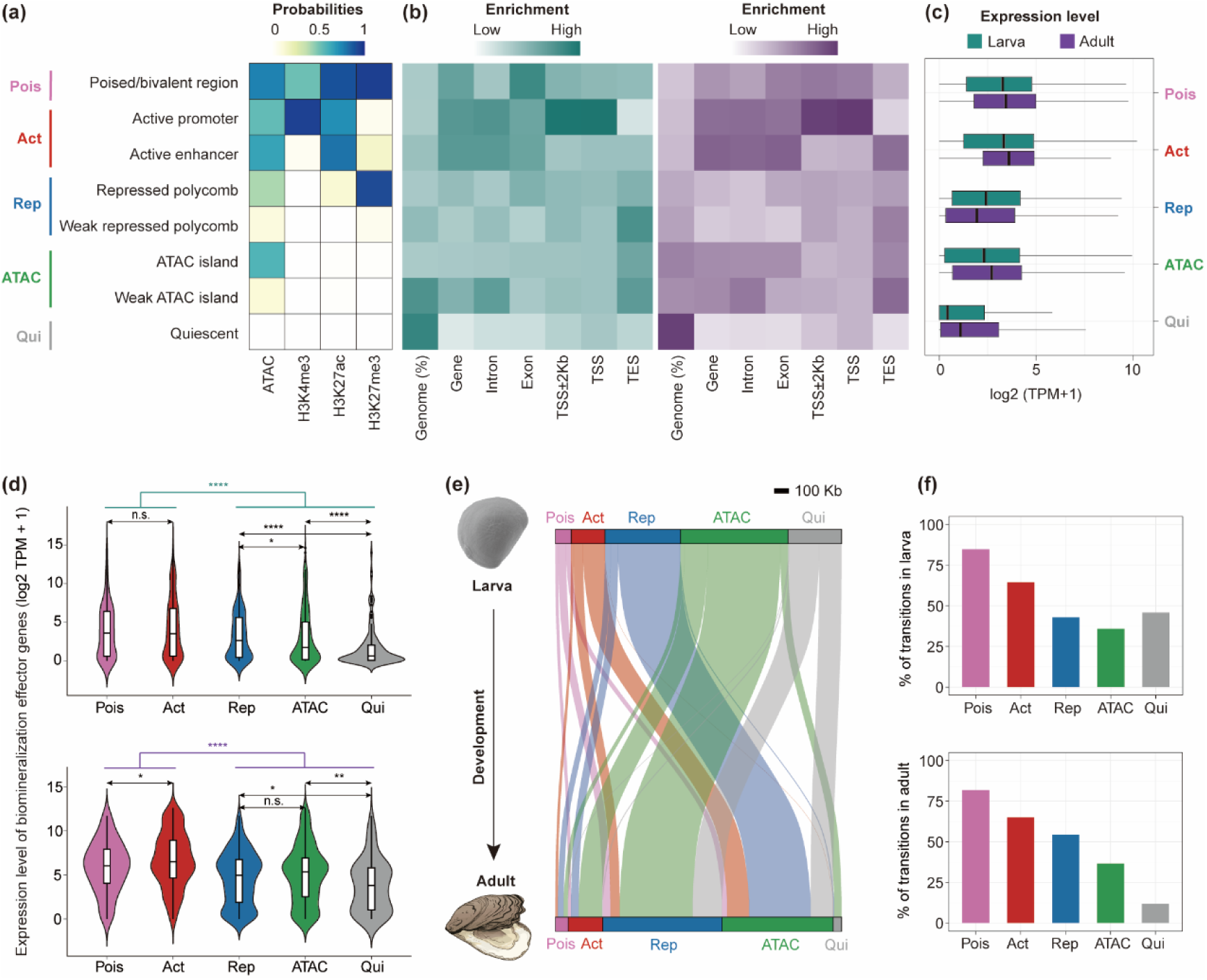
Dynamic remodeling of chromatin states across larval and adult stages orchestrates the regulation of biomineralization effector genes during shell formation. **(a)** ChromHMM eight chromatin state models grouped into five categories: poised (Pois), active (Act), repressed (Rep), accessible chromatin (ATAC), and quiescent (Qui) states, based on ATAC-seq and histone modification profiles. **(b)** Genomic feature enrichments for each chromatin state at larval and adult stages. **(c)** Expression levels of genes with each chromatin state across both stages. Error bars represent the median with quartiles. **(d)** Gene expression levels of biomineralization effectors with each chromatin state at larval (top) and adult (bottom) stages. Error bars represent the median with quartiles. Two-sided Wilcoxon tests were used for comparisons between states (\**P* < 0.05; \*\**P* < 0.01; \*\*\*\**P* < 0.0001; n.s., no significance). **(e)** Sankey diagram showing chromatin state transitions of biomineralization effector genes from larva (top) to adult (bottom) stages. **(f)** Top: percentage of chromatin states linked to biomineralization effector genes in larvae that undergo transitions to other chromatin state in adults. Bottom: percentage of chromatin states linked to biomineralization effector genes in adult that undergo transitions from other chromatin state in larvae.

Focusing on regulatory elements associated with biomineralization effector genes, a large proportion of poised (84.8%) and active (64.4%) regions in larvae transitioned to alternative chromatin states in adults, whereas most poised (81.8%) and active (65.0%) regions observed in adults originated from other chromatin states in larvae (Fig. 4e, f). These results indicated that oyster regulatory programs for shell formation were extensively rewired between life stages, with regulatory elements associated with biomineralization effector genes selectively activated to meet stage-specific requirements of larval or adult shell formation. Such dynamic remodeling establishes novel chromatin landscapes that enable the activation of adult-specific biomineralization programs.

Taken together, dynamic changes of chromatin landscape mediate the epigenomic regulation of biomineralization effector genes, and establish a biphasic regulatory program controlling larval and adult shell formation. This program comprises two temporally distinct epigenetic phases: an initial phase during larval shell-forming stages, where active histone modifications, particularly H3K4me3 and H3K27ac, facilitate the transcriptional activation of larva-enriched biomineralization effector genes for larval shell formation, followed by an adult phase characterized by chromatin remodeling that regulates the expression of adult-enriched genes for adult shell construction. These two phases coordinate stage-specific gene regulation, driving the developmental and functional complexity of shell biomineralization across oyster ontogeny.

### Functional divergence of biomineralization paralogs for larval and adult shell formation

Gene duplication is a major driver of molecular innovation in shell biomineralization, facilitating the emergence of novel or specialized functions among paralogous biomineralization genes in molluscs^36–38^, often accompanied by changes in cis-regulatory elements^39^. To explore the evolutionary origins and divergence times of the paralogous genes involved in larval and adult shell formation, we identified 188 paralogous genes encoding biomineralization effectors using phylogenetic inference (Fig. 5a and Supplementary Data 10). Phylostratigraphic analysis^40^ revealed that most (91.5%) of these genes originated before the emergence of molluscs (pre-molluscan origin), whereas their paralog duplication events occurred predominantly at or after the origin of molluscs, thus representing mollusc-specific gene expansions for shell formation (Fig. 5b, Supplementary Fig. 11a and Supplementary Data 10). In addition, this phenomenon was also observed in other molluscan species (Supplementary Fig. 11b and Supplementary Data 11). To investigate the functional specialization of biomineralization effector genes between larval and adult shell formation, we classified these biomineralization effector genes into three groups (Supplementary Data 10), based on differential expression analysis between larval and adult stages in *C. nippona* (Fig. 3d). Larva-enriched and adult-enriched biomineralization effector genes were respectively involved in larval and adult shell formation, whereas genes without significant differential expression likely function in both stages (Supplementary Data 10). This stage-specific expression framework provides a valuable basis for dissecting the developmental and regulatory mechanisms underlying shell formation. Interestingly, 193 (47.0%) of the analyzed paralog gene pairs exhibited divergent expression patterns between larval and adult stages, indicating functional divergence. Of these, 62.2% resulted from duplication events specific to molluscan lineages (Fig. 5c and Supplementary Data 10), suggesting that the functional divergence of biomineralization genes between larval or adult shell formation may represent molecular innovation during mollusc evolution.

**Fig. 5.**
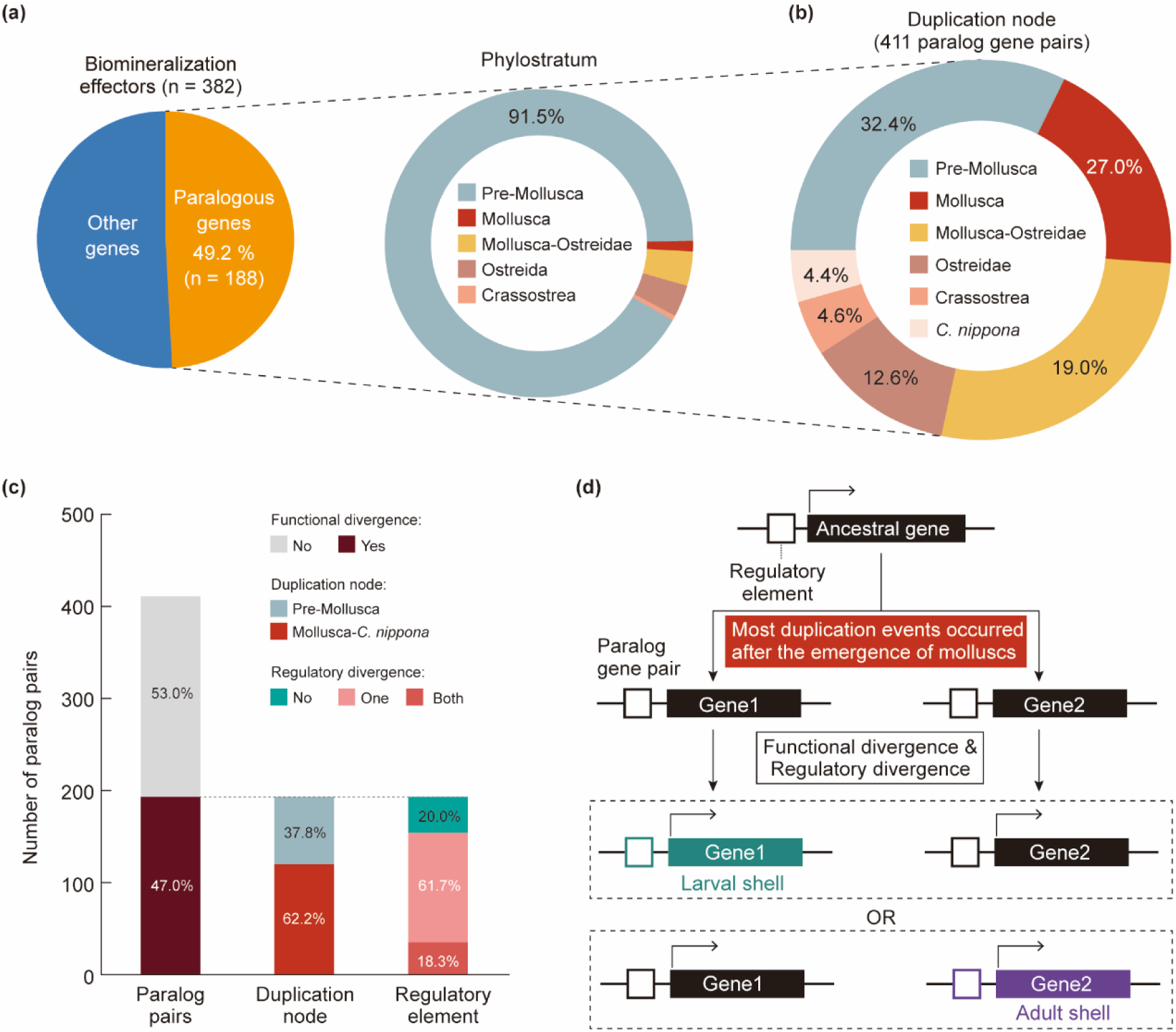
Functional specialization of biomineralization paralogs in larval or adult shell formation. **(a)** Percentage of paralogous genes in biomineralization effector genes, and the distribution of their phylostrata. **(b)** Duplication nodes of paralog gene pairs. **(c)** Percentage of paralog gene pairs with functional divergence, and the distribution of duplication nodes and regulatory element divergence among them. **(d)** Schematic representation of paralog specialization following gene duplication. After a mollusc-specific duplication from an ancestral gene, one paralog retained the ancestral regulatory program and function, while the other acquired distinct regulatory elements and functions, driving specialization for larval or adult shell formation.

To further assess the underlying regulatory basis for this functional divergence, we investigated active chromatin marks (H3K4me3 and H3K27ac) associated with each paralog in larvae and adults. The regulatory divergence among functionally specialized paralog pairs was categorized into three types: no divergence (both genes sharing similar regulatory activity across stages), divergence in one gene of the pair, and divergence in both genes (Supplementary Note 3, Supplementary Data S10). Strikingly, among paralog pairs that had already diverged in expression patterns, the majority (61.7%) displayed regulatory divergence in a single paralog, while few fraction (18.3%) showed divergence in both genes (Fig. 5c). This asymmetric regulatory evolution suggests that, in many cases, one paralog retains the ancestral regulatory program and potentially the ancestral function, while the other acquires novel stage-specific regulatory elements, such as active elements only in larval or adult tissues, leading to functional innovation (Fig. 5d). Such one-sided acquisition of regulatory activity may facilitate rapid neofunctionalization, as the newly specialized paralog can adopt novel roles without compromising the original function preserved in its counterpart. These findings indicate that changes in cis-regulatory architecture, particularly the stage-specific gain of active chromatin regions in a single paralog, are the major mechanism driving the functional specialization of biomineralization effector genes during molluscan evolution.

### Inferred regulatory networks supporting conserved TFs in larval and adult shell formation

The coordination between transcription and active chromatin landscape suggests that *cis*-motif and *trans*-factors mediating GRNs could govern shell formation. To bridge TFs and target motifs, we identified 1,123 TFs genome-wide in *C. nippona* and assigned 312 of them to transcription factor binding sites (TFBSs) derived from the JASPAR database (Supplementary Data 12). By integrating the ATAC-seq footprinting and analysis algorithm for networks specified by enhancers (ANANSE) ^41^ with predicted *cis*-motifs, we then constructed the GRNs for larval and adult shell formation (Supplementary Fig. 12a). Two independent networks, one for larvae and one for adult, were generated, consisting of nodes (TFs and effectors) and edges (the interactions between TFs and target genes) (Supplementary Fig. 12b and Supplementary Data 13). In total, 239 putative TFs were identified, with 220 and 196 upstream TFs present in the larval and adult networks, respectively (Fig. 6a, Supplementary Fig. 12b). Notably, a total of 177 TFs (74.1% of all putative TFs) were common to both networks, including 29 TFs previously validated by ISH or single-cell RNA-seq (Fig. 6a and Supplementary Data 4). These shared TFs were expressed during both larval and adult stages, with no significant differences between larvae and adult mantles (Supplementary Fig. 13a, b). By contrast, TFs specific to the larval or adult GRNs exhibited higher expression levels during the corresponding stages (Supplementary Fig. 13c). Together, these results support that a largely conserved set of upstream TFs orchestrates distinct transcriptional programs for larval and adult shell formation in the oyster.

**Fig. 6.**
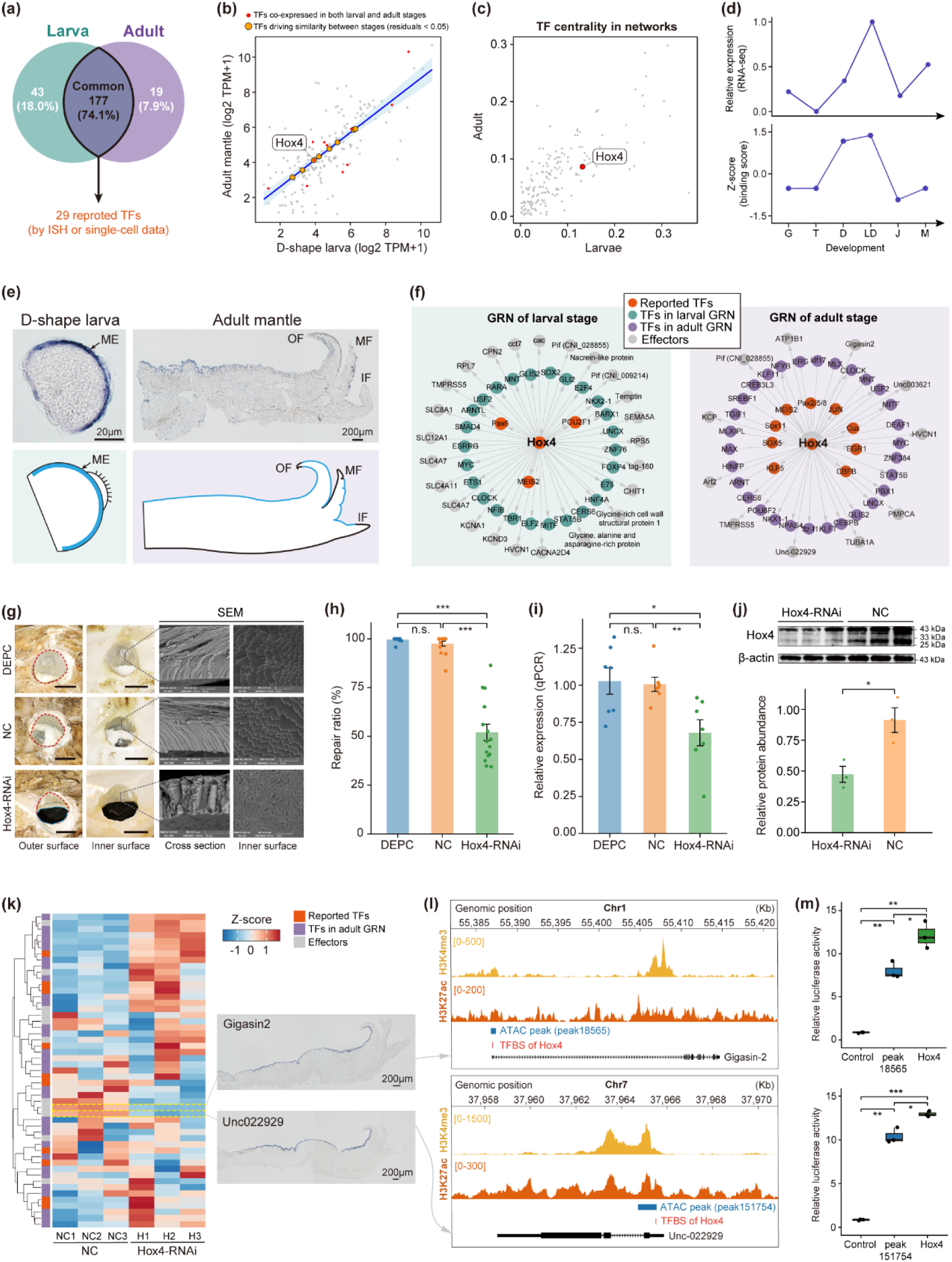
Conserved biomineralization TFs regulating both larval and shell formation in oysters. **(a)** Venn diagram showing biomineralization TFs in larval (green) and adult (purple) shell formation GRNs, with 29 reported TFs (orange) conserved across stages. **(b)** Scatter plot of gene expression of biomineralization TFs between D-shape larvae and adult mantles. A weighted least squares (WLS) regression model accounts for heteroscedasticity. The blue line indicates the ordinary least squares regression fit. Orange points represent candidate TFs with consistent expression relationships between stages (|residual| < 0.05). Red points highlight co-expressed TFs from Supplementary Fig. 13h. **(c)** TF centrality in larval (x axis) and adult (y axis) GRNs, with *Hox4* highlighted in red. **(d)** Expression and binding scores of *Hox4* across six stages/tissues. Abbreviations: same as (Fig. 1b). **(e)** ISH (top) and schematic (bottom) showing *Hox4* expression in D-shaped larvae and adult mantle. Abbreviations are the same as Fig. 3i. **(f)** GRNs showing *Hox4*-regulated biomineralization genes in larvae (left) and adult (right). **(g)** Shell repair six days after drilling, following injections of DEPC water (DEPC), negative control siRNA (NC), or *Hox4*-targeting siRNA. Left: bright-field photographs showing drill holes (red dashed) and unrepaired regions (blue dashed). Scale bar = 2 mm. Right: SEMs. **(h)** Shell repair ratio (repaired/hole area) in each group (n = 15; mean ± SE; two-sided Student’s t-test: \*\*\**P* < 0.001; n.s., no significance). Full images are shown in Supplementary Fig. 14. **(i)** *Hox4* expression in mantle tissues following RNAi experiments (n = 7; mean ± SE; two-sided Student’s t-test: \**P* < 0.05; \*\**P* < 0.01; n.s., no significance). **(j)** Western blot analysis of Hox4 protein abundance in mantle tissues following RNAi experiments (n = 3; mean ± SE; two-sided Student’s t-test: \**P* < 0.05). **(k)** Heatmap showing expression of *Hox4*-regulated downstream biomineralization genes after RNAi (n = 3), with spatial expression in the mantles of two down-regulated genes (right). **(l)** Regulatory activity around *Gigasin2* (top) and *Unc022929* (bottom), where regulatory elements located at 5’ UTRs contain predicted *Hox4* binding sites (TFBS). **(m)** Dual-luciferase assays validating *Hox4*-mediated activation of *Gigasin2* (top) and *Unc022929* (bottom) in human 293T cells (n = 3; two-sided Student’s t-test: \**P* < 0.05; \*\**P* < 0.01; \*\*\**P* < 0.001). Control: pGL3-basic; Peak: pGL3 + peak; Hox4: pGL3 + peak + *Hox4*.

We reasoned that the diversification of larval and adult shells might be attributed to the duplication and functional divergence of paralogous genes during molluscan evolution. However, phylostratigraphic analysis revealed that all upstream biomineralization TFs are of pre-molluscan origin, with their paralog duplication events predominantly occurring prior to molluscan emergence (Supplementary Fig. 13d, e). Thus, biomineralization TFs are likely evolutionarily ancient components that were co-opted to regulate recently evolved effector genes for molluscan shell formation. Moreover, GO enrichment analysis revealed that TFs in the two GRNs were not only enriched for terms related to skeletal system development and morphogenesis, but also for body plan pattern specification, cell fate determination, and other ancient early developmental processes (Supplementary Fig. 14). To explore the potential co-option of early developmental TFs into shell formation, we investigated the transcriptional similarity of common TFs between the D-shape larval and adult stages (Fig. 6b). Given that these TFs regulated distinct transcriptional programs in larval and adult shell, we also performed weighted correlation network analysis (WGCNA) ^42^ separately on developmental stages and adult tissues, identifying 24 co-expressed TFs (Supplementary Fig. 15). Notably, we observed that *Hox4* was among the set of TFs with conserved expression across both D-shape larva and adult stages (Fig. 6b and Supplementary Fig. 15h), and showed mantle-specific expression among adult tissues (Supplementary Fig. 15g). *Hox4* was also one of the most central TFs in larval and adult shell formation GRNs (Fig. 6c). Consistent with its previously reported role in early axial patterning and larval shell formation in molluscs^26^, *Hox4* was overrepresented and upregulated on the basis of its binding score at D-shape larva stage (Fig. 6d), and was spatially expressed in the mantle edge of D-shape larvae and epithelial cells of the adult mantle in *C. nippona* (Fig. 6e and Supplementary Fig. 16). In addition, *Hox4* was predicted to regulate distinct sets of downstream effector genes and TFs in larvae and adults, which were involved in the larval and adult shell formation, respectively (Fig. 6f).

To further confirm the function of *Hox4* in shell formation, we performed RNA interference (RNAi) experiments in adult oysters during shell repair. Compared to the control groups (DEPC and NC groups), the inner surfaces of foliated layer of the repaired shells showed irregular growth in the *Hox4*-RNAi groups (Fig. 6g). Furthermore, the *Hox4*-RNAi oysters exhibited a significantly lower shell repair ratio than the control oysters (*P* < 0.001) (Fig. 6h and Supplementary Fig. 17). Both gene expression levels and protein abundances of Hox4 were also significantly decreased in the mantles of *Hox4*-RNAi oysters compared with those of controls (*P* < 0.05) (Fig. 6i, j). Transcriptomic analysis of the oyster mantles after RNAi experiments revealed a significant down-regulation of two downstream SMP genes (*Gigasin2* and *Unc022929*) in the Hox4-mediated GRN (Fig. 6f, k and Supplementary Fig. 18). These genes were expressed in the epithelial cells of the adult mantle in *C. nippona* and potentially associated with shell formation (Fig. 6f). Finally, dual-luciferase reporter assay further validated that Hox4 regulates *Gigasin2* and *Unc022929* by binding to their promoter regions (Fig. 6l, m and Supplementary Data 14). Overall, these results suggest that the ancient *Hox4* gene, with biological functions related to early development^26, 43, 44^, has been co-opted for oyster shell formation by regulating the expression of downstream SMP genes.

### A conserved TF toolkit for larval exoskeleton formation in lophotrochozoans

Among the 239 biomineralization TFs in GRNs regulating shell formation of *C. nippona* (Fig. 6a and Supplementary Fig. 12), 36 one-to-one orthologs of these TFs were also supported by published experimental data for shell formation in other molluscs (Supplementary Data 4 and 15). The 36 TFs were further compared with those known to regulate the formation of other biomineralized or hard structures in lophotrochozoans (Fig. 7a and Supplementary Data 15). Several orthologs of these TFs contribute to the formation of other molluscan biomineralized products, including gastropod radulae and scales, as well as chiton spicules (Supplementary Data 15). Moreover, 14 orthologous TFs reported to date from other lophotrochozoan exoskeletons, such as brachiopod shells and chaetae, and annelid tubes and chaetae, were found to be involved in oyster shell formation, with nine of them for both larval and adult shell formation in *C. nippona* (Supplementary Data 15). The widespread recruitment of orthologous TFs across diverse biomineralized exoskeletons and other hard structures suggests the potential existence of a conserved biomineralization toolkit derived from the ancestral lophotrochozoan genome^45–47^.

**Fig. 7.**
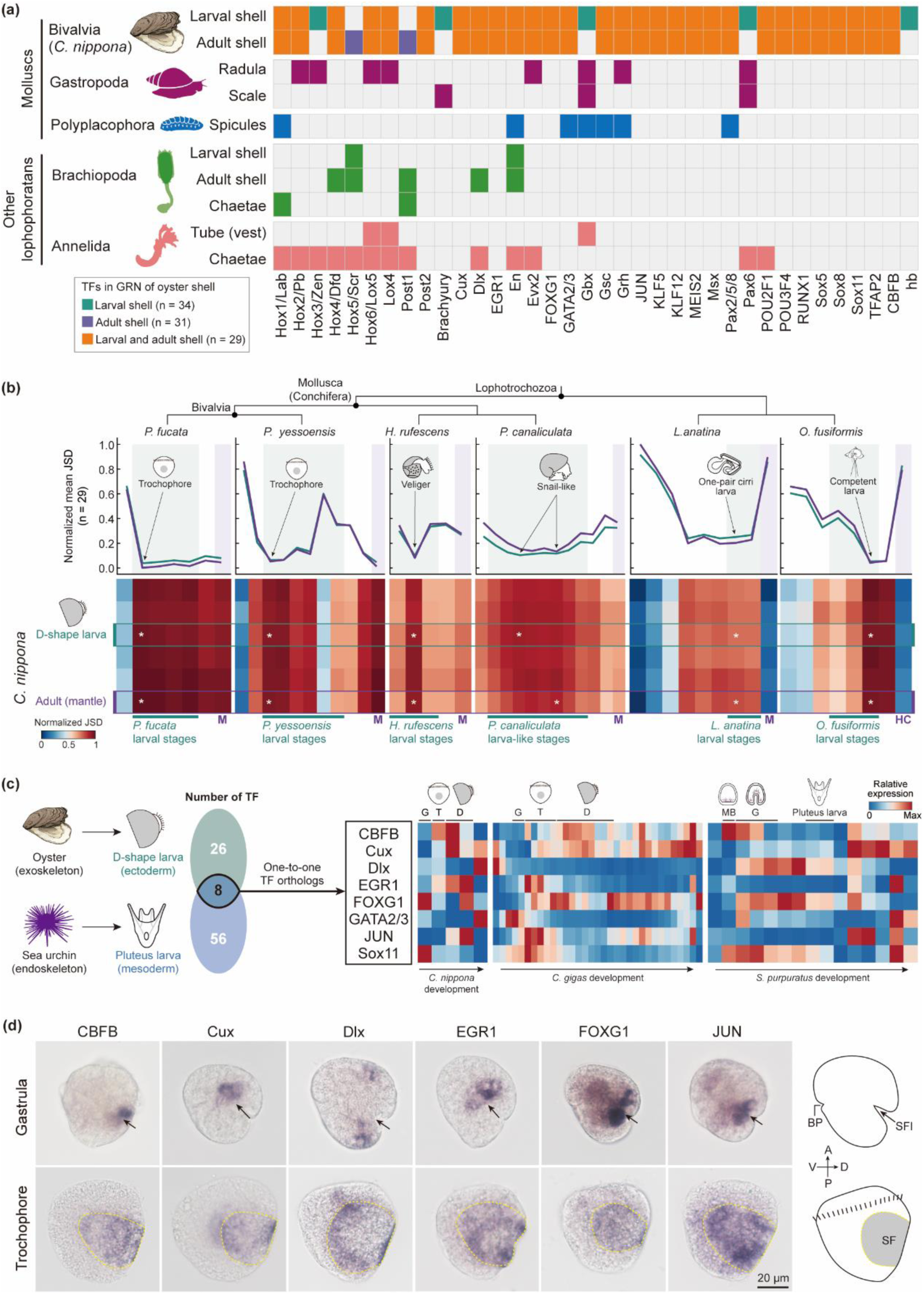
A biomineralization toolkit of TFs in lophotrochozoan larvae, with co-option for endoskeleton development in echinoderms. **(a)** TFs involved in biomineralized structure formation across major lophotrochozoan lineages. Colored squares indicate the recruitment of a TF into the GRN for each structure; while grey squares denote absence. TFs associated with only larval shell formation (green), only adult shell formation (purple), or both stages (orange) in Bivalvia (*C. nippona*) are compared to those involved in other molluscan structures, including the radula and scale in gastropods (dark magenta) and the spicules in polyplacophorans (blue), as well as exoskeletons in other lophotrochozoans, namely the larval and adult shells and chaetae of brachiopods (medium green) and the tube (vest) and chaetae of annelids (salmon). Source data are detailed in Supplementary Data 13. **(b)** Heatmaps of pairwise normalized JSD values of 29 orthologous TFs between *C. nippona* and other lophotrochozoans (*P. fucata*, *P. yessoensis*, *H. rufescens*, *P. canaliculate*, *L. anatina*, and *O. fusiformis*) across life stages. Lower values indicate higher expression similarity. Asterisks mark stages of minimal divergence of each species to *C. nippona* D-shape larvae (green) and adult mantle (purple); corresponding average relative JSDs are shown above each heatmap. Fully labelled heatmaps are shown in Supplementary Fig. 16a. Abbreviations: M adult mantle; HC head and chaetae. **(c)** Comparison of TF orthologs (left) between the exoskeleton GRN of D-shape oyster larvae and endoskeleton GRN of pluteus-stage sea urchin larvae. Right: developmental expression patterns of these TFs in *C. nippona* (from gastrula to D-shape larvae and juvenile), *C. gigas* (from rotary movement to juvenile), and *S. purpuratus* (from cleavage to juvenile). **(d)** Spatial expression patterns of six conserved biomineralization-related TFs in the gastrula and trochophore of *C. nippona*. Whole-mount ISH results (lateral view) showed expression covering the ectodermal shell field invagination (SFI) during gastrulation and the shell field (SF) in trochophore. Arrows indicate the SFI. Yellow dashed lines indicate the SF region in trochophore. Right: schematic diagrams of gastrula (top) and trochophore (bottom) in lateral view. Abbreviations: A anterior; P posterior; V ventral; D dorsal; BP blastopore. Dorsal views in Supplementary Fig. 23.

Next, we investigated whether orthologs for the 29 TFs commonly involved in both larval and adult shell formation in *C. nippona* also contribute to the formation of biomineralized exoskeletons across the life cycle of other lophotrochozoans (Fig. 7a). Orthologs of these TFs were identified in seven molluscs and three other lophotrochozoans with representative biomineralized exoskeletons and available developmental or tissue-specific transcriptome data, as well as in the ecdysozoan *Amphibalanus amphitrite* as an outgroup (Supplementary Data 16). Indeed, the overall expression dynamics of these 29 genes were largely consistent across lophotrochozoans, with high expression levels at early exoskeleton-forming larval stages (trochophore or D-shape larva in bivalves, veliger in gastropods, one-pair cirri larva in the brachiopod *Lingula anatine*, and mitraria larva in the annelid *Owenia fusiformis*) and adult skeletogenic tissues (Supplementary Fig. 19a, b). In contrast, these genes exhibited comparatively lower expression in the tube-forming organs (collar and opisthosoma) of the tubeworm *Paraescarpia echinospica*, consistent with their low expression in the mantle of *A. amphitrite*. To obtain a comparative view of transcriptional dynamics of these TFs in lophotrochozoans, we calculated the pairwise Jensen-Shannon divergence (JSD) index of gene expression similarity between *C. nippona* and other lophotrochozoans, respectively (Fig. 7b, Supplementary Fig. 20 and 21). The early shell-forming larval stages of other lophotrochozoans showed the highest similarity with D-shape larval stage *C. nippona*, thus appearing as a mid-developmental transition between two phases of higher transcriptomic dissimilarity, namely the embryonic stage and the later larval stage (Fig. 7b, Supplementary Fig. 20). These results further support the hypothesis of a shared TF toolkit present in the early larval development of lophotrochozoans, regulating the formation of the biomineralized exoskeleton. Notably, gene expression divergence decreased after metamorphosis of molluscs, particularly in bivalves, but increased in adult skeletogenic organs of non-molluscan lophotrochozoans. Additionally, the adult mantle of *C. nippona* showed higher expression similarity with those of other molluscs, compared to the skeletogenic organs of non-molluscan lophotrochozoans (Supplementary Fig. 21). These findings suggest that the temporal co-option of the 29 TFs into larval and adult shell formation probably represents an evolutionary innovation in molluscs.

### Independent co-option of biomineralization TFs driving the convergence of exoskeletons and endoskeletons in larvae

To assess whether the transcriptional dynamics of biomineralization TFs found in biomineralized exoskeleton of lophotrochozoan larvae are also present in endoskeletons of other bilaterian larvae, we reconstructed the skeletogenic GRN of early pluteus larvae in the sea urchin *Strongylocentrotus purpuratus* using previously published datasets^48, 49^ (Supplementary Fig. 22a, Supplementary Data 17 and 18). 64 biomineralization TFs were identified in pluteus larvae (Supplementary Data 19). Transcriptomic analysis across developmental stages in *S. purpuratus* revealed that these genes are highly expressed during early skeletogenic phases, particularly at gastrula stages, and in adult spines (Supplementary Fig. 22b), suggesting their potential roles in skeletal development and biomineralization. However, the expression of these genes showed no significant differences among adult tissues (Supplementary Fig. 22c), implying that these TFs may play more critical roles during early development of the endoskeleton rather than in adult skeletal maintenance. Moreover, a comparative analysis of transcriptomic similarity throughout the entire development of *S. purpuratus* and another sea urchin species, *Lytechinus variegatus*, demonstrated that the gastrula stage exhibits the highest transcriptional similarity in the life cycles of sea urchins, followed by the pluteus larva stage (Supplementary Fig. 22d and Supplementary Data 19). Similarly, when comparing *S. purpuratus* with the sea cucumber *Apostichopus japonicus*, another echinoderm that forms biomineralized spicules but lacks a substantial endoskeleton during larval development^50^, the blastula is the period of maximal transcriptomic similarity (Supplementary Fig. 22e), corresponding to a skeletogenic mesenchyme in sea cucumbers^51^. These findings suggest that larval skeletogenic mesenchyme formation in echinoderms may be regulated by evolutionarily conserved biomineralization TFs, possibly through heterochronic activation of ancestral adult skeletogenic programs^15, 52^.

We further extended our comparisons of biomineralization TFs between D-shape larvae of the oyster *C. nippona* and pluteus larvae of the sea urchin *S. purpuratus*, and identified only eight orthologous TFs shared between the two species (Fig. 7c). Although these genes are associated with early skeletal development in both organisms, their temporal expression profiles differed markedly. Most of them were highly expressed during trochophore or D-shape larval stages in oysters, whereas they exhibited high expression predominantly in the mesenchyme blastula and gastrula stages of the sea urchin (Fig. 7c). This result indicates a potential heterochronic shift in the deployment of these biomineralization TFs, reflecting lineage-specific timing in the activation of skeletogenic GRNs. In addition to temporal divergence, the embryonic origin of skeletogenic cells also differs fundamentally between molluscan and echinoderm larvae. The shell field forms within the dorsal ectoderm in oyster larvae and later develops into the mantle for shell formation, whereas sea urchin spicules are produced by mesenchymal skeletogenic cells^53^. All eight TFs exhibited spatial expression patterns that covered the shell field of trochophore in oysters, as supported by previous studies^54, 55^ and our ISH results (Fig. 7d, Supplementary Fig. 23). These developmental differences underscore that, despite limited conservation of regulatory TFs, the complexity and morphogenetic mechanisms of skeletal formation have diverged substantially between molluscs and echinoderms. Overall, the convergent evolution of larval exoskeletons in molluscs and larval endoskeletons in echinoderms likely reflects the independent co-option of a restricted set of ancestral TFs into distinct embryonic and developmental contexts.

## Discussion

The evolutionary origin of biomineralized exoskeletons in lophotrochozoans is a topic of ongoing debate^4, 25, 56–61^. Current hypotheses assumed that the common ancestor of lophotrochozoans possessed non-mineralized structures that were independently biomineralized in each lineage^4, 56–59^, while some have argued that the biomineralized exoskeleton is an ancestral lophotrochozoan trait^60, 61^. The construction of GRNs offers a powerful framework to uncover the evolutionary history and hierarchical logic underlying this complex biological process, whereby upstream regulators coordinate the activation of intermediate modules and downstream effectors. However, to date, systematic molecular characterization of biomineralization GRNs in lophotrochozoans remains scarce, with existing studies mostly focusing on the spatial expression patterns of TFs and effector genes^56, 57, 60^. In this study, our comprehensive profiling of multi-stage transcriptomes and epigenomes in *C. nippona* provided a novel perspective on the establishment of shell formation GRNs in molluscs and revealed a biphasic regulatory program orchestrated by a set of conserved TFs that govern distinct processes of molluscan shell formation across life stages. Surprisingly, comparative analyses of transcriptional dynamics suggested a high overall similarity in larval regulatory modules during lophotrochozoan skeletogenesis (Fig. 7a, b), reflecting an ancient and conserved regulatory toolkit for larval exoskeleton formation in lophotrochozoans (Fig. 7). Although the exact timing remains uncertain, this conservation indicated that the GRNs of larval exoskeletons probably originated prior to the divergence of molluscs, brachiopods and annelids, even tracing back to the common ancestor of lophotrochozoans. However, the adult molluscan shell appears to represent a phylum-specific evolutionary innovation. The rapid evolution of downstream effector genes and lineage-specific functional divergence of paralogs can be proposed to have supplied the genetic foundation underlying the remarkable diversity of adult shells, facilitated by dynamic chromatin remodeling and heterochronic deployment of largely conserved upstream TFs shared with the larval regulatory program (Fig. 7). This model is not only consistent with previous studies showing the extensive incorporation of novel or evolutionarily young genes into the shell gland cell and mantle transcriptomes of molluscs^31, 62^, but further highlights the pivotal roles of epigenetic reprogramming in modulating GRNs underlying shell biomineralization across life stages^32^.

Regulatory genes underlying bilaterian developmental programs are evolutionarily ancient, predating the origin of animals, and constitute a conserved developmental toolkit that likely facilitated the rapid diversification of metazoans during the Cambrian^63^. Emerging evidences, including our findings, suggest that a subset of deeply conserved TFs, originally involved in early embryonic patterning, have been co-opted into the biomineralization toolkit of lophotrochozoans^26, 45–47, 64^. When extending comparisons to deuterostomes, we found eight developmental TFs independently co-opted into the early skeletogenic GRNs in both oyster and sea urchin larvae (Fig. 7c, d). Echinoderm endoskeletons have been considered as a result of reutilization of developmental networks deriving from a single evolutionary origin^4^. However, larval skeletogenic cells likely evolved from the heterochronic activation of adult skeletogenic programs^15^. Although further functional studies are required to test these hypotheses, this is strikingly similar to our proposed model for the evolutionary scenario of adult shell in molluscs, where the heterochronic deployment of larval biomineralization TFs activates the adult shell GRN (Fig. 8). Likewise, our approach also has limitations, as TF binding motif associations were inferred through sequence homology-based transfer from the JASPAR database, specifically built from model metazoan species. The lenient matching strategy we used may limit the accuracy of one-to-one TF-target predictions, such as those identified through ChIP-seq or DAP-seq analyses^65, 66^. In addition, bulk-level epigenomic profiling, particularly in mixed larval samples, may capture averaged and noisy signals across heterogeneous cell populations and individuals. The co-occurrence of H3K27ac and H3K27me3 at some genomic loci likely reflects cellular heterogeneity rather than bivalency within single cells, as these modifications cannot coexist on the same histone tail (Fig 4a). Given the potential cellular heterogeneity within larval and adult biomineralizing tissues, comparative single-cell multi-omic data from molluscs and other biomineralizing lophotrochozoans in future work will be critical for resolving cell-type-specific contributions to the GRNs and transcriptomic similarities we observed. Despite these limitations, our integrative analyses provide valuable insights into the evolutionary strategies of biomineralization GRNs across bilaterians.

**Fig. 8.**
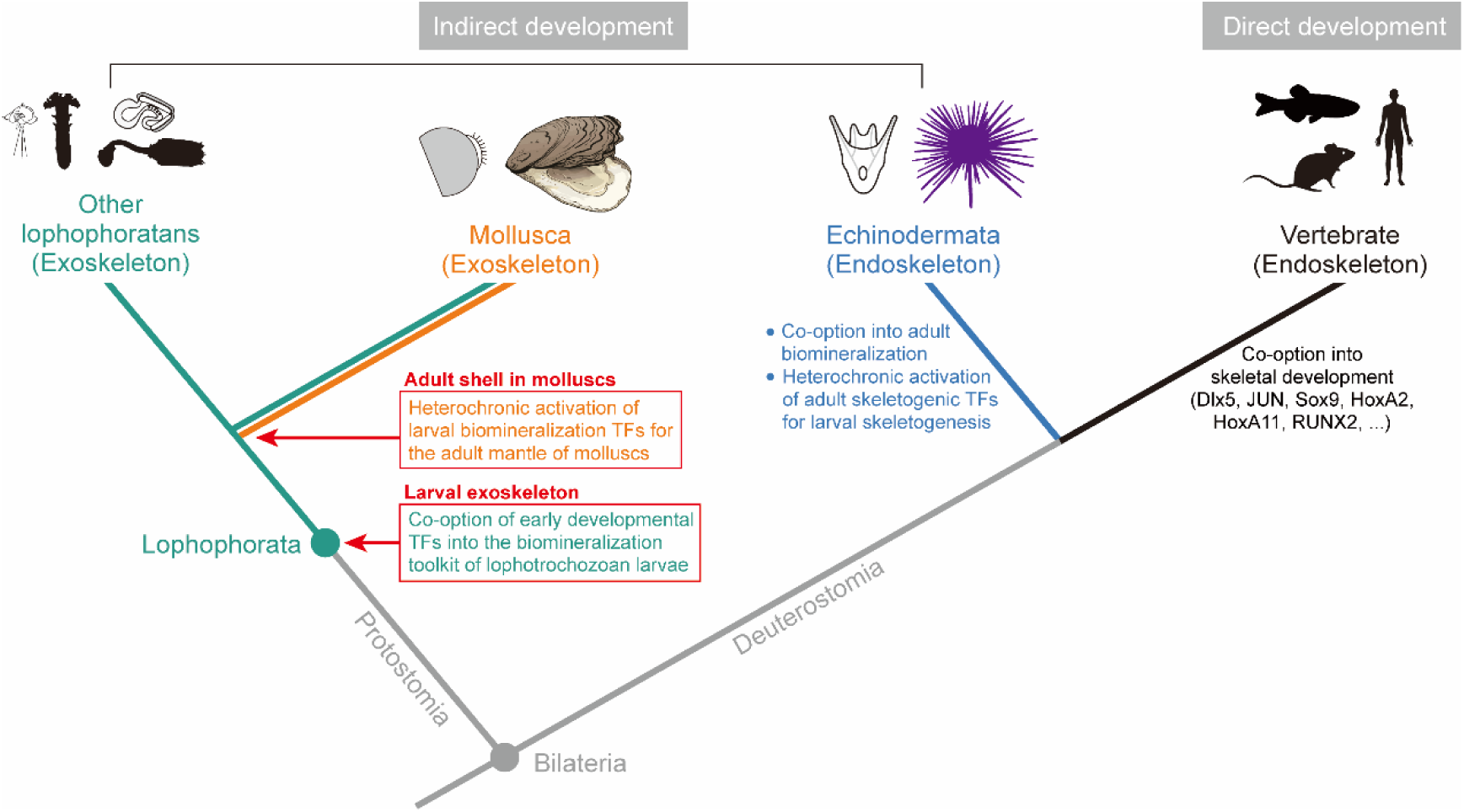
Evolutionary scenario of TF co-option for biomineralized skeletons across bilaterian life cycles. The schematic phylogeny illustrates the convergent and heterochronic co-option of TFs into biomineralization GRNs in major bilaterian clades. In the common ancestor of lophotrochozoans, early developmental TFs were co-opted into larval biomineralization GRNs, establishing a conserved exoskeleton toolkit. In molluscs, these TFs were heterochronically activated in the adult mantle to regulate shell formation. By contrast, adult skeletogenic TFs were repurposed during early development to form the larval endoskeleton in echinoderms, while contributing to adult biomineralization^15^. Vertebrates, which typically undergo direct development, exhibit recurrent co-option of developmental TFs (e.g., *Dlx5*, *JUN*, *Sox9*, *HoxA2*, *HoxA11*, *RUNX2*) into skeletal development^13, 67, 68^. These parallel and lineage-specific events highlight the independent evolutionary origins of biomineralized skeletons across Bilateria through the repeated redeployment of ancient regulatory modules.

Notably, many biomineralization TFs in lophotrochozoans and echinoderms belong to gene families also essential for vertebrate skeletogenesis, such as *Dlx5*, *JUN*, *Sox9*, *HoxA2*, *HoxA11*, and *RUNX2*^13, 67, 68^. Although these TFs are not one-to-one orthologs, their recurrent recruitment into biomineralization programs across diverse bilaterian lineages implies the inherent regulatory potential of these TF families for controlling skeletal development (Fig. 8). This widespread convergence likely reflects a deep evolutionary constraint on upstream regulatory modules, wherein components of ancient developmental toolkits are repeatedly co-opted for similar functional demands during the independent evolution of biomineralized skeletons in bilaterians^4^. Beyond mineral deposition, such regulatory conservation may extend to broad physiological roles of skeletal systems. A recent study has revealed that molluscan shells harbor a bone marrow-like hematopoietic stem cell niche, where stem cells contribute to both shell regeneration and systemic blood cell supply, under the regulation of deeply conserved TFs also essential for vertebrate skeletogenesis^69^. Thus, the molluscan shell highlights how conserved developmental programs can be redeployed through regulatory co-option to generate lineage-specific innovations with ancient physiological roles, such as hematopoiesis, thereby transforming a simple biomineralized secretion into a complex and life-sustaining organ.

In summary, our study provides a novel (epi)genomic and evolutionary framework to understand molluscan shell formation. We highlight that dynamic chromatin landscapes, coupled with conserved regulatory modules co-opted from developmental programs, shape the diversity of biomineralized exoskeletons across life stages. Regardless of scenario, our data generates novel hypotheses and uncovers a hierarchical decoupling of evolutionary conservation and innovation of biomineralized skeletons, which may represent a general pathway contributing to biological complexity in lophotrochozoans and other bilaterian clades.

## Methods

### Animal culture, sample collection and shell morphology

*C. nippona* adults were cultured in an oyster farm of Rushan, China. Embryos and larvae were obtained through artificial fertilization and reared at 26-28 ℃. Over 10,000 embryos/larvae were sampled in duplicates at key shell-forming stages: gastrula (8 hours post fertilization, hpf), trochophore (14 hpf), D-shaped larvae (22 hpf), and later D-shaped larvae (3 days post fertilization, dpf), then flash-frozen in liquid nitrogen and stored at -80 °C. As embryonic development is not perfectly synchronous, the 8 hpf samples likely includes embryos transitioning from morula to early gastrula stages. For scanning electron microscopy (SEM), larvae were fixed in 2.5% glutaraldehyde at 4 °C overnight and processed as described previously^70^.

Whole soft bodies of juvenile oysters (50-day-old, n = 5 per replicate) were pooled, immediately flash-frozen in liquid nitrogen, and stored at -80 °C. To stimulate shell formation, two-year-old adult oysters were shell-damaged by drilling and cultured as previously described^36^. Once shell repair was completed, mantle tissues surrounding the repaired sites were collected, flash-frozen in liquid nitrogen, and preserved at - 80 °C. Repaired shells were carefully separated from the original shells, processed, and imaged under SEM as previously described^36^.

### Iso-seq and bulk RNA-seq

Total RNA was extracted from samples of all six stages/tissues described above, using TRIzol reagent (Invitrogen, USA). RNA quality assessment was performed using a NanoDrop spectrophotometer and Agilent 5400 Fragment Analyzer system. To obtain a reliable transcript annotation, an Iso-seq library was constructed by pooling equal amounts of RNA from all samples, and sequenced on a PacBio Sequel II SMRT cell, generating a total of 64.2 Gb long reads. In addition, short-read mRNA sequencing was performed on three biological replicates per stage/tissue (Supplementary Data 1), using the Illumina NovaSeq 6000 platform (PE150), generating ∼ 6 Gb of data per replicate.

### Improved genome annotation of *C. nippona*

Gene prediction was performed using Iso-seq and bulk RNA-seq data generated in this and previous studies^36, 71^, following an established pipeline with modifications^36^. Briefly, short RNA-seq reads were aligned to the *C. nippona* genome using HISAT2 (v.2.2.1) ^72^ and assembled with Trinity (v.2.15.1) ^73^, then integrated with evidence derived from protein homology and de novo predictions from Augustus (v.3.4.0) ^74^ with the MAKER pipeline (v.3.01.03) ^75^. Iso-seq subreads were processed and aligned to the genome assembly with SMRT Link software (v12.0) (https://www.pacb.com/support/software-downloads/). The TAMA package^76^ was used to collapse redundant transcripts and merge full-length transcriptomes. Final gene models were obtained by combining MAKER and Iso-seq annotations using AGAT (v.1.3.0) (https://github.com/NBISweden/AGAT). Completeness was assessed with BUSCO (v.5.4.7) ^77^, and functional annotation was performed using DIAMOND (v2.1.10.164) ^78^ against NCBI non-redundant (NR), Uniprot/SwissProt, eggNOG (v.5.0) ^79^, GO categories, and Kyoto Encyclopedia of Genes and Genomes (KEGG) databases, with domain prediction using InterProScan (v.5.64-96.0) ^80^.

### Time-series RNA-seq analysis

Raw RNA-seq reads were processed with fastp (v.0.23.4) ^81^, and gene expression (Transcripts per million, TPM) was quantified using salmon (v.1.10.1) ^82^ with predicted transcript sequences as reference. Trimmed mean of M-values (TMM) normalization was applied by a Trinity utilities script. Differentially expressed genes were determined using edgeR (v.3.40.2) ^83^ (FDR < 0.05, |log_2_foldchange| > 1).

For clustering analyses, normalized TPM values of biological replicates were averaged to obtain a single gene expression value per gene. *K*-means cluster analysis of expressed genes (normalized TPM > 1 in at least one sample) was performed with *Z*-scaled TPM using the mfuzz package (v.2.62) ^84^. Heatmaps were generated with the ComplexHeatmap package (v.2.18.0) ^85^. GO enrichment for each gene cluster was analyzed using the clusterProfiler package (v.4.10.0) ^86^, against all expressed genes annotated by the eggnog (v.5.0) database^79^, with a significance cutoff of Q-value < 0.05.

### Transcription factor (TF) characterization and analysis

We used the AGAT toolkit to generate a *C. nippona* non-redundant proteome with only the longest isoform per gene. Genome-wide identification of TFs was performed by combining homology-based searches against AnimalTFDB 4.0 database^87^ using BLASTp (e-value < 1e-10), and de novo domain-based prediction using HMMER (v3.4) and InterPro HMM profiles^88^, with the parameter “cut_tc” or a cut-off of 0.0001 for those without trusted cutoffs. The final TF repertoire was defined as the union of results from both methods.

### Shell matrix protein (SMP) extraction

Larval SMPs were extracted from *C. nippona* D-shape larvae (22 hpf) following a modified protocol^27^. Five mL of larvae were washed and then centrifuged (1,000 g, 3 min, 4 °C) to remove the supernatant. The remaining shells were immersed in 5% NaOCl for 15 min under 4 °C with gentle shaking, washed with Milli-Q water three times, and then cleaned with a 1-min ultrasound treatment in the Milli-Q water to ensure thorough cleaning (Supplementary Fig. 24). Cleaned larval shells were filtered using a nylon mesh and decalcified in 1 M acetic acid at 4 °C for 12 h. Acid-soluble matrix (ASM) and acid-insoluble matrix (AIM) fractions were separated by centrifugation (4,000 g, 30 min, 4 °C), and the AIM fractions was further washed. Both fractions were mixed and concentrated using an Ultracel-10 centrifugal filter unit.

Adult shells (including repaired ones) of three *C. nippona* individuals were treated with 5% NaOCl for 12 h at RT, then washed, fragmented, ultrasonically cleaned for 2 min, and air-dried. After grinding, the shell powder (∼ 40 g) was decalcified in 1 M acetic acid at 4 °C overnight. AIM and ASM were isolated as described above.

### LC-MS/MS analysis and protein identification

AIM and ASM proteins from larvae and adult shells were respectively treated with SDT-lysis buffer (4% SDS, 100 mM DDT, 100 mM Tris-HCl, pH 7.6) in a boiling water bath for 10 min. After cooling to RT, the supernatant was collected by a short centrifugation and mixed with UA buffer (8 M Urea, 150 mM Tris-HCl, pH 8.0). Sample preparation and LC-MS/MS analysis were performed following a previous protocol^36^. In brief, the mixtures were processed by ultrafiltration, alkylation, enzymatic digestion, and peptide desalting, and then analyzed using a Q-Exactive Plus mass spectrometer coupled with an EASY-nLC 1200 system (Thermo Fisher Scientific).

MS/MS spectra from adult samples and previous datasets^36^ were searched against the *C. nippona* predicted proteome using MaxQuant (v.2.0.10) ^89^ (FDR < 0.01, unique peptides ≥ 2). For larval SMPs, raw spectra were also analyzed as described above similarly. To comprehensively identify the SMPs of the oyster larvae, we ran BLASTp best-reciprocal hits between the *C. nippona* proteome and a larval shell proteome of a closely related oyster species *C. gigas*^27^ (e-value < 1e-20, sequence identity > 80%) to retrieve one-to-one orthologues. All proteins identified by both methods were considered as larval SMPs in *C. nippona*. Finally, 101 larval and 289 adult SMPs were identified in *C. nippona* (Supplementary Data 20). Signal peptides of proteins were predicted with SignalP-6.0^90^.

### Biomineralization gene characterization and analysis

Biomineralization TFs in *C. nippona* were identified by BLASTp best-reciprocal hits (e-value < 1e-10) between TFs expressed in *C. gigas* trochophore shell field cells^62^ and the *C. nippona* TF repertoire, as well as from known molluscan shell-related TFs based on previous ISH data (Supplementary Data 3). Biomineralization TFs in *S. purpuratus* pluteus larvae were defined as those expressed in skeleton-related cell clusters from published single-cell transcriptome data^49^.

In addition to SMPs, we identified other candidate biomineralization effector genes involved in shell formation, including biomineralization enzymes and transmembrane ion transporters^19, 23, 25^, by searching the *C. nippona* proteome against InterPro HMM profiles^88^ using HMMER (v3.4) with the “cut_tc” parameter (Supplementary Data 21). RNA-seq data of adductor muscle, gills, digestive gland, and hemolymph from our previous study^36^ were also processed following the same pipeline described above and were normalized together with newly generated RNA-seq data. Gene co-expression analysis was performed using the WGCNA package^42^ across multiple developmental stages and adult tissues, and identified 11 gene modules (MEs) significantly correlated with shell-forming stages and mantle tissue (Supplementary Fig. 25). Effectors within these modules were considered involved in *C. nippona* shell formation.

Biomineralization genes in other mollusks (*C. gigas*, *Pinctada fucata*, *Patinopecten yessoensis*, *Haliotis rufescens*, *Pomacea canaliculata*) were identified using similar approaches. SMP data for *P. fucata* larvae, *P. canaliculata* adult, and *C. gigas* larvae and adult were obtained from previous studies^27, 91^. MS/MS datasets for *P. fucata* adult shells (PXD006786 and PXD010130) were downloaded from the PRoteomics IDEntifications Database (PRIDE), and protein identification was performed using MaxQuant (v.2.0.10) ^89^. For species lacking SMP data, only biomineralization enzymes and transmembrane ion transporters were considered. WGCNA analysis was performed to further identify candidate biomineralization effector genes in these molluscs, which were highly expressed in larval or adult shell-related modules (Supplementary Fig. 26).

### ATAC-seq and CUT&Tag experiments

We performed two replicates of ATAC-seq on five developmental stages and adult mantle tissue used for RNA-seq of *C. nippona* using ∼ 50,000 cells per sample, following the Omni-ATAC protocol^92^. After nuclei isolation and tagmentation at 37 °C for 30 min, libraries were purified, PCR-amplified (12-cycle) (TruePrep® DNA Library Prep Kit V2 for Illumina; primers from TruePrep® Index Kit V2 for Illumina), and then size-selected (0.55× and 1.5× ratios) using VAHTSTM DNA Clean Beads as indicated by the supplier. Final libraries were quantified (> 10 ng/μL) using a Qubit 4 fluorometer and quality checked for fragment distribution on a Qsep-400 system (Bioptic), ensuring a clear ladder pattern with a major peak around 200 bp, no smearing or adapter contamination. Sequencing was performed on an Illumina NovaSeq 6000 platform in PE150 mode.

CUT&Tag assays were conducted with two replicates of D-shaped larva and adult mantle samples (∼ 100,000 nuclei per sample) used for RNA-seq and ATAC-seq of *C. nippona*, following a previous protocol with minor modifications^93^. Nuclei were bound to activated concanavalin A-coated magnetic beads, resuspended in Dig-Wash Buffer (0.05% digitonin and 2 mM EDTA). Primary antibodies, including H3K4me3 (Active motif, #61379), H3K27ac (Active motif, #39133) and H3K27me3 (Abcam, #AB6002), were added at a 1:50 dilution and incubated at room temperature for 2 h, followed by a 1:50 dilution of secondary antibody (IgG, Proteintech, #B900210) for 1 h. After washing, nuclei were treated with pG-Tn5 adapter complex (1:200 in Dig-300 Buffer) for 1 h, followed by tagmentation in 10 mM MgCl₂ at 37 ℃ for 1 h. The reaction was terminated with the addition of 0.5 M EDTA, 10% SDS, and Proteinase K, and samples were incubated at 55 °C for 1 h. DNA was then purified using phenol-chloroform and ethanol precipitation. Libraries were PCR-amplified (16 cycles), purified with AMPure XP beads (Beckman Coulter), and sequenced on an Illumina NovaSeq 6000 system (PE150).

### ATAC-seq and CUT&Tag data analysis

Raw reads were quality filtered with fastp (v.0.23.4) ^81^ and mapped to the *C. nippona* genome using Bowtie2 (v.2.5.2) ^94^. PCR duplicates, unmapped reads, multi-mapped reads, low-quality reads (MAPQ < 20), and reads mapped to the mitochondrial genome were removed using alignmentSieve tool from deepTools (v.3.5.4) ^95^. For ATAC-seq data, the mapped reads were shifted +4/-5bp depending on the strand of the reads to reflect Tn5 cleavage sites. Peak calling was performed using MACS3 (v.3.0.0b3) ^96^ (ATAC-seq: --nomodel --shift -75 --extsize 150 --qvalue 0.05; H3K4me3 and H3K27ac: --qvalue 0.05; H3K27me3: --broad --broad-cutoff 0.05). Reproducible peaks across biological replicates were identified using the irreproducible discovery rate method (IDR < 0.05) (v.2.0.4). For data visualization, merged bam files of two biological replicates were converted to RPKM (Reads Per Kilobase per Million mapped reads) normalized bigwig files with 10 bp bin size using deepTools (v.3.5.4)^95^. Peak tracks and gene structures were visualized with pyGenomeTracks (v.3.9) ^97^. Consensus peaks were generated with DiffBind (v.3.12.0) ^98^ and annotated to genomic regions using UROPA^99^. The whole genome was divided into four regions: promoter (from -2,000 bp to +500 bp of TSS), genic (from +500 bp of TSS to TES), distal (other region within 100 Kb of the nearest gene) and no annotation (> 100 Kb away from the nearest gene). Reads count were quantified using featureCounts^100^, and TMM-normalized by the Trinity utilities script. Differential peaks were identified using edgeR (v.3.40.2) ^83^ (FDR < 0.05, |log_2_foldchange| > 0).

### Chromatin state analysis

We used ChromHMM (v.1.26) ^33^ with the default parameter to train the chromatin state prediction model by integrating CUT&Tag (H3K4me3, H3K27ac and H3K27me3) and ATAC-seq data merged from two biological replicates of D-shape larvae and adult mantles. Models ranging from 4 to 16 states were built, and an eight-state model on strong correlations among states, ensuring a balance between depth and clarity in the results (Supplementary Fig. 7a). Binarization was performed at a resolution of 200 bp. The annotation of each state was achieved by the “OverlapEnrichment” and “NeighborhoodEnrichment” functions provided by ChromHMM.

### GRNs construction and analyses

GRNs for larval and adult shell formation were constructed by integrating established methods^101–103^ with slight modifications (Supplementary Fig. 10a). Genes with TPM > 1 and TF binding motifs in active promoter or genic regions were used to build *cis*-regulation networks. Motif information was obtained from the JASPRA non-redundant CORE metazoan database, and *C. nippona* TF motifs were inferred by reciprocal BLASTp best-hits (e-value < 1e-10, identity > 30%) against JASPAR TFs (Supplementary Data 11). For ANANSE^41^, ATAC-seq IDR peaks with at least 50% overlap with poised or active regions were extracted and re-centered their coordinates on the summits of chromatin accessibility peaks. ANANSE binding was run with ATAC-seq, CUT&Tag (H3K27ac) BAM files and the summit-centered peaks to generate genome-wide TF binding profiles for D-shape larvae and adult mantles, respectively. ANANSE network was then used to construct GRNs respectively for larvae and adults based on the predicted TF binding profiles and RNA-seq data. The top 25% of regulatory edges (probability score > 75th quantile) were retained as the final ANANSE networks. For TF footprinting analysis, ATAC-seq BAM files of two biological replicates were merged using samtools (v.1.13) ^104^ and analyzed with TOBIAS (v.0.16.1) ^105^ to identify bound TF binding sites (footprint score above BINDetect threshold). Footprint-based networks were constructed for larval (trochophore and D-shaped stages) and adult (mantle tissue) phases. Final GRNs were generated by combining ANANSE and footprint networks, defining biomineralization genes and TFs as targets. Networks were visualized with Cytoscape (v.3.10.1) (https://cytoscape.org/), and TF centrality (outdegree index) was calculated using the igraph R package. The same pipeline was applied to establish GRNs for skeleton formation in *S. purpuratus* larvae, using ATAC-seq peaks and gene expression data, with positive marker genes from skeleton-related clusters in the published single-cell transcriptome^49^ as targets (Supplementary Fig. 19a).

### Phylostratigraphy analysis

Gene age was estimated for both species using the GenERA tool (v.1.4.2) ^40^, which performs genomic phylostratigraphy by searching for homologs in the NCBI NR database (Release 2024_01_01) and predict gene origins based on NCBI Taxonomy. Biomineralization effector genes were classified as pre-Mollusca if they originated at or before the Cellular organisms-Lophotrochozoa ancestor, while genes originating from between Mollusca and Mollusca-specific families/species were considered Mollusca-family/species origin genes (Supplementary Fig. 9a and Supplementary Data 22). Paralog identification and dating in *C. nippona* and five other molluscs followed a previously described protocol^106^. Paralogs and their duplication timing across 40 metazoan species (Supplementary Data 23) were inferred using OrthoFinder (v2.5.2) ^107^. The phylostratum of each duplication node was classified as described above (Supplementary Fig. 9a).

### Comparative analyses using TFs

We identified putative one-to-one orthologs of 29 biomineralization TFs between *C. nippona* and ten other lophotrochozoans, as well as *A. amphitrite*, by performing reciprocal best-hit BLASTp searches with an e-value threshold of 1e-3 (Supplementary Data 15). The results were consistent with those obtained using OrthoFinder (v2.5.2) ^107^. Publicly available RNA-seq datasets of developmental time courses and adult tissues for these species were downloaded from the NCBI SRA database (Supplementary Data 24) and processed using the same pipeline described above for *C. nippona*. Following established methods from a previous study^108^, we then performed a quantile transformation to TPM values and calculated the JSD comparing the expression profiles of the 29 single-copy orthologous TFs between each species and *C. nippona*. To ensure statistical robustness, 1,000 bootstrap replicates were performed to estimate mean JSD values. Raw mean JSD values were adjusted by the number of orthologs and normalized using the global minimum and maximum of all adjusted values across comparisons. Relative JSD values were similarly normalized within each pairwise comparison. Transcriptomic divergence (JSD values) of biomineralization TFs between *S. purpuratus* and two other echinoderms was calculated using the same method (Supplementary Data 19 and 24), yielding mean and standard deviation values.

### ISH experiments

Gene-specific primers containing SP6/T7 adaptor sequences were designed using Primer premier (v.6.0) (Premier Biosoft International, Palo Alto, CA) to amplify cDNA fragments for each gene (Supplementary Data 25). Digoxigenin-labelled RNA probes (antisense and sense) were synthesized using purified PCR products and DIG RNA labeling Kit (SP6/T7) (Roche, Switzerland). The synthesized probes were purified using the MEGAclear™ Transcription Clean-Up Kit (Thermo Fisher Scientific, USA) and stored at -80 °C until use.

ISH in oyster larvae and adult mantles was performed as previously described^36, 70^. Larvae and adult mantles were fixed in 4% paraformaldehyde (PFA) overnight at 4 °C, dehydrated with 100% methanol, and stored at -30 °C. For ISH of larvae, samples were rehydrated, digested with proteinase K (with D-shape larvae pre-treated with 0.5 M EDTA), post-fixed in 4% PFA, pre-hybridized, and hybridized overnight at 65 °C with 500 ng/mL RNA probe at 65°C. After washing, samples were blocked and then incubated with anti-DIG AP antibody (1:5000) (Roche, Switzerland) overnight at 4°C. Finally, samples were washed and incubated with NBT/BCIP substrate (Roche, Switzerland) for signal detection. For ISH of adult mantles, fixed mantles cleared in xylene, embedded in paraffin, and sectioned at 5 μm. After deparaffinization, rehydration, proteinase K digestion, prehybridization, hybridization (with RNA probes at 1 ng/mL), followed by antibody incubation (1:3000) and NBT/BCIP staining at 4 °C. Negative controls were done for all genes using sense probes, as well as by including all hybridization ingredients except probes. Images were acquired using an Olympus BX53 microscope.

### Luciferase reporter assay

Dual-luciferase reporter assays were performed to evaluate the transcriptional activities of two predicted promoters (peak18565 for *Gigasin2*, and peak151754 for *Unc022929*) following our previously published protocol^109^. Genomic DNA from mantle tissue of three *C. nippona* adults was extracted using the phenol-chloroform method. The promoter sequences and coding sequence of *Hox4* were amplified and cloned into pGL3-basic and pcDNA3.1(+) plasmids, respectively (Supplementary Data 25). HEK293T cells were seeded in 24-well plates (Corning, USA) and cultured in DMEM (Hyclone, USA) with 1% 1×penicillin-streptomycin antibiotics and 10% fetal bovine serum (FBS) (Hyclone, USA) at 37°C with 5% CO_2_. Once cells reached 80% confluence, they were transfected with serum- and antibiotic-free medium using Lipofectamine 3000 (Invitrogen, USA) at a 1:1 ratio of promoter to TF plasmids. DNA-lipofectamine complexes were added dropwise to the medium. After 48 h, luciferase activity was measured using the Dual-Luciferase Reporter Assay System (Promega, USA) according to the manufacturer’s instructions. Firefly and Renilla luciferase activities were quantified using the Synergy™ H1 Multimode Reader (BioTek, USA). Relative luciferase activity was assessed by calculating the ratio of firefly luciferase activity to Renilla luciferase activity. Each targeted fragment was tested in triplicate.

### Knockdown of *Hox4* using RNAi

A shell repair assay was performed on two-year-old *C. nippona* adults (n = 15 per group). A hole (∼2 mm in diameter) was drilled into the left valve of each oyster. Oysters were divided into three groups: *Hox4* siRNA (Hox4-RNAi), negative control siRNA (NC), and DEPC water control. *Hox4*-RNAi and NC were synthesized by GenePharma (Shanghai, China) (Supplementary Data 24). The selected siRNA was optimized in a pilot experiment (Supplementary Fig. 27 and Supplementary Data 25). Based on the results, we selected the most effective siRNA “*Hox4*-437”. The siRNAs were diluted to 1 µg/µL in DEPC-treated water containing phenol red (1:20 dilution) for visual tracking during injection. A small gap was created near the adductor muscle on the right shell using forceps, allowing access for injection. In the experimental group, each oyster was injected with 20 µL of *Hox4*-RNAi solution into the pericardial cavity using a microsyringe. The control groups were treated with an equal volume of NC solution and DEPC water, respectively. Injections were performed every 48 h for a total of three times. During the experiment, oysters were cultured at 18-20 °C and fed daily with *Chlorella* sp. No oyster died during the experiments. Mantle tissues surrounding the shell holes were collected one day after the final injection for mRNA and protein quantification. Newly grown shells were cleaned, photographed, and analyzed using ImageJ (https://imagej.net/ij/) to assess the areas of newly grown shells and calculate shell repair rates (Supplementary Fig. 14).

### Real-time quantitative polymerase chain reaction (RT-qPCR) analysis

Total RNA was extracted from mantle tissues of seven randomly selected individuals from each group in the RNAi experiment using TRIzol reagent (Invitrogen, USA), following the manufacturer’s protocol. Total RNA (1 µg) was reverse transcribed into cDNA using the Evo M-MLV RT Mix Kit (Accurate Biotechnology, China) according to the manufacturer’s instructions. Primers for the *Hox4* gene and internal control genes *EF1α* and *GAPDH* were designed with Primer premier (v.6.0) (Supplementary Data 25). All primer pairs were validated by melting curve analysis. RT-qPCR was performed using the SYBR Green Premix Pro Taq HS qPCR Kit (Accurate Biotechnology, China) on a LightCycler 480 real-time PCR system (Roche, Switzerland). Relative gene expression levels were calculated using the 2^-ΔΔCT^ method^110^.

### Western blot analysis

Mantle tissues from three individuals, also used for RT-qPCR, each from the NC and *Hox4*-RNAi groups were homogenized in PBS and centrifuged to collect the supernatant. Protein concentrations were quantified using a BCA Protein Assay Kit (Beyotime, China). Samples were diluted to equal concentrations with SDS-PAGE loading buffer (Solarbio, China), denatured at 95 °C for 10 min, and 40 µg of protein per sample was separated on a 12% SDS-PAGE gel. The separated proteins were transferred onto 0.4 µm polyvinylidene fluoride (PVDF) membranes (Beyotime, China). Membranes were blocked at room temperature for 2 h with 5% non-fat milk in TBST buffer and then incubated overnight at 4 °C with primary antibodies, including Hox4 (Thermo Fisher Scientific, USA, #H00003221-M02) and β-actin (Beyotime, China, #AF0003). After three washes with TBST, membranes were incubated with HRP-conjugated goat anti-rabbit secondary antibody (1:1000, Beyotime, China) at RT for 2 h. Finally, the blots were measured with SuperPico ECL chemiluminescence kit (Vazyme, China) and visualized using the GE ImageQuant LAS4000mini system (GE, USA). The experiments were repeated twice in independent experiments using the same protocol, and yielded consistent results (Supplementary Fig. 28 and Supplementary Data 26).

### RNA-seq after RNAi experiment

Total RNA from mantle tissues of three individuals for Western blot analysis was used for short-read mRNA sequencing, with three biological replicates. Sequencing was performed using the Illumina NovaSeq X Plus platform (PE150), generating ∼ 6 Gb of data per replicate. The analysis workflow of the RNA-seq data was conducted as described in the previous section.

## Ethics statement

All animal experiment guidelines were approved by the Institutional Animal Care and Use Committee of Ocean University of China (OUC-IACUC), with approval numbers 2020-0032-0517 and 2023-0032-0039.

## Data availability

The raw sequence data have been deposited on NCBI under the BioProject accession number PRJNA1267763. The shell proteomic data are available from ProteomeXchange (accession number: PXD051659). Improved genome annotation of *C. nippona* is accessible on Figshare (https://doi.org/10.6084/m9.figshare.29204657). Source data are provided with this paper.

## Code availability

The scripts and codes for this study can be found on GitHub: https://github.com/StevenBai97/OysterChromatinDynamics.

## Acknowledgements

This work was supported by grants from National Natural Science Foundation of China (W2411019, 32373115), the China Agriculture Research System Project (CARS-49), the Taishan Industrial Experts Program, and Innovation Project from Qingdao Institute of Blue Seed Industry. We acknowledge the support of the High-Performance Biological Supercomputing Center at the Ocean University of China, and the Marine Biodiversity and Evolution Research Institute’s Instrument and Equipment Sharing Platform for providing the high-speed centrifuge (XPN-100) for this research.

## Author contributions

Y.B., S.L., and Q.L. conceived and designed the project. Y.B., Y.H., and S.J. collected the samples. Y.B. and Y.M. performed the experiment and collect the data. Y.B. performed the data analyses. Q.L., H.Y., and L.K. contributed materials and reagents. Y.B. wrote the manuscript with input from S.L., D.M., S.D., and Q.L. All authors reviewed and approved the manuscript.

## Competing interests

The authors declare no competing interests.

**Correspondence** and requests for materials should be addressed to Qi Li.

