## Supplementary notes, figures, and data for "Evolutionary innovation within conserved gene regulatory networks underlying biomineralized skeletons in Bilateria": Description of Additional Supplementary Files.pdf

**Supplementary Data 1.** Statistics of improved gene prediction of the *C. nippona* genome.

**Supplementary Data 2.** Statistics of RNA-seq libraries for *C. nippona*.

**Supplementary Data 3.** GO enrichment analysis of RNA-seq clusters. GO terms mentioned in the main text are highlighted with yellow backgrounds.

**Supplementary Data 4.** Transcription factors involved in molluscan shell formation for which data is available in previous studies.

**Supplementary Data 5.** RNA-seq clustering of biomineralization effector genes and their categories.

**Supplementary Data 6.** Differentially expressed biomineralization effector genes between larval stages and the adult mantle.

**Supplementary Data 7.** Statistics of ATAC-seq experiments of *C. nippona*.

**Supplementary Data 8.** Statistics of histone modification experiments of *C. nippona*.

**Supplementary Data 9.** Differential chromatin accessibility and histone modifications associated with differentially expressed biomineralization effector genes between D-shape larvae and adult mantles.

**Supplementary Data 10.** Specialized biomineralization effector paralogs for larval and adult shells formation in *C. nippona* generated by OrthoFinder with 40 species used for the analysis.

**Supplementary Data 11.** Phylostrata and duplication nodes of biomineralization effector paralogs in five other mollusks.

**Supplementary Data 12.** Association of JASPAR motif to *C. nippona* genes.

**Supplementary Data 13.** Gene regulatory networks (GRNs) of larval and adult shell formation. The interaction probability was predicted by ANANSE. Gene names are based on annotations from the eggNOG database.

**Supplementary Data 14.** Experimental data obtained from the dual-luciferase reporter assay system.

**Supplementary Data 15.** Transcription factors (TFs) involved in larval and adult shell formation in Bivalvia (*C. nippona*) and other hard structures in lophotrochozoans for which data is available. These TFs were identified in GRNs regulating shell formation of *C. nippona* (Fig. 6a), and were also supported by published experimental data (Supplementary Data 3). Reference numbers in parentheses indicate sources of experimental data.

**Supplementary Data 16.** List of single-copy orthologue biomineralization TFs between *C. nippona* and other lophotrochozoans, which were identified as key TFs of both larval and adult shell formation GRNs in *C. nippona*.

**Supplementary Data 17.** Association of JASPAR motif to *S. purpuratus* genes.

**Supplementary Data 18.** Skeletogenic GRN of early pluteus larvae in *S. purpuratus*. The interaction probability was predicted by ANANSE.

**Supplementary Data 19.** List of single-copy orthologue biomineralization TFs between *S. purpuratus* and other echinoderms.

**Supplementary Data 20.** SMPs of larval and adult shell in *C. nippona*.

**Supplementary Data 21.** Gene families and HMM domains of biomineralization effector genes. Due to the absence of an HMM domain, voltage-dependent calcium channels (VDCCs) in *C. nippona* were identified by homology-based search using the BLAST algorithm (e-value < 1e-5) against known VDCC sequences from the NR database.

**Supplementary Data 22.** Phylostratigraphic annotation of biomineralization effector genes in *C. nippona*.

**Supplementary Data 23.** Metazoan genomes used for the identification of paralog pairs by OrthoFinder analysis.

**Supplementary Data 24.** RNA-seq datasets used for comparative transcriptomic analyses.

**Supplementary Data 25.** Primers used in this study. For ISH, Sp6/T7 promoter sequences were underlined. For luciferase reporter assays, lowercase letters indicate added restriction enzyme recognition sites and flanking sequences for cloning.

**Supplementary Data 26.** Grayscale values of the Western Blot, analyzed using ImageJ.
