## Supplementary notes, figures, and data for "Evolutionary innovation within conserved gene regulatory networks underlying biomineralized skeletons in Bilateria": Supplementary Information.pdf

**Supplementary Note 1: Improvement of gene annotation**

A new full-length transcriptome (mixed stages, mRNA) was generated from six shell-forming stages/tissues (gastrula, trochophore, D-shaped larva, later D-shaped larva, juvenile, and adult mantle) using Iso-seq on the PacBio Sequel II platform. From 28,238,173 raw subreads, a total of 458,171 circular consensus sequence (CCS) reads were obtained (Supplementary Fig. 2). After filtering, 348,493 high-quality full-length reads were retained. Of these, 346,373 reads were mapped to the *C. nippona* genome. Transcript models were subsequently reconstructed, collapsed to remove redundancy, and merged with our previously published Iso-seq dataset[1]. Low-support isoforms were filtered out, and the resulting transcript models were further refined using short-read RNA-seq data and rule-based filtering to enhance annotation accuracy and confidence. In total, 89,768 high-confidence isoforms were identified, of which 85,575 were predicted to encode proteins. BUSCO analysis[2] of these protein-coding isoforms using the metazoan dataset (n=954) revealed 81.2% completeness (28.0% single-copy, 53.2% duplicated), with 1.4% fragmented and 17.4% missing BUSCOs, indicating a high level of transcript completeness. These isoforms were then merged with gene models from the MAKER annotation using AGAT (v.1.3.0) (https://github.com/NBISweden/AGAT), resulting in a final non-redundant gene annotation containing 29,577 gene models (Supplementary Data 1).

Among the 29,577 predicted gene models, 28,663 were identified as protein-coding genes and 914 as non-coding genes, with an average gene length of 10,641 bp

(Supplementary Data 1). The total number of predicted genes is comparable to that of our previous annotation[1], while both the average gene length (10,641 bp) and CDS length (1,521 bp) in this study are longer than those of the previous annotation (9,005 bp and 1,476 bp, respectively) [1]. In addition, we annotated a total of 117,732 isoforms, strongly supported by full-length transcript evidence (Supplementary Data 1). Importantly, the completeness of the predicted protein-coding gene set reached 96.4% and 96.6% for metazoan and molluscan orthologs, respectively (Supplementary Fig. 3a), and 99.66% of protein-coding genes annotated by at least one public database (Supplementary Fig. 3b). These results collectively highlight the notable improvement and high quality of the gene annotation achieved in this study.

**Supplementary Note 2: GO enrichment of each gene cluster**

*K*-means clustering was performed on the full transcriptome dataset to identify potential gene sets and biological processes related to specific developmental stages, yielding ten gene clusters that broadly corresponded to the major developmental stages (Fig. 2a). Gene ontology (GO) enrichment analyses of these clusters revealed distinct biological programs underpinning each cluster and developmental stage (Supplementary Fig 5, Supplementary Data 3).

Clusters C1 and C2 were predominantly associated with the gastrula stage, encompassing the maternal-to-zygotic transition and rapid cell proliferation. Cluster C1 was enriched for GO terms related to mRNA processing, RNA splicing, and translational initiation, as well as histone modification and chromatin remodeling, consistent with a role in maternal mRNA clearance and zygotic genome activation. Cluster C2 showed enrichment in DNA replication, chromosome segregation, and double-strand break repair, reflecting the mitotic activity during embryonic development. These patterns are consistent with transcriptomic dynamics previously reported in *Mytilus galloprovincialis*[3].

Cluster C3, highly expressed during the trochophore stage, was enriched for glycoprotein biosynthesis, endoplasmic reticulum stress response, and vesicle transport, reflecting the larval organogenesis, cuticle formation, and epithelial morphogenesis. In contrast, Cluster C4, marking the transition from trochophore to D-larva, was dominated by genes involved in cilium assembly, intraciliary transport, and axoneme organization, consistent with the development of motile and sensory cilia essential for larval movement and environmental sensing. Clusters C5 and C6, specific to the D-shaped larval stages, exhibited functional signatures linked to both metabolism and shell formation. These included GO terms related to lipid oxidation, peroxisomal transport, and motile cilium assembly, suggesting a coupling between metabolic energy production and larval motility. In addition, genes within cluster C5 and C6 were associated with ribosomal structure, extracellular matrix and calcium ion transport, supporting roles in protein biosynthesis (e.g., for shell matrix proteins) and ion regulation during larval shell formation. These transcriptional signatures closely parallel those identified in *Patinopecten yessoensis*[4], in which gene modules upregulated during the trochophore and D-larval stages were enriched for processes related to ciliary motilityand shell formation, including chitin metabolism and calcium ion binding. These observations suggest the existence of a conserved developmental framework among bivalves, wherein the coordinated activation of gene regulatory programs governing larval motility and biomineralization facilitates the construction of larval shell during early ontogeny.

At the juvenile stage, clusters C7 and C8 were associated with benthic life and immune activation. C7 was characterized by GO terms related to chitin metabolism and extracellular matrix organization, likely supporting adult shell formation. C8 showed enrichment for antiviral defense responses, such as leukocyte differentiation, lymphocyte activation, and response to virus, indicating an activation of innate immunity following settlement. Clusters C9 and C10, predominantly expressed in the adult mantle, were enriched in actin cytoskeleton organization, Rho GTPase signaling, actomyosin structure formation, and striated muscle development. These functions are likely associated with tissue homeostasis and functions involved in mantle activity and adult shell formation.

In summary, these findings support the delineation of six major developmental stages in the oyster ontogeny, each characterized by a distinct combination of anatomical features and transcriptional programs. The stage-specific enrichment of GO terms highlights both evolutionarily conserved processes, such as ciliogenesis and innate immunity, and Mollusca-specific innovations, such as peroxisomal energy metabolism during larval shell formation and cytoskeletal remodeling in the adult mantle. These features likely contribute to ecological resilience and morphological diversification in molluscan shell formation.

**Supplementary Note 3: Classification of regulatory divergence among functionally diverged paralogous genes**

To define the regulatory divergence of biomineralization gene paralogs, we compared their expression levels and enrichment of active histone modifications (H3K4me3 and H3K27ac) between D-shaped larvae and adult mantle tissue. A gene was considered regulatory specialized for larval shell formation if it showed significantly higher expression and stronger enrichment of associated histone marks in larvae. In contrast, a gene was considered regulatory specialized for adult shell formation if it exhibited higher expression and histone signals in the adult mantle. Gene pairs with no significant differences in both expression and regulatory marks between stages were defined as shared regulators for both larval and adult shell formation. Following this framework, we defined three categories of regulatory divergence among functionally specialized paralogous gene pairs:

a) No divergence (No): both genes exhibit similar regulatory profiles across stages, with no significant differences in H3K4me3 or H3K27ac levels between larval and adult stages. This suggests a lack of regulatory asymmetry, despite functional divergence.
b) One-sided divergence (One): only one gene in the pair shows stage-specific regulatory enrichment, indicated by a marked increase in active histone modifications in either larvae or adults. This suggests that divergence in only one paralog may drive functional specialization.
c) Both-sided divergence (Both): both genes exhibit distinct regulatory profiles

between stages, with one gene marked in larvae and the other in adults. This indicates coordinated, reciprocal regulatory specialization, where both paralogs have acquired distinct cis-regulatory landscapes adapted to stage-specific functions.

These classifications allowed us to systematically link expression specialization with regulatory evolution and support the role of epigenetic remodeling in the divergence of biomineralization gene function across oyster life stages.

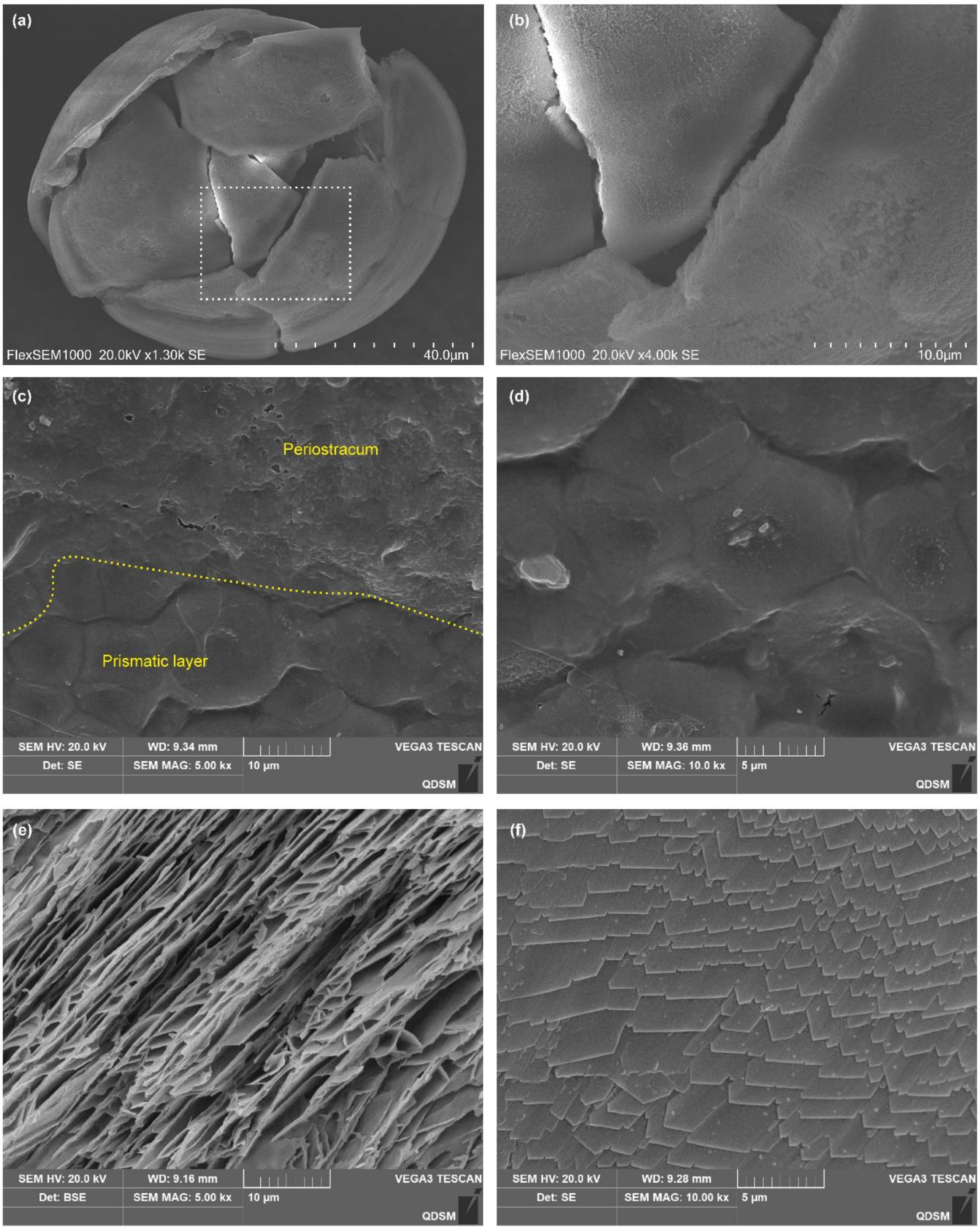


**Supplementary Fig 1. Morphology and structure of larval and adult shells in *C. nippona* showed by scanning electron microscopy (SEM). (a)** Aragonitic shell of the D-stage larva at 72 hours post fertilization. **(b)** Magnified view of the region outlined by the white dashed box in **(a)**. **(c)** Outer surface of the adult calcitic shell. The area above the yellow dashed line corresponds to the periostracum, while the area below corresponds to the prismatic layer. **(d)** Outer surface of the prismatic layer in the adult calcitic shell. **(e)** Inner surface of the chalky layer in the adult calcitic shell. **(f)** Inner surface of the foliated layer in the adult calcitic shell.

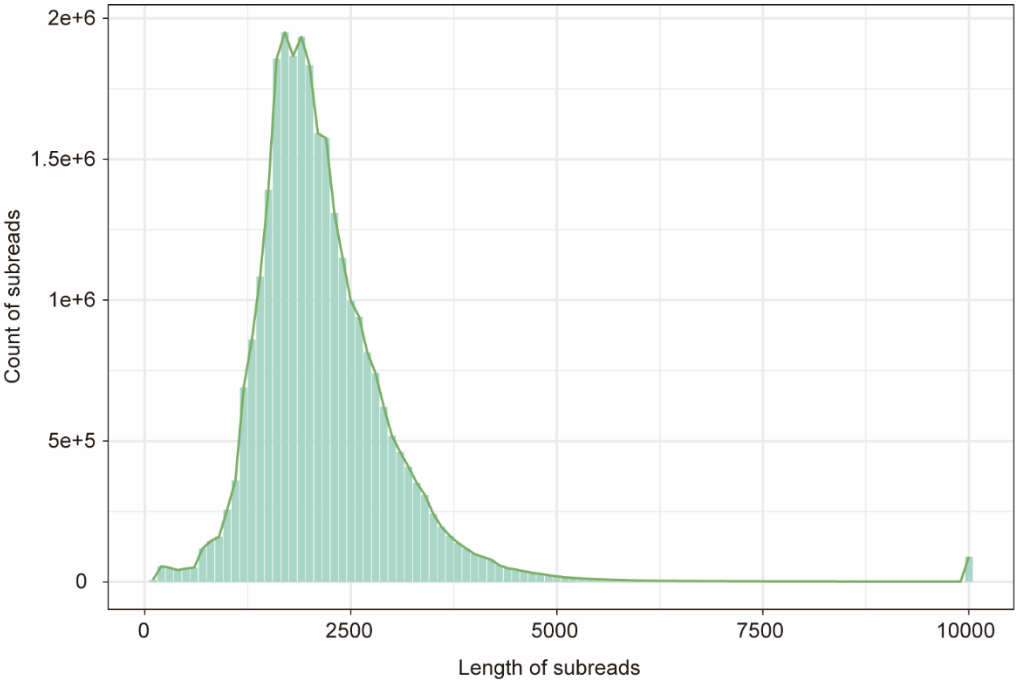


**Supplementary Fig 2. Statistics of length distributions of the raw subreads from Iso-seq.**

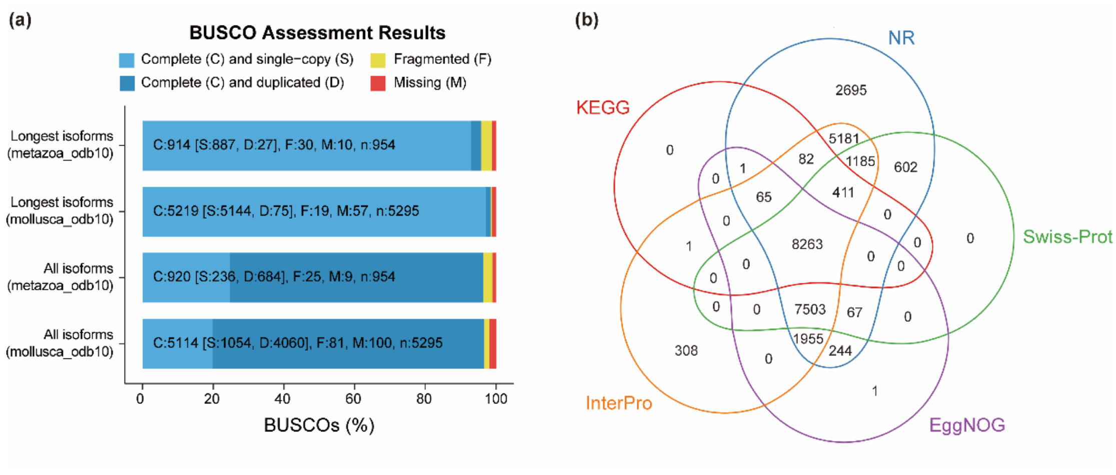


**Supplementary Fig 3. Quantify assessment of improved gene predictions in *C. nippona*. (a)** BUSCO analysis of the predicted protein sequences using both the longest isoforms and all isoforms, based on the metazoa_odb10 and mollusca_odb10 databases. **(b)** Venn diagram showing the overlap of functionally annotated protein-coding genes across five databases: NR, Swiss-Prot, KEGG, InterPro, and EggNOG. The numbers represent the counts of genes uniquely or commonly annotated in each database.

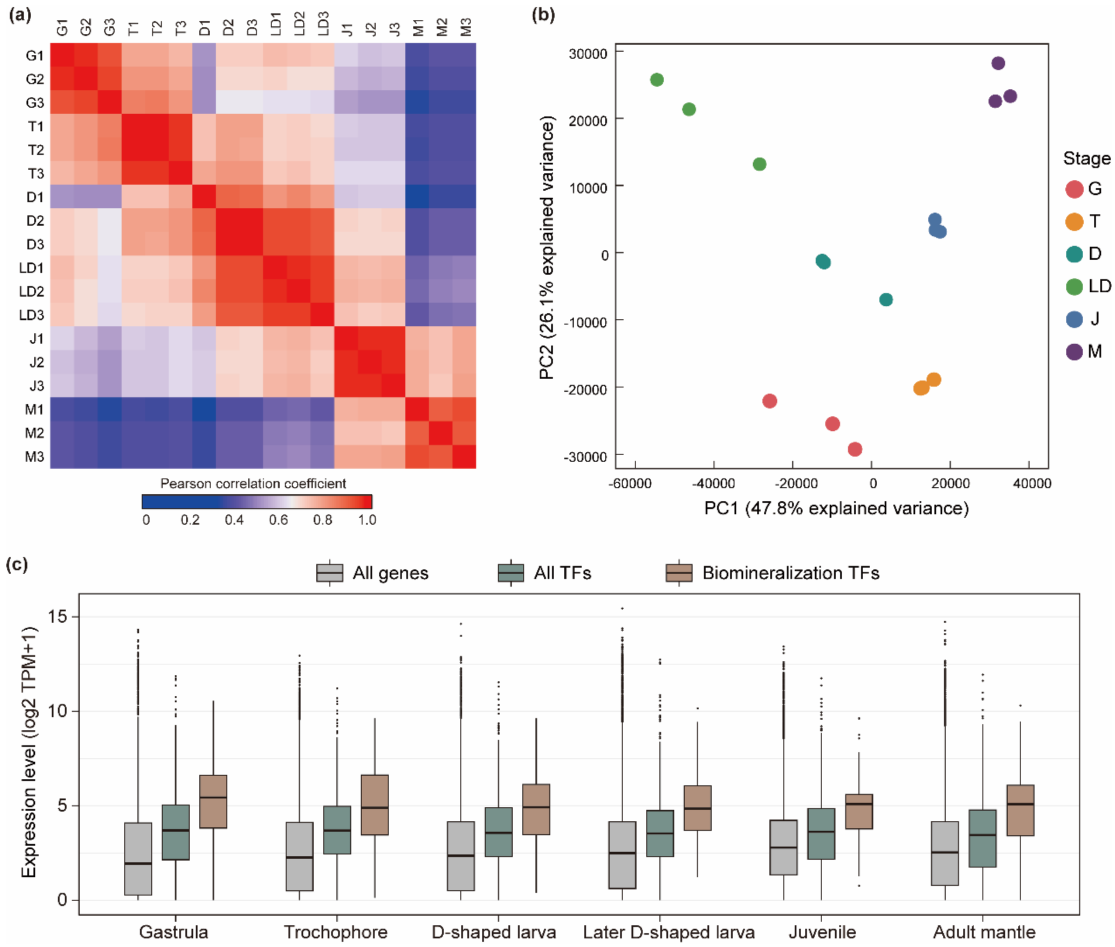


**Supplementary Fig 4. Gene expression dynamics across six developmental stages in *C. nippona*. (a)** Correlation matrix based on gene expression profiles, indicating strong within-stage consistency and distinct stage-specific transcriptomic signatures. **(b)** Principal component analysis (PCA) of gene expression, showing stage-wise separation along PC1 (47.8% variance explained) and PC2 (26.1% variance explained). **(c)** Expression levels of all genes, all transcription factors (TFs), and biomineralization TFs across six developmental stages. Biomineralization TFs show consistently higher expression levels than other TFs or all genes, particularly during shell-forming stages. Abbreviations: G gastrula; T trochophore; D D-shaped larva; LD later D-shaped larva; J juvenile; M adult mantle.

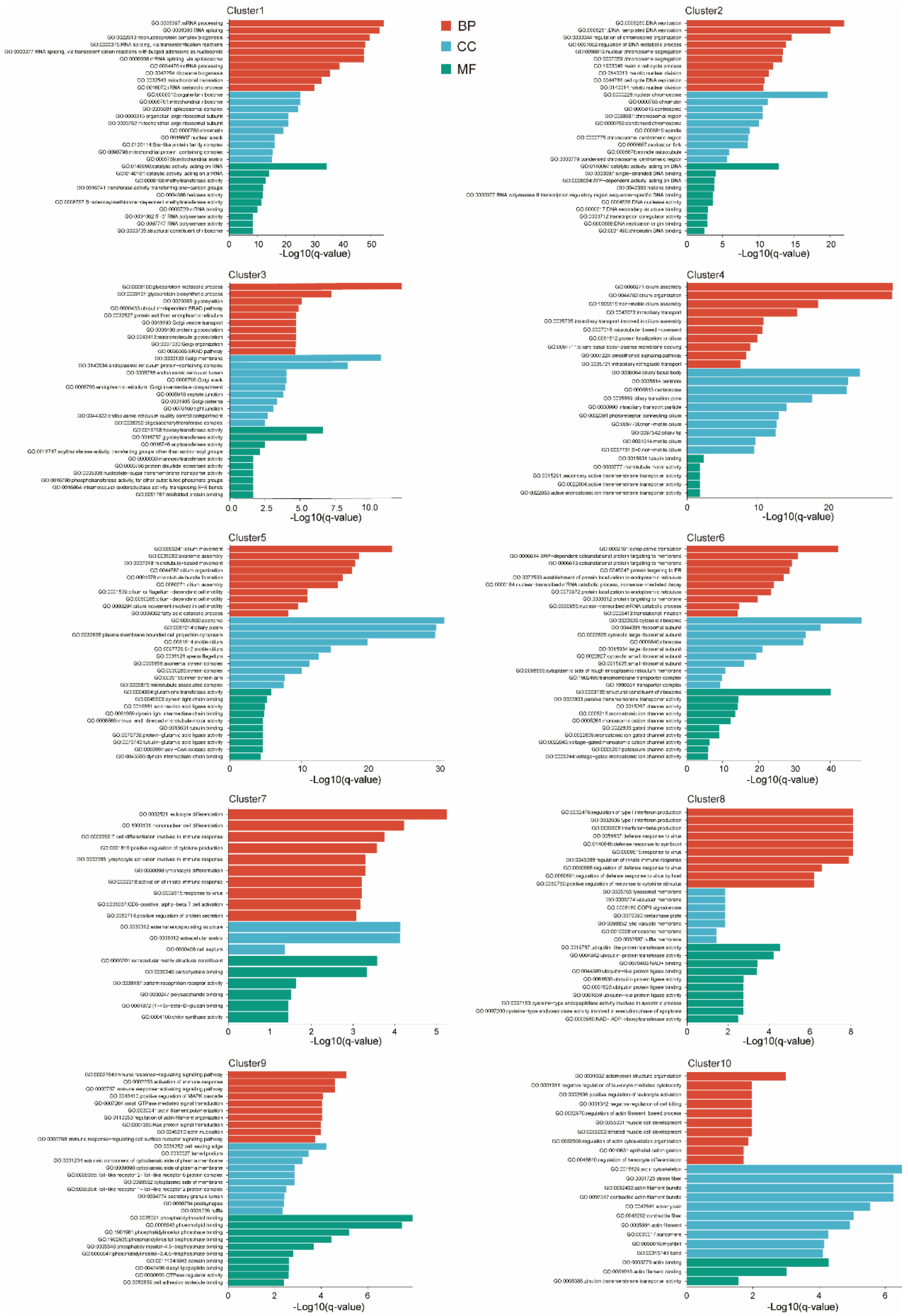


**Supplementary Fig 5. GO terms enrichment of RNA-seq cluster of *C. nippona*.**

Bar plots depicting adjusted *P*-values (q-value) of the top 10 GO terms for biological process (BP), cell component (CC), and molecular function (ME).

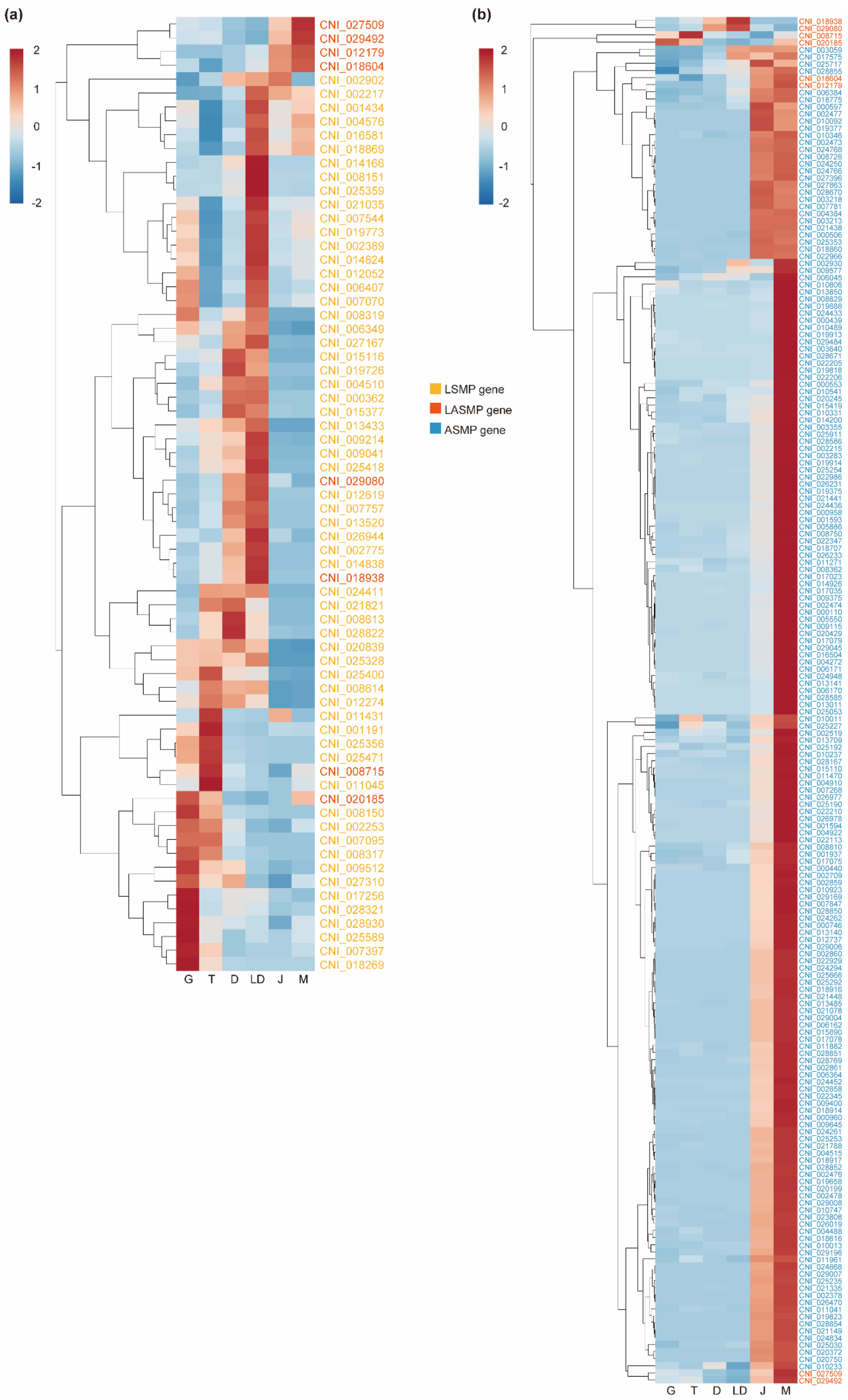
(a)
(b)
2
1
0
-1
-2
LSMP gene
LASMP gene
ASMP gene
CNI_027509
CNI_029492
CNI_012179
CNI_018604
CNI_002902
CNI_002217
CNI_001434
CNI_004576
CNI_016581
CNI_018869
CNI_014166
CNI_008151
CNI_025359
CNI_021035
CNI_007544
CNI_019773
CNI_002389
CNI_014824
CNI_012052
CNI_006407
CNI_007070
CNI_008319
CNI_006349
CNI_027167
CNI_015116
CNI_019726
CNI_004510
CNI_000362
CNI_015377
CNI_013433
CNI_009214
CNI_009041
CNI_025418
CNI_029080
CNI_012619
CNI_007757
CNI_013520
CNI_026944
CNI_002775
CNI_014838
CNI_018938
CNI_024411
CNI_021821
CNI_008613
CNI_028822
CNI_020839
CNI_025328
CNI_025400
CNI_008614
CNI_012274
CNI_011431
CNI_001191
CNI_025356
CNI_025471
CNI_008715
CNI_011045
CNI_020185
CNI_008150
CNI_002253
CNI_007095
CNI_008317
CNI_009512
CNI_027310
CNI_017256
CNI_028321
CNI_028930
CNI_025589
CNI_007397
CNI_018269
G T D LD J M
G T D LD J M

**Supplementary Fig 6. Gene expression patterns of larval (a) and adult (b) shell matrix proteins across six developmental stages in *C. nippona*.** Abbreviations: G gastrula; T trochophore; D D-shaped larva; LD later D-shaped larva; J juvenile; M adult mantle. LSMP larval shell matrix protein; LASMP larval and adult shell matrix protein; ASMP adult shell matrix protein.

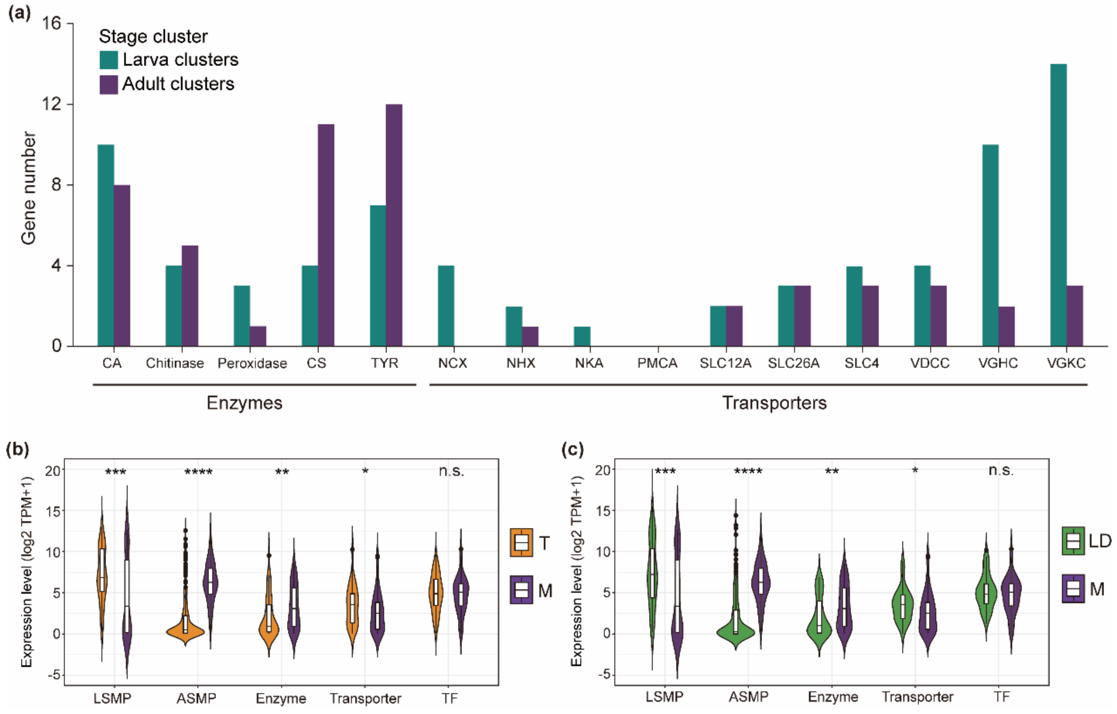


**Supplementary Fig 7. Expression patterns of biomineralization effector genes in *C. nippona*.** **(a)** Number of genes in each biomineralization effector family identified in larval and adult expression clusters. **(b, c)** Comparison of expression levels for each effector category between the trochophore and adult stages **(b)**, or between the later D-shaped larva and adult stages **(c)**. Boxplots include a median with quartiles and outliers above the top whisker. Statistical significance was assessed using a two-sided Wilcoxon rank-sum test (*$P < 0.05$; **$P < 0.01$; ***$P < 0.001$; ****$P < 0.0001$; n.s., no significance). Abbreviations: LSMP larval shell matrix protein; ASMP adult shell matrix protein; CA carbonic anhydrase; CS chitin synthase; TYR tyrosinase; NCX $Na^+/Ca^{2+}$-exchangers; NHX $Na^+/H^+$ exchangers; NKA $Na^+/K^+$-ATPase; PMCA $Ca^{2+}$-ATPase; VGHC voltage-gated $H^+$ channel; VGHC voltage-gated $K^+$ channel; VDCC voltage-dependent $Ca^{2+}$ channel; SLC solute carrier; TF transcription factor.

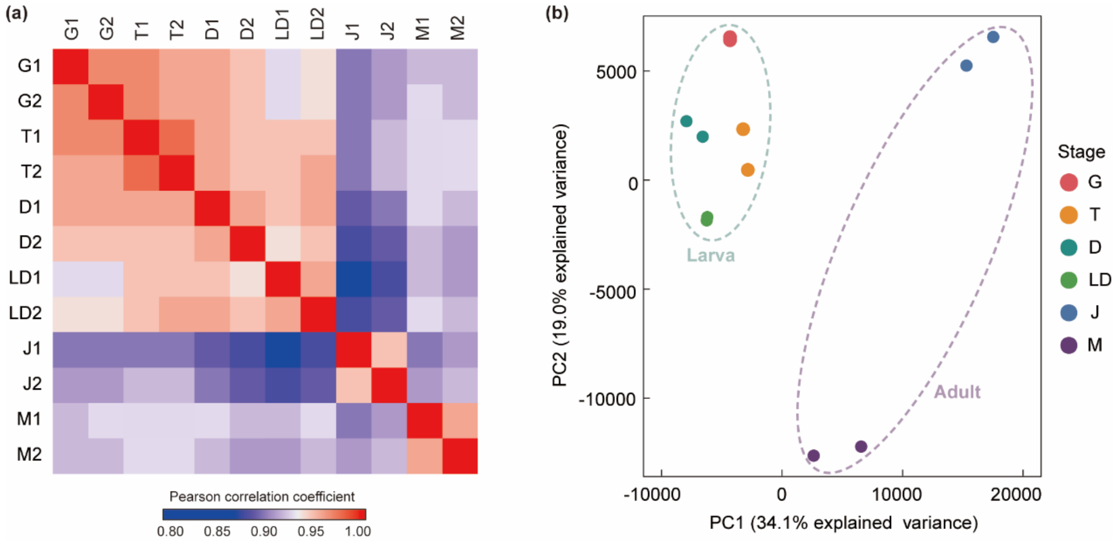


**Supplementary Fig 8. Chromatin dynamics across six developmental stages or tissues in *C. nippona*. (a)** Correlation matrix based on peak accessibility of the consensus ATAC-seq peak. **(b)** PCA of ATAC-seq samples based on peaks as in **(c)**, showing stage-wise separation along PC1 (34.1% variance explained) and PC2 (19.0% variance explained). The green and purple dashed circles outline the distribution of larval and adult samples, respectively.

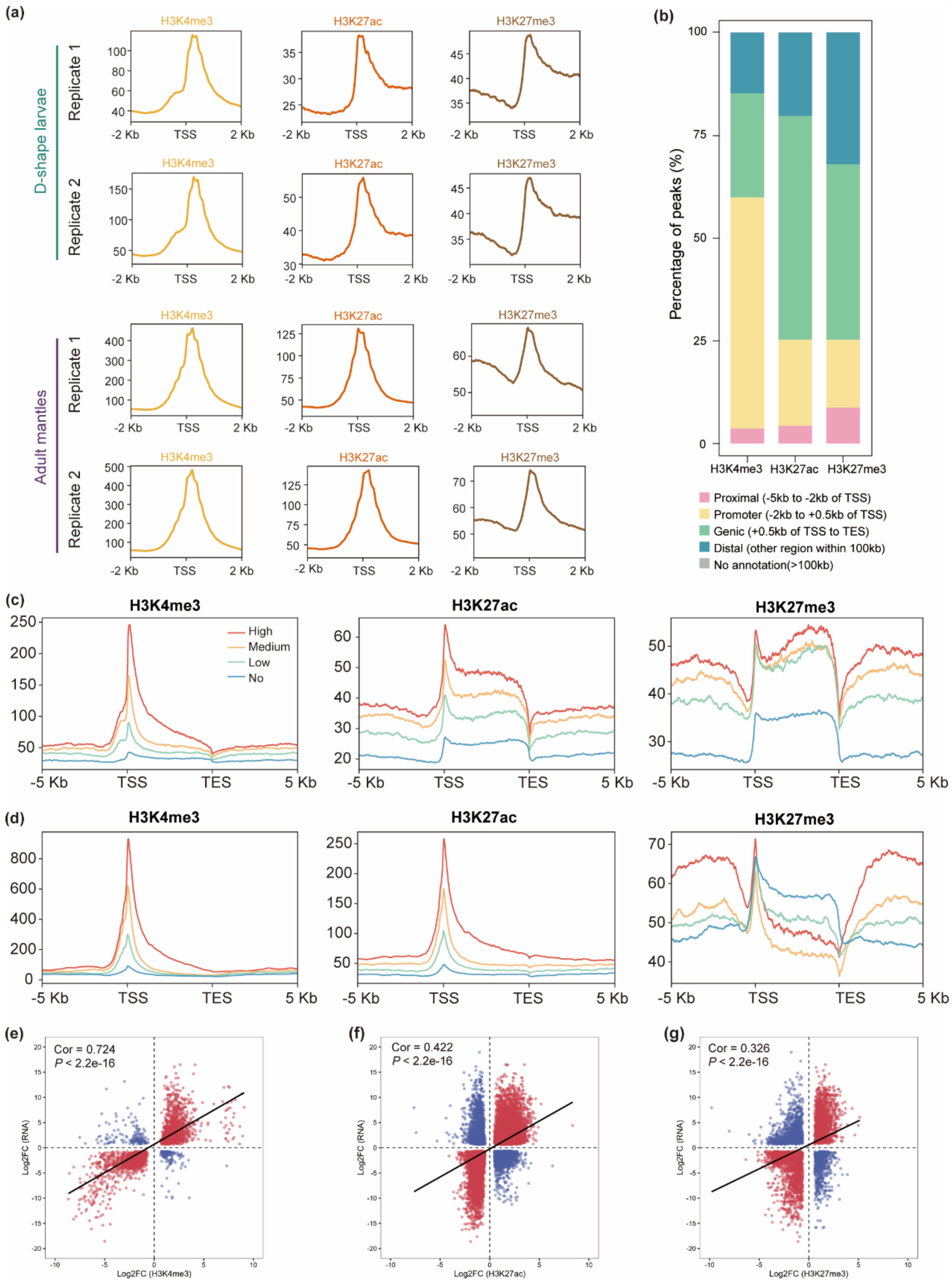


**Supplementary Fig 9. Genome-wide mapping of three histone marks (H3K4me3, H3K27ac, and H3K27me3) in D-shape larvae and adult mantles of *C. nippona* and their correlation with gene expression patterns. (a)** Summary plots of three histone mark enrichment around transcription start sites (TSS; ±2 Kb). **(b)** Genome-wide distribution of peaks of histone marks. **(c, d)** The levels of three histone mark in genes with different transcription levels in D-shape larvae **(c)** and adult mantles **(d)**. In the analysis, genes were grouped into four categories based on their expression levels: “no” indicates genes with low or no expression (TPM < 1); “low”, “medium”, and “high” represent the bottom, middle, and top one-third of expressed genes (TPM ≥ 1), respectively. The plots show the distribution of CUT&Tag signals for three histone modifications across these gene expression categories. **(e, f)** The correlation between variation fold changes in gene expression and H3K4me3 **(e)**, H3K27ac **(f)** or H3K27me3 **(g)** levels of genes in D-shape larvae and adult mantles. Pearson correlation coefficients were calculated to assess the linear relationship between the level changes of gene expression and histone modification. Statistical significance was determined using a two-sided Student’s t-distribution.

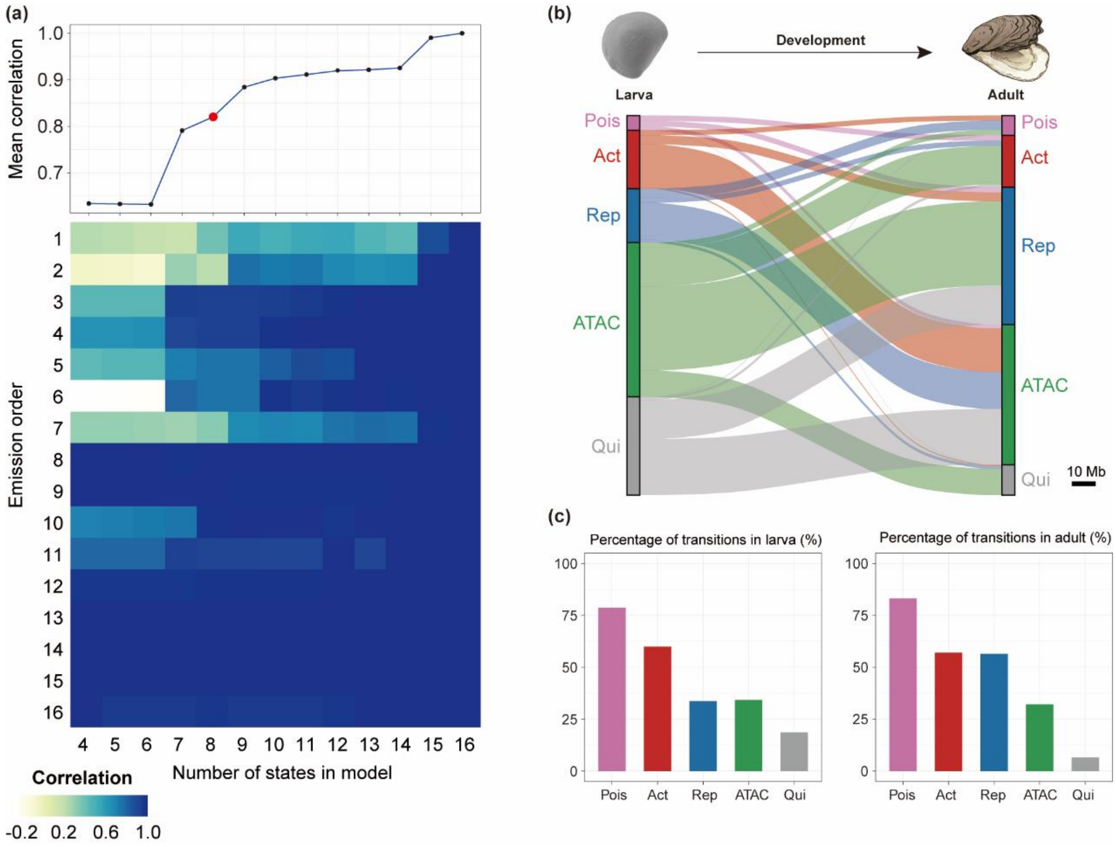


**Supplementary Fig 10. ChromHMM-inferred genome-wide chromatin states and their transitions from larva to adult stages in *C. nippona*. (a)** Correlation heatmap of different ChromHMM-trained chromatin state models. The color represents the level of correlation. **(b)** Chromatin state transitions of genes from larva (top) to adult (bottom) stages. **(c)** Top: percentage of chromatin states linked to genes in larvae that undergo transitions to other chromatin state in adults. Bottom: percentage of chromatin states linked to genes in adult that undergo transitions from other chromatin state in larvae.

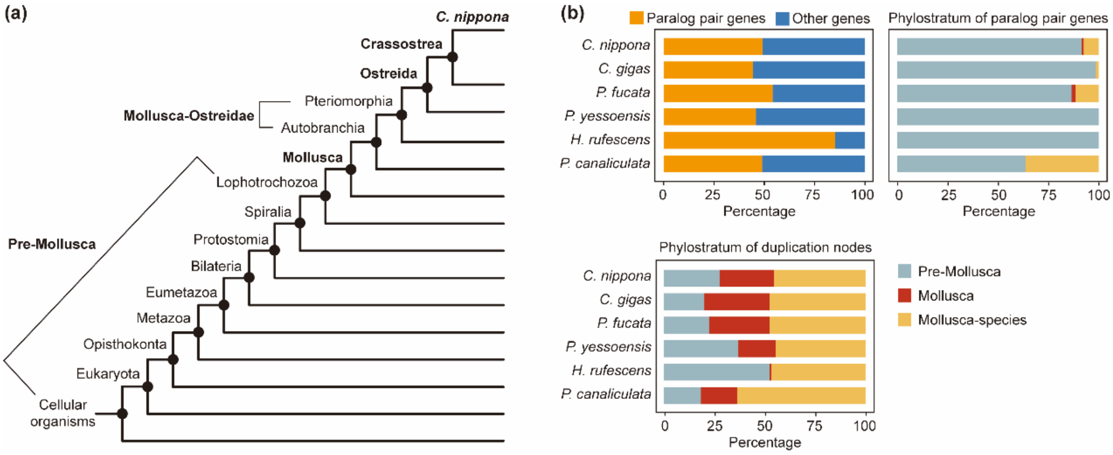


**Supplementary Fig 11. Gene age of paralogous genes and duplication dating of biomineralization effectors in *C. nippona*. (a)** Phylogenetically hierarchical classification of gene age in *C. nippona*. **(b)** The proportion of paralogous genes among biomineralization effectors in five molluscs, along with their phylostrata and duplication timing of paralog pairs.

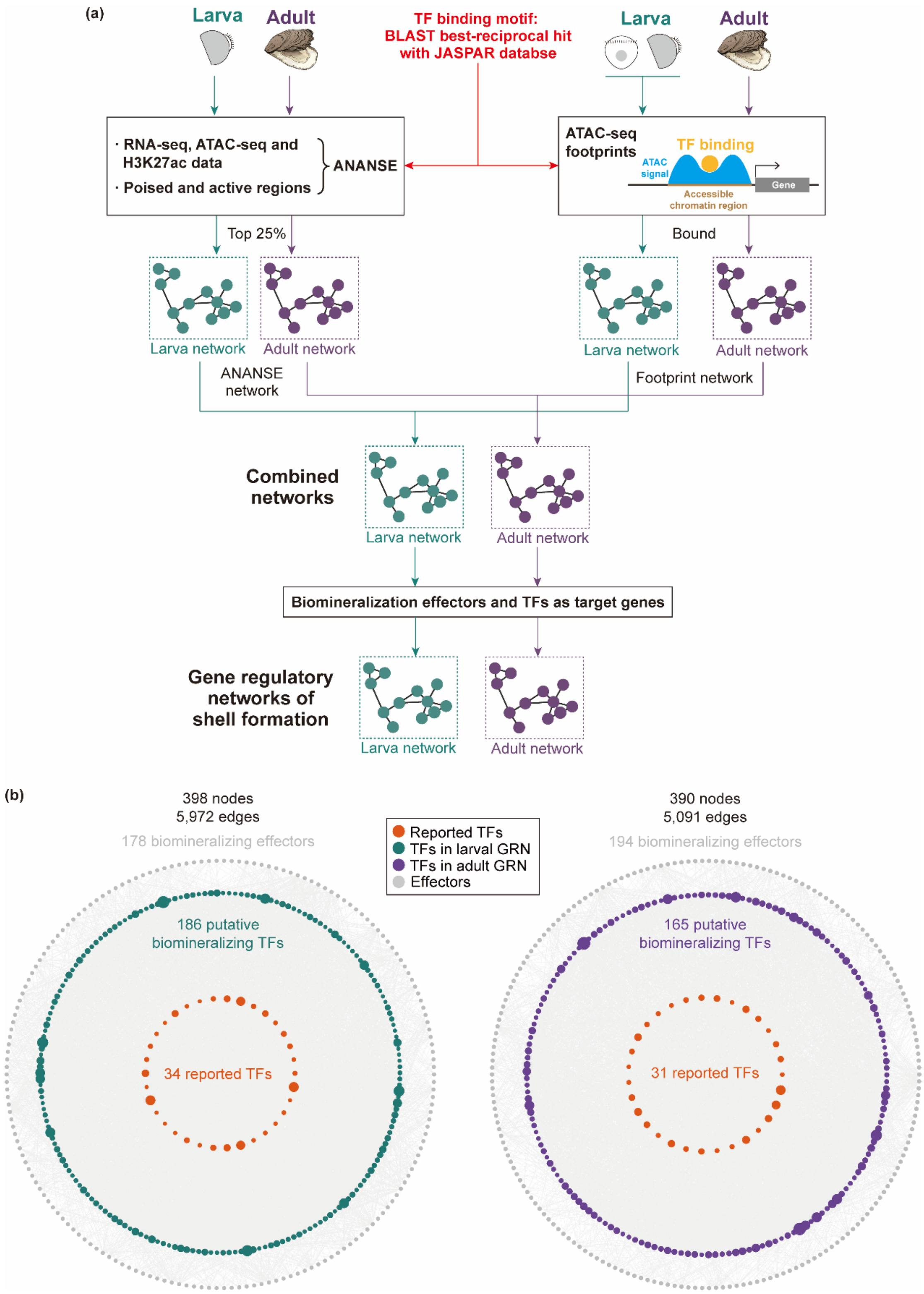


**Supplementary Fig 12. Construction of gene regulatory networks (GRNs) for larval and adult shell formation in *C. nippona*. (a)** Schematic pipeline illustrating networks of *cis*-regulatory. **(b)** Overview of GRNs of larval (left) and adult (right)

shell formation. Orange dots represent previously reported TFs, while green (left) and purple (right) dots represent TFs identified in larval and adult GRNs, respectively. Outer grey dots represent biomineralization effector genes. Larval and adult GRNs comprise 398 and 390 nodes, and 5,972 and 5,091 edges, respectively. A total of 220 and 196 putative biomineralization TFs were identified in the larval and adult networks, respectively, including 34 and 31 TFs previously reported in biomineralization. Dot size reflects the outdegree of each node, indicating the number of target genes regulated by a given TF.

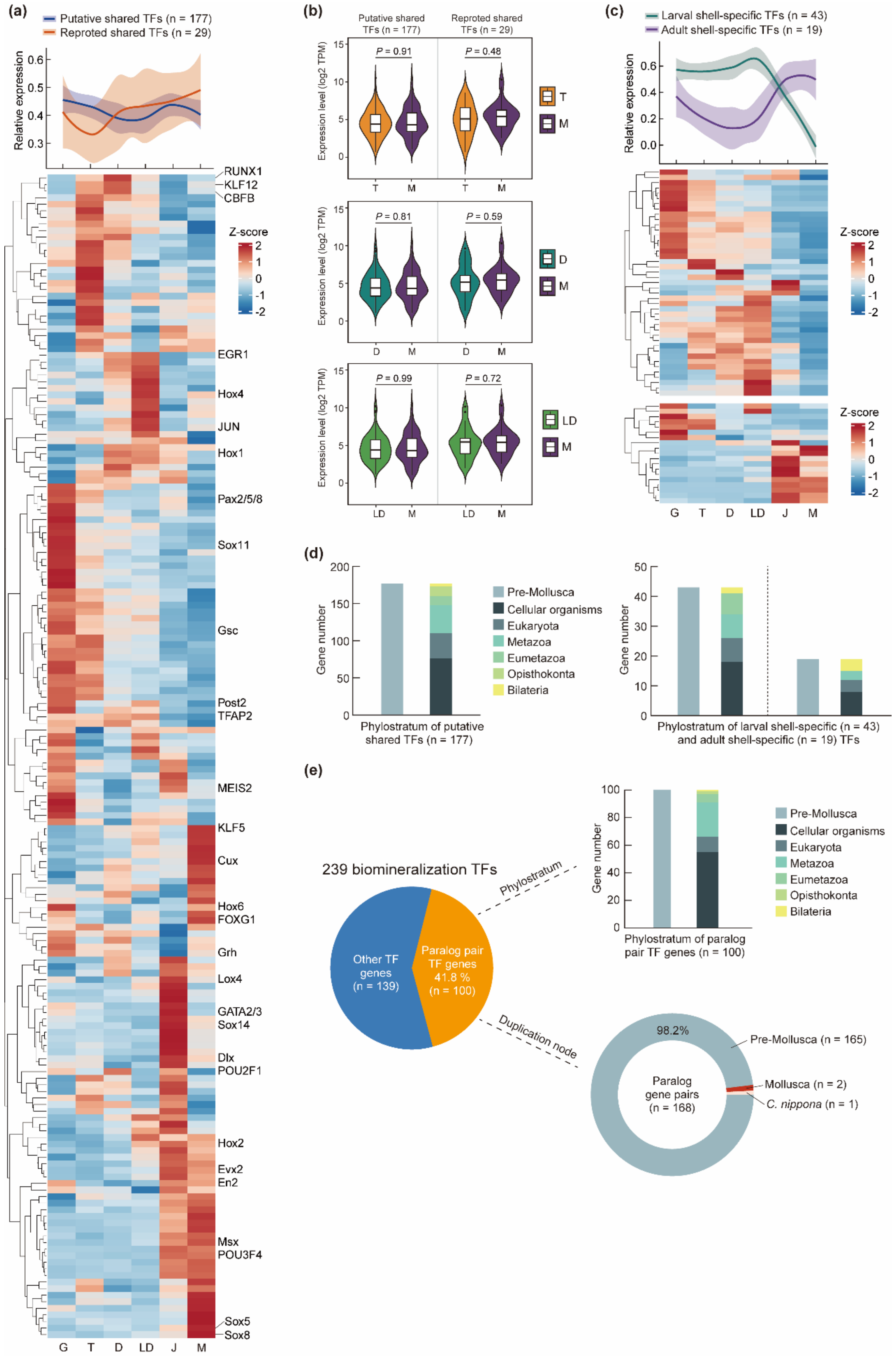


**Supplementary Fig 13. Gene expression profiles of putative biomineralization**

**TFs in *C. nippona* and their phylostrata. (a)** Gene expression dynamics (top) of 177 shared TFs (blue) predicted in both larval and adult GRNs, including 29 previously reported TFs (orange), across six developmental stages in *C. nippona*. Curves are locally estimated scatterplot smoothing (LOESS), colored shaded areas represent standard error of the mean. Gene names of 29 previously reported TFs are labeled in the right of heatmap (bottom). **(b)** Comparison of expression levels for TFs between larva and adult stages. Boxplots include a median with quartiles and outliers above the top whisker. The two-sided Wilcoxon rank-sum test was used to assess significance. **(c)** Gene expression dynamics of larval (green) or adult (purple) shell-specific TFs predicted in larval or adult GRNs, respectively, across six developmental stages in *C. nippona*. Curves are LOESS, colored shaded areas represent standard error of the mean. **(d)** Phylostratum of 177 shared TFs (left) and stage-specific TFs (right). All TFs are of pre-mollusca origin, and further classified as origin from Cellular organisms, Eukaryota, Metazoan, Eumetazoa, Opisthokonta and Bilateria. **(e)** Percentage of paralogous genes among biomineralization TFs, as well as the distribution of their phylostrata and duplication nodes. Abbreviations are the same as Supplementary Fig 3.

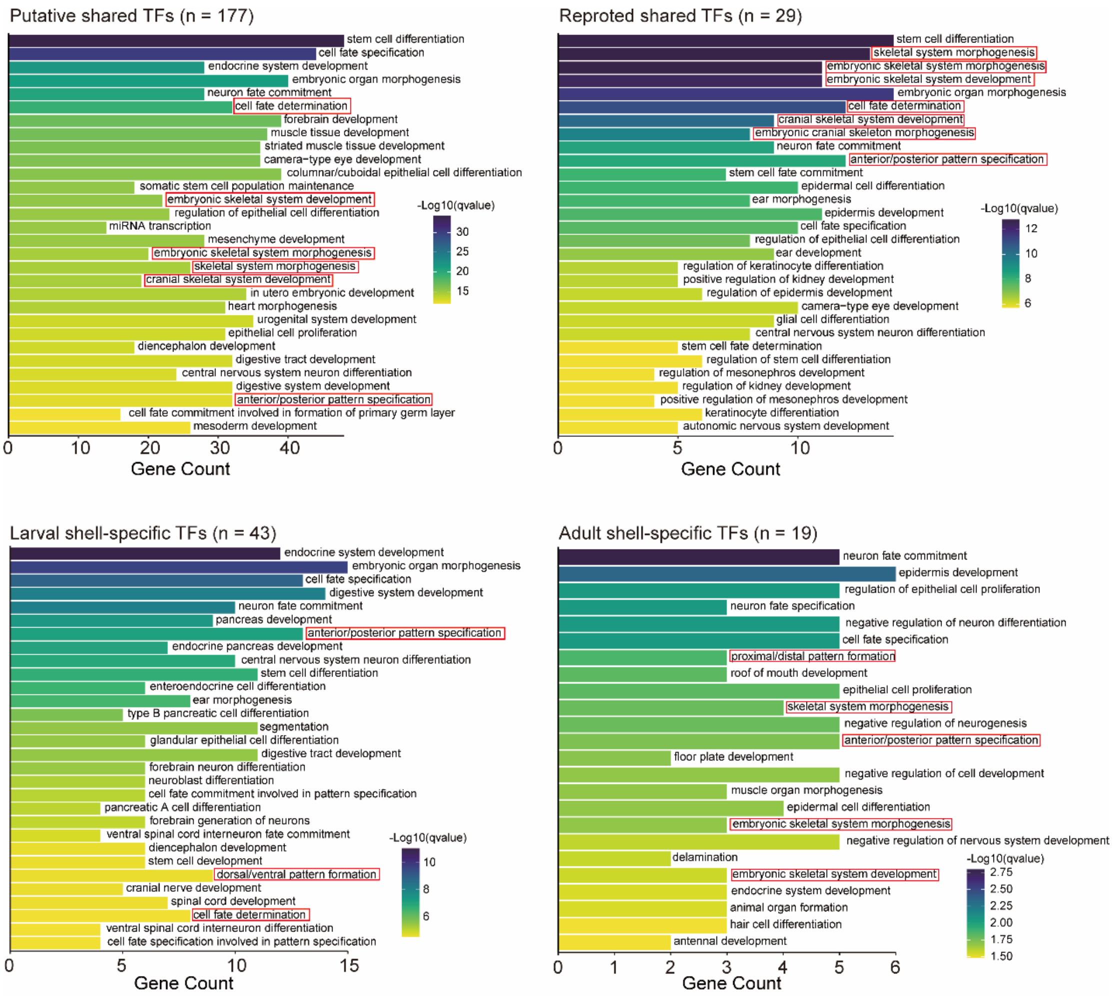


**Supplementary Fig 14. GO terms enrichment of biomineralization TFs in shell formation GRNs of *C. nippona*.** Bar plots depicting gene counts, while colors indicate adjusted *P*-values (qvalue) of the top 30 GO terms for biological process. Terms mentioned in the main text are highlighted with red boxes.

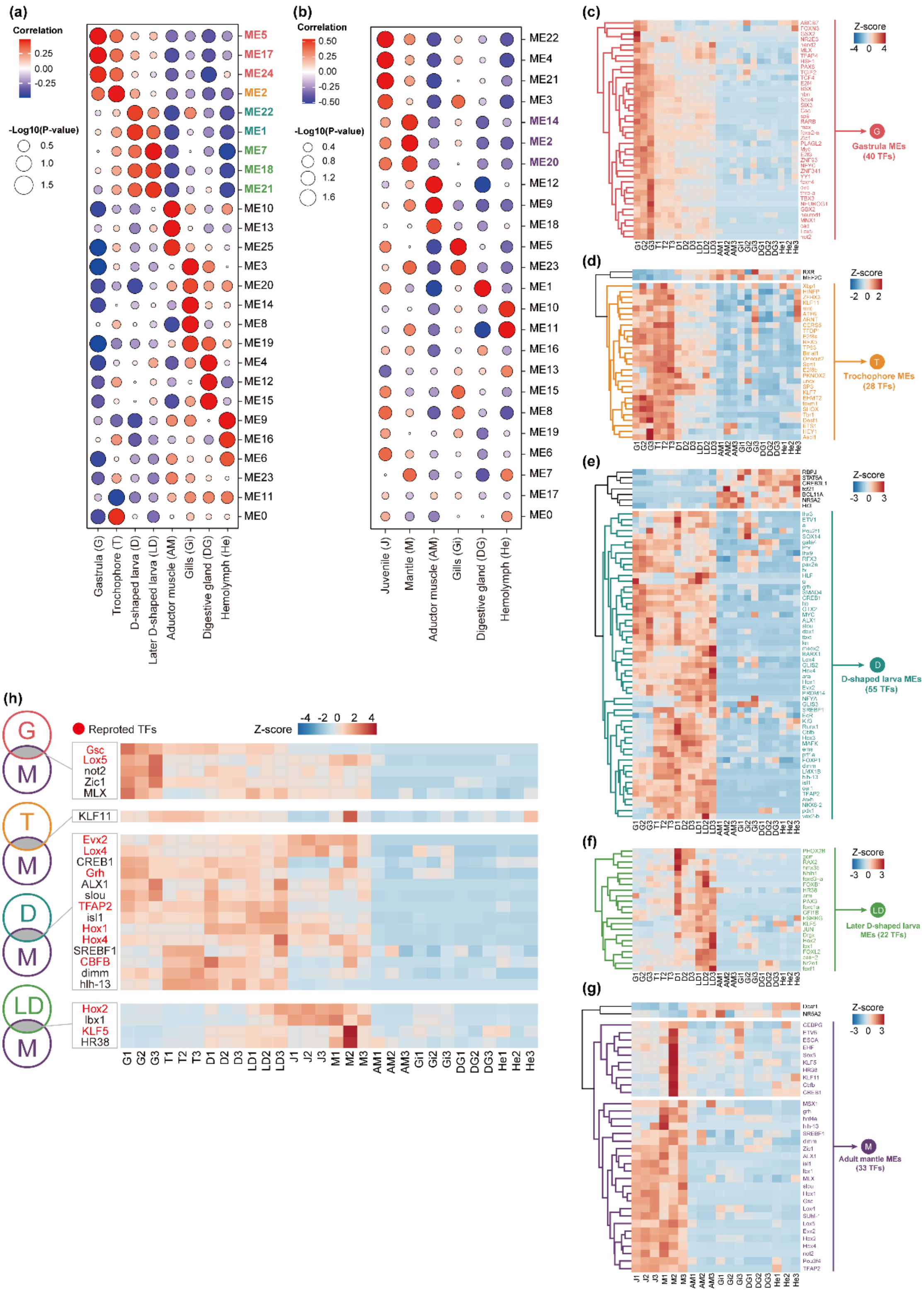


**Supplementary Fig 15. WGCNA analysis across developmental stages and adult tissues to identify of co-expressed TF genes in both larval and adult shell formation of *C. nippona*. (a)** Gene modules (MEs) associated with larval shell formation identified by WGCNA clustering of gene expression profiles of larval stages and non-shell-forming tissues: gastrula (salmon: ME5, ME17 and ME24), trochophore (orange: ME2), D-shape larva (green: ME1 and ME22), and later D-shape larva (medium green: ME7, ME18 and ME21). Dot size indicates the significance (-log10 *P*-value) of the correlation, and the color represents the correlation coefficient. **(b)** MEs associated with adult shell formation in the mantle tissue (purple: ME2, ME14 and ME20), identified through WGCNA clustering of gene expression profiles of juvenile and adult tissues. Correlation and *P*-value are also indicated by dot size and color, respectively. **(c-g)** Heatmaps showing gene expression patterns in MEs associated with larval stages and adult mantle: gastrula **(c)**, trochophore **(d)**, D-shape larva **(e)**, later D-shape larva **(f)**, and adult mantle **(g)**. TFs within MEs were hierarchically clustered to ensure stage- or tissue-specific expression. Genes labeled in black were excluded from the stage- or tissue-specific MEs. **(h)** Co-expressed TFs shared each larval stage and adult mantle, along with their expression profiles across developmental stages and adult tissues. Biomineralization TFs reported in previous studies (Supplementary Data 4) are highlighted in red.

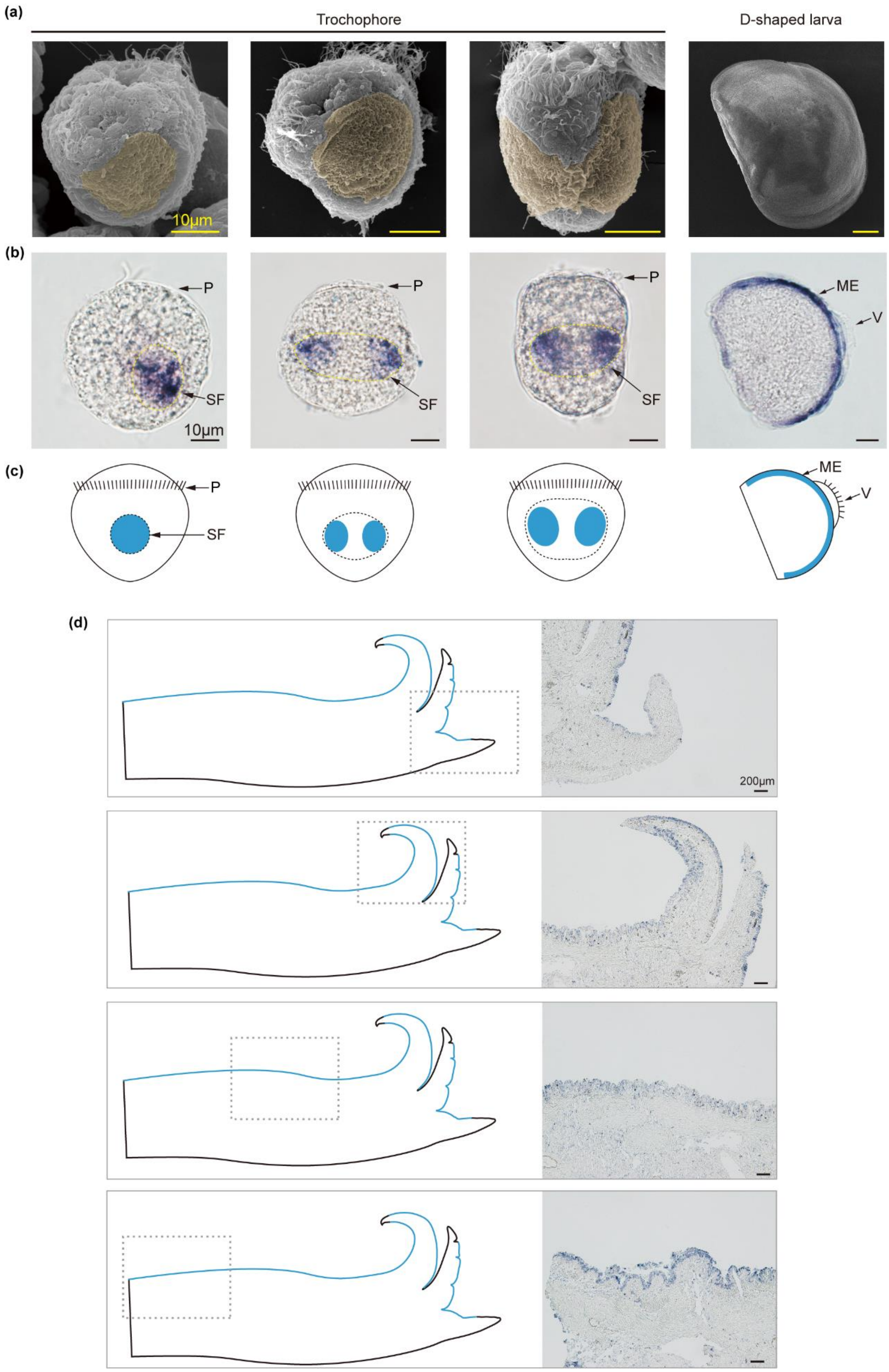


**Supplementary Fig 16. Spatial expression pattern of the *Hox4* gene in *C. nippona***

**larvae and adult mantle. (a)** Scanning electron micrographs (SEMs) of *C. nippona* larvae. SF in trochophore larvae is highlighted in yellow. Scale bar = 10 μm. **(b-c)** ISH results **(b)** and corresponding schematic illustration **(c)** showing *Hox4* expression (blue) in the SF regions of trochophore larvae and mantle edge (ME) of D-shape larvae. **(d)** Schematic diagram (left) showing the *Hox4* expression (blue) in the adult mantle. The gray dashed boxes indicate the region magnified in the ISH images on the right. Scale bar = 200 μm. Abbreviations are the same as Supplementary Fig 10.

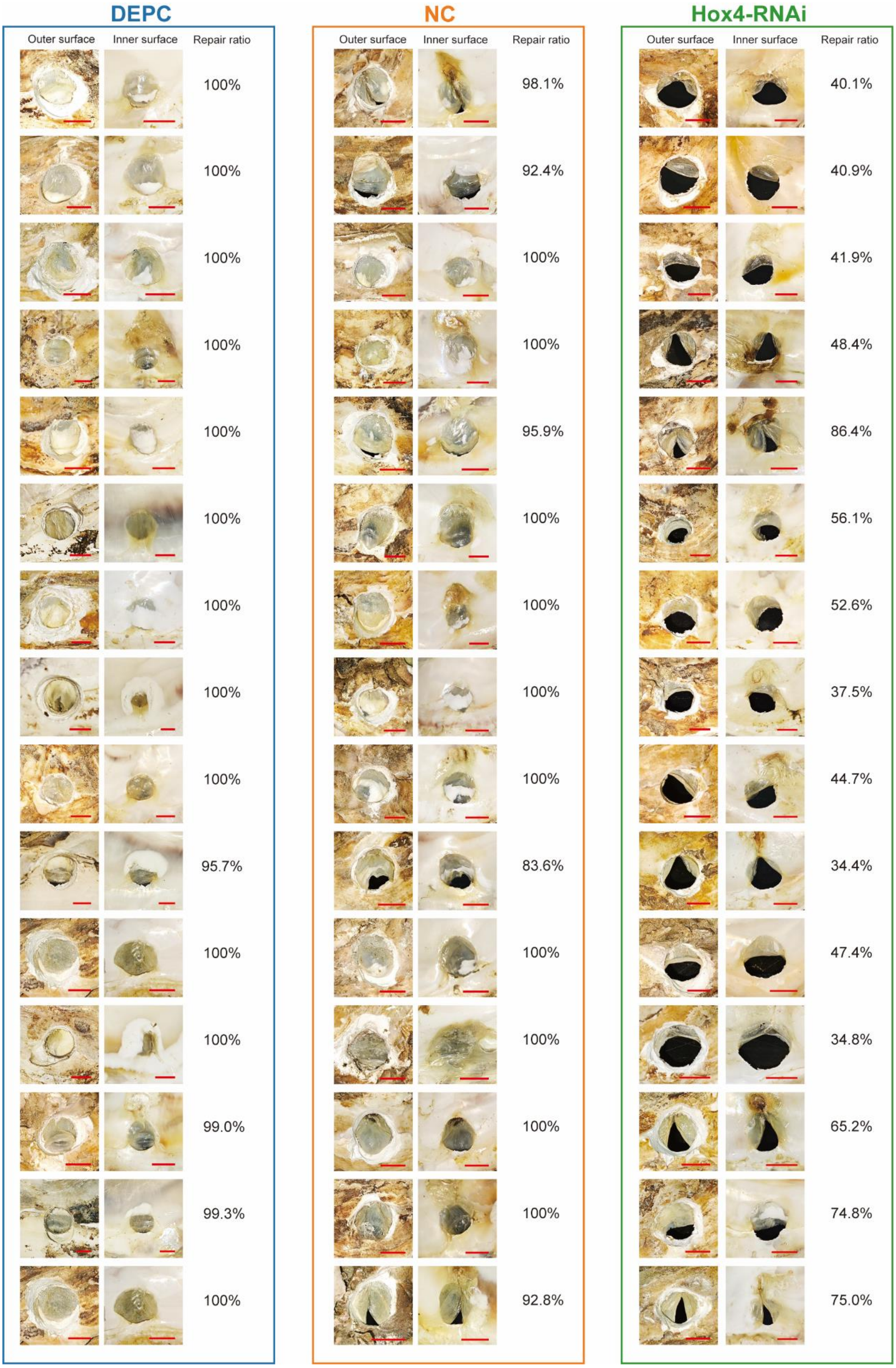


**Supplementary Fig 17. Bright-field photographs showing repaired shells six days after drilling.** Samples are arranged from left to right as DEPC (bule), NC (orange), and *Hox4*-RNAi (green) groups (n = 15 per group). Shell repair ratio was assessed as the repaired area divided by the area of the original hole.

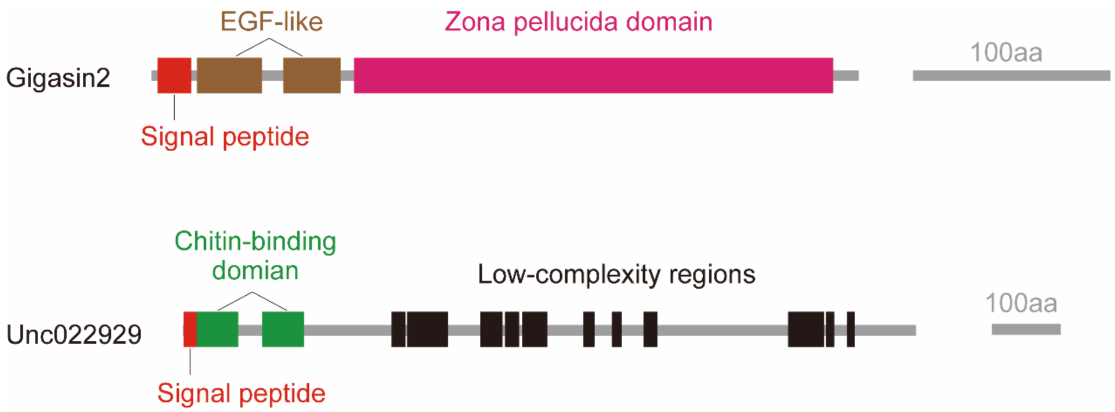


**Supplementary Fig 18. Protein domain structures of *Gigasin2* and *Unc022929* gene.** Red boxes represent signal peptide domains; brown: EGF-like domains (IPR000742); magenta: zona pellucida domains (IPR001507); green: chitin-binding domains (IPR002557); black: low-complexity regions.

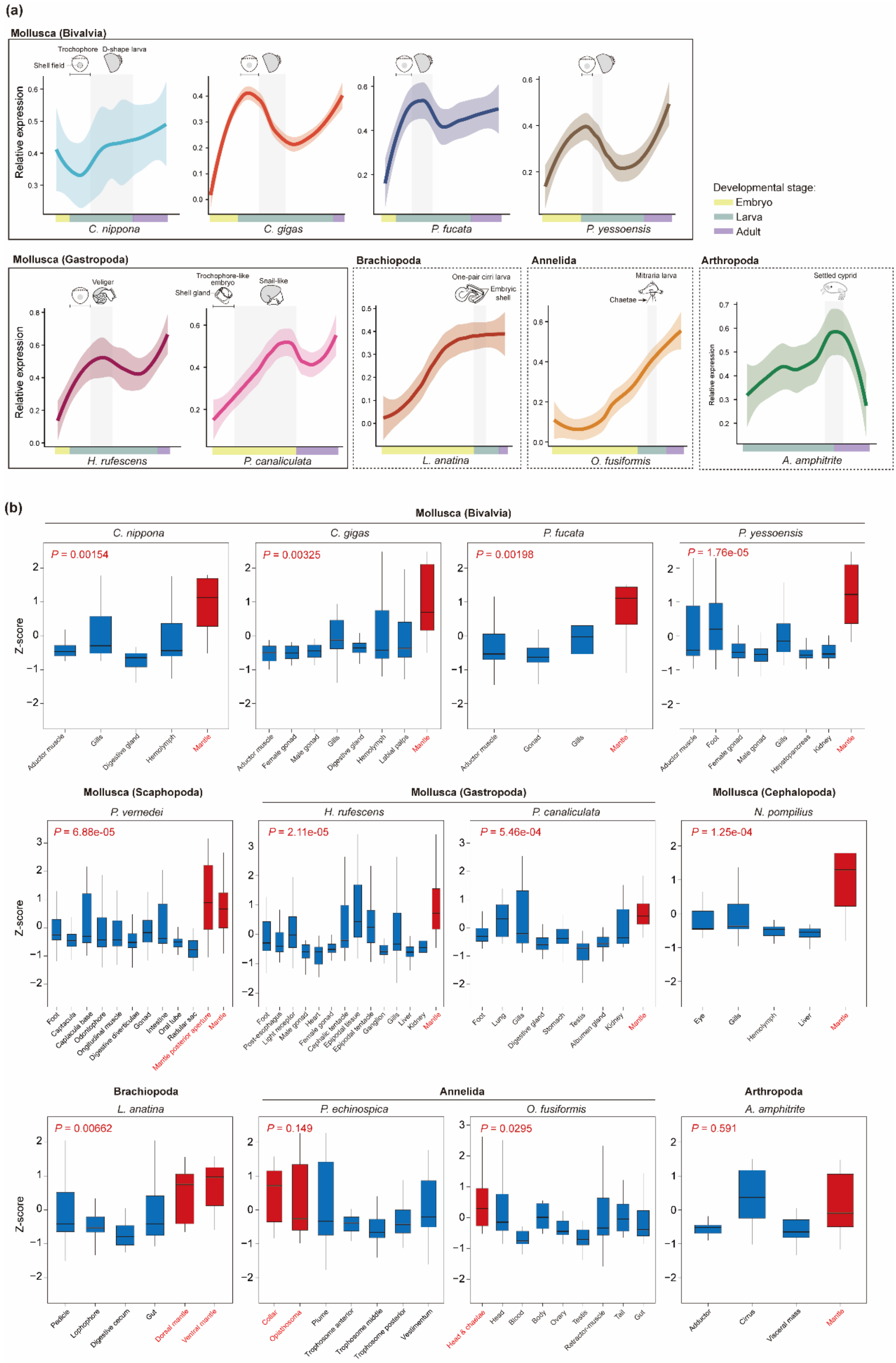


**Supplementary Fig 19. Gene expression patterns of 29 biomineralization TFs shared by the larval and adult shell formation GRNs across developmental stages**

**(a) or adult tissues (b) in bilaterians.** Expression curves are fitted using LOESS, with shaded areas indicating the standard error of the mean. In mollusks, the peak expression of these TFs occurs during the trochophore stage, which is a critical period for shell field development. In other lophotrochozoans, the peak expression typically coincides with early exoskeleton-forming larval stages. In contrast, the arthropod *Amphibalanus amphitrite* exhibit peak expression during the late larval stage prior to metamorphosis (settled cyprid). Moreover, in mollusks, as well as in *Lingula anatina* and *Owenia fusiformis*, these TF genes exhibit significantly higher expression levels (two-sided Wilcoxon rank-sum test: $P < 0.05$) in exoskeleton-forming organs. However, no significant tissue-specific expression differences were observed in *Paraescarpia echinospica* and *A. amphitrite*.

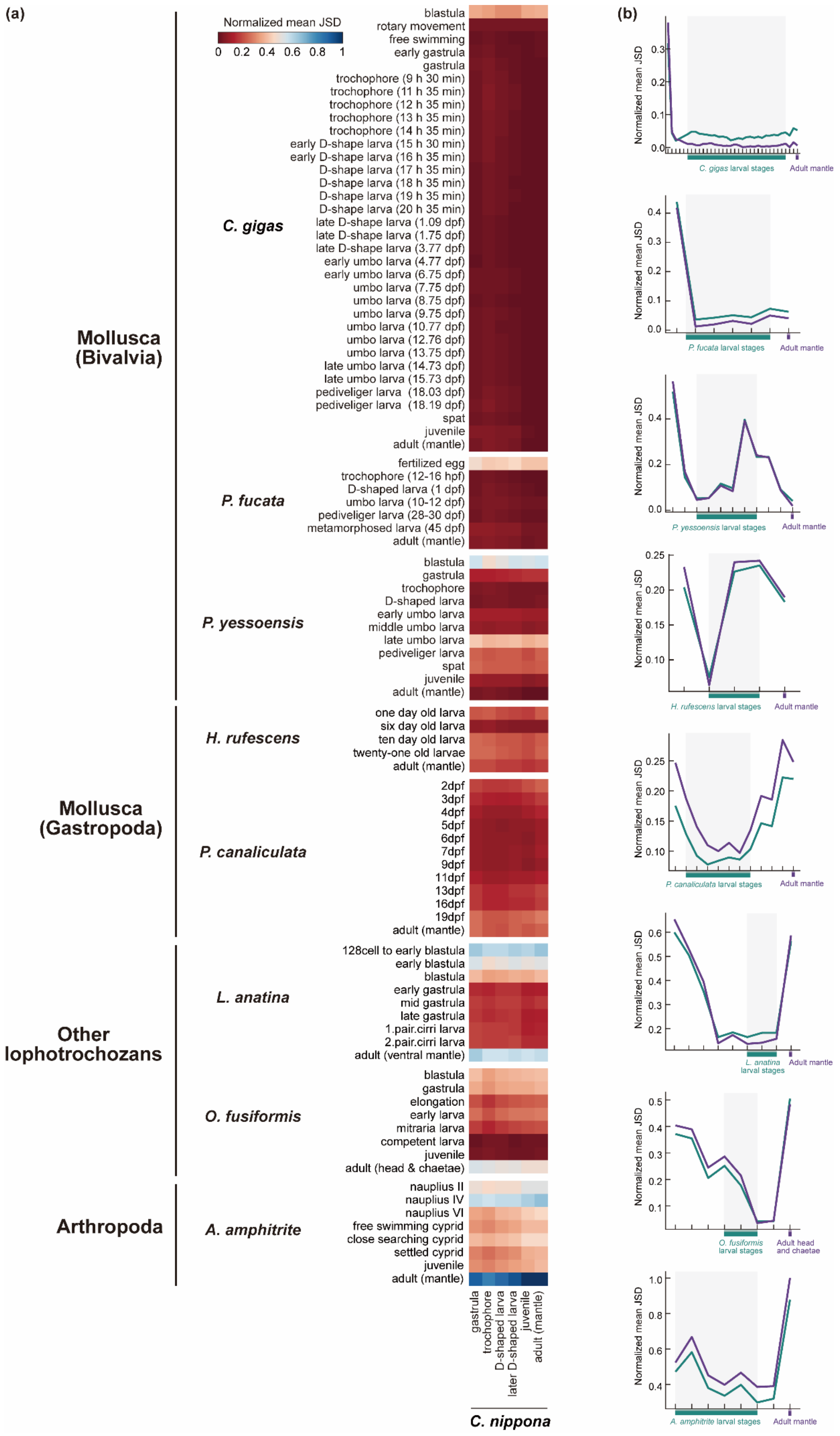

(a)
Normalized mean JSD
0 0.2 0.4 0.6 0.8 1
Mollusca (Bivalvia)
C. gigas
blastula
rotary movement
free swimming
early gastrula
gastrula
trochophore (9 h 30 min)
trochophore (11 h 35 min)
trochophore (12 h 35 min)
trochophore (13 h 35 min)
trochophore (14 h 35 min)
early D-shape larva (15 h 30 min)
early D-shape larva (16 h 35 min)
D-shape larva (17 h 35 min)
D-shape larva (18 h 35 min)
D-shape larva (19 h 35 min)
D-shape larva (20 h 35 min)
late D-shape larva (1.09 dpf)
late D-shape larva (1.75 dpf)
late D-shape larva (3.77 dpf)
early umbo larva (4.77 dpf)
early umbo larva (6.75 dpf)
umbo larva (7.75 dpf)
umbo larva (8.75 dpf)
umbo larva (9.75 dpf)
umbo larva (10.77 dpf)
umbo larva (12.76 dpf)
umbo larva (13.75 dpf)
late umbo larva (14.73 dpf)
late umbo larva (15.73 dpf)
pediveliger larva (18.03 dpf)
pediveliger larva (18.19 dpf)
spat
juvenile
adult (mantle)
P. fucata
fertilized egg
trochophore (12-16 hpf)
D-shaped larva (1 dpf)
umbo larva (10-12 dpf)
pediveliger larva (28-30 dpf)
metamorphosed larva (45 dpf)
adult (mantle)
P. yessoensis
blastula
gastrula
trochophore
D-shaped larva
early umbo larva
middle umbo larva
late umbo larva
pediveliger larva
spat
juvenile
adult (mantle)
Mollusca (Gastropoda)
H. rufescens
one day old larva
six day old larva
ten day old larva
twenty-one old larvae
adult (mantle)
P. canaliculata
2dpf
3dpf
4dpf
5dpf
6dpf
7dpf
9dpf
11dpf
13dpf
16dpf
19dpf
adult (mantle)
Other lophotrochozans
L. anatina
128cell to early blastula
early blastula
blastula
early gastrula
mid gastrula
late gastrula
1.pair.cirri larva
2.pair.cirri larva
adult (ventral mantle)
O. fusiformis
blastula
gastrula
elongation
early larva
mitraria larva
competent larva
juvenile
adult (head & chaetae)
Arthropoda
A. amphitrite
nauplius II
nauplius IV
nauplius VI
free swimming cyprid
close searching cyprid
settled cyprid
juvenile
adult (mantle)
gastrula
trochophore
D-shaped larva
later D-shaped larva
juvenile
adult (mantle)
C. nippona
(b)
Normalized mean JSD
C. gigas larval stages
Adult mantle
P. fucata larval stages
P. yessoensis larval stages
H. rufescens larval stages
P. canaliculata larval stages
L. anatina larval stages
O. fusiformis larval stages
Adult head and chaetae
A. amphitrite larval stages

**Supplementary Fig 20. Lophotrochozoan larvae share maximal transcriptional similarity of biomineralization TFs at early larval stages. (a)** Heatmaps of normalized Jensen-Shannon divergence (JSD) from pairwise comparisons of 29 single copy one-to-one TF orthologs between *C. nippona* and eight bilaterians. **(b)** Average relative JSD for the stages of minimal divergence to the D-shape larva stage (green) and adult mantle tissues (purple) of *C. nippona* in **(a)**.

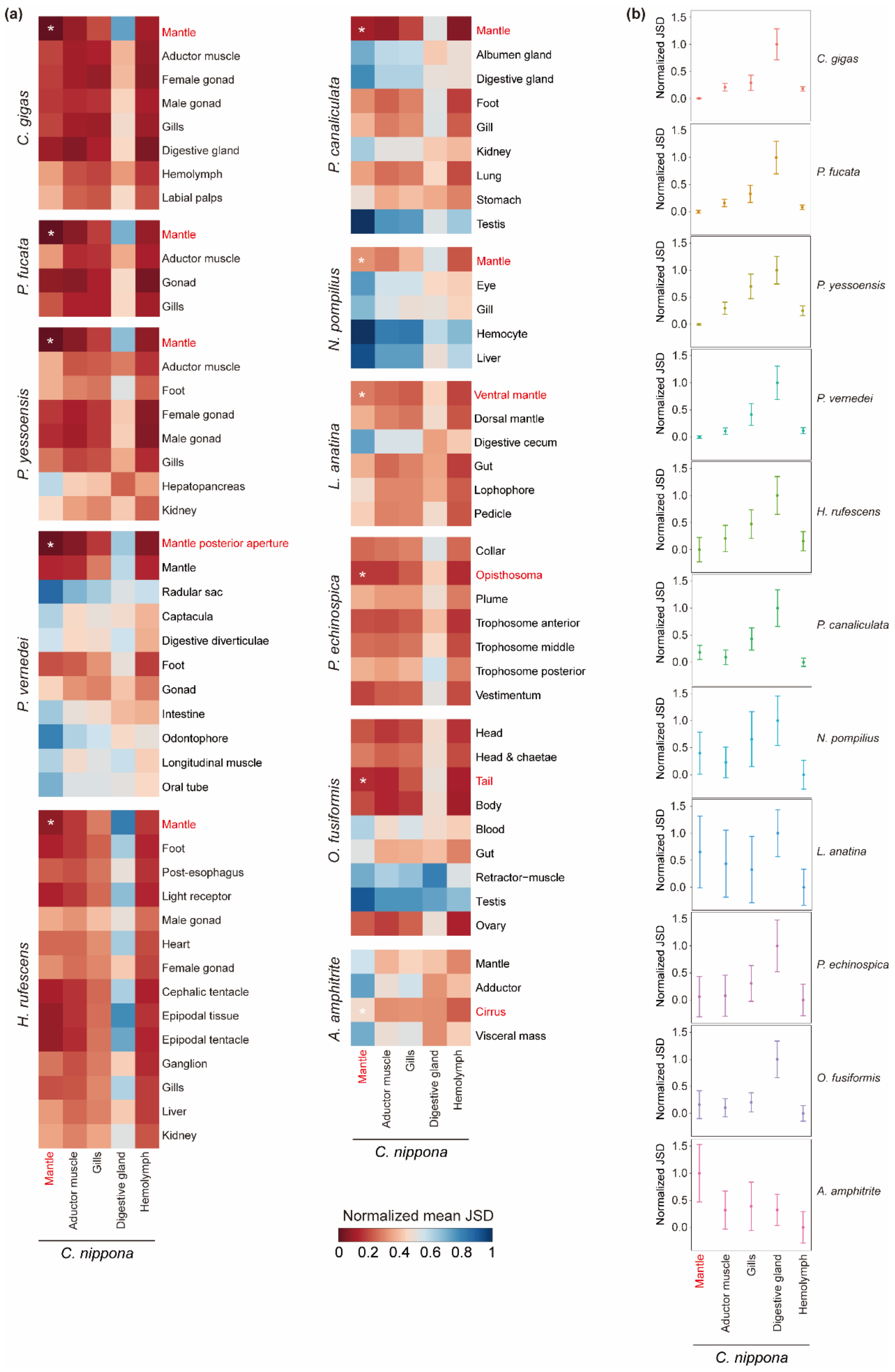


**Supplementary Fig 21. Molluscan mantles share maximal transcriptional similarity of biomineralization TFs. (a)** Heatmaps of normalized JSD from pairwise comparisons of 29 single copy one-to-one TF orthologs between *C. nippona* and 11 bilaterians. Asterisk indicates the tissue of minimal JSD of each species to the mantle of *C. nippona*. **(b)** Average relative JSD for the datasets shown in **(a)** from tissues of minimal JSD to each *C. nippona* tissue. Confidence intervals represent the standard deviation from 1,000 bootstrap replicates of the ortholog sets.

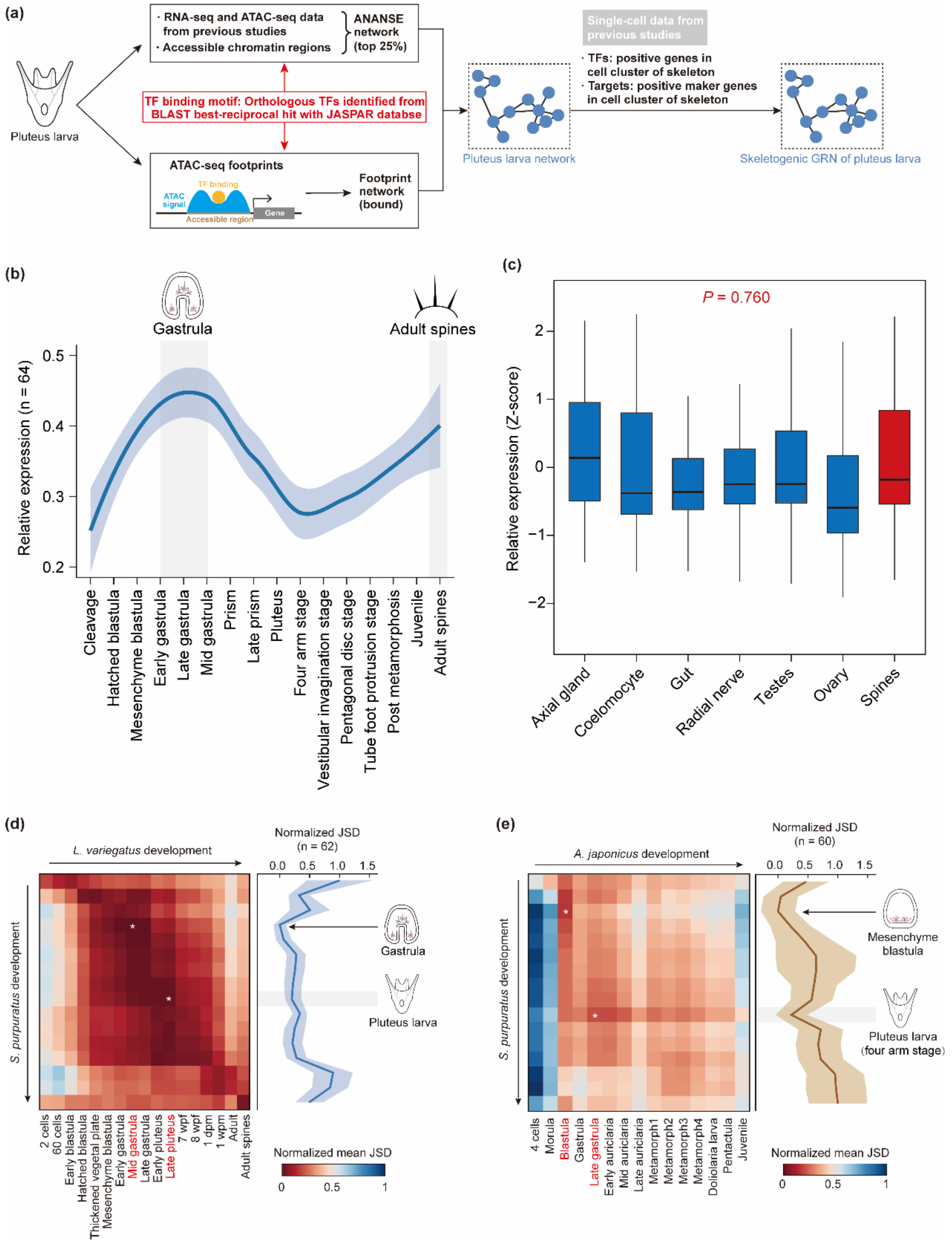


**Supplementary Fig 22. The sea urchin *Strongylocentrotus purpuratus* shares high transcriptional similarity of biomineralization TFs with two other echinoderms during early embryonic and larval development. (a)** Schematic overview of the pipeline used to reconstruct of skeletogenic GRN in *S. purpuratus* larvae. **(b-c)** Gene expression patterns of 64 biomineralization TFs during *S. purpuratus* development **(b)**

and across adult tissues **(c)**. A two-sided Wilcoxon rank-sum test was used to assess significance between expression levels in spines and other tissues. **(d-e)** JSD-based transcriptional similarity of biomineralization TFs from pairwise comparisons of *S. purpuratus* with *Lytechinus variegatus* and *Apostichopus japonicus* across developmental stages, respectively. The highest transcriptomic similarity between the two sea urchins occurs at the gastrulation, whereas between *S. purpuratus* and *A. japonicus*, it peaks at the mesenchyme blastula stage. The second-highest similarity stage is indicated with a grey background.

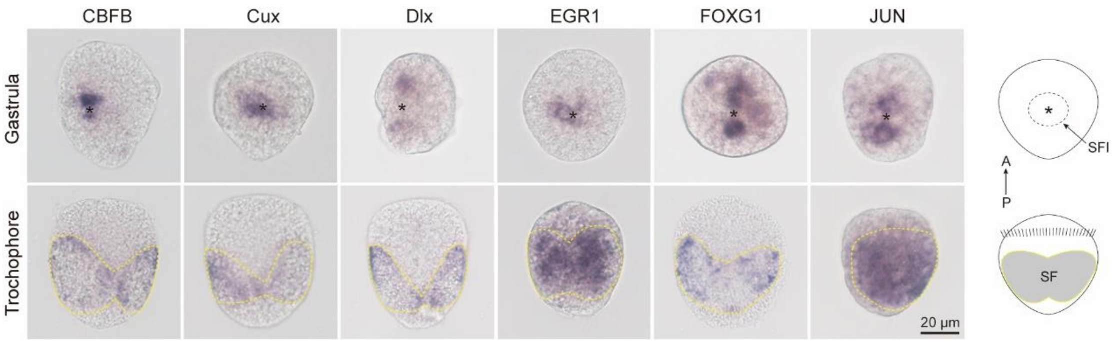


**Supplementary Fig 23. Whole-mount ISH results (dorsal views) of six conserved biomineralization-related TFs in the gastrula and trochophore of *C. nippona*.** The most part of the shell field is invaginated during gastrulation and the asterisks indicate the opening. Yellow dashed lines indicate the shell field (SF) region in trochophore. Right: schematic diagrams of gastrula (top) and trochophore (bottom) in dorsal view. Abbreviations: A anterior; P posterior; SFI shell field invagination.

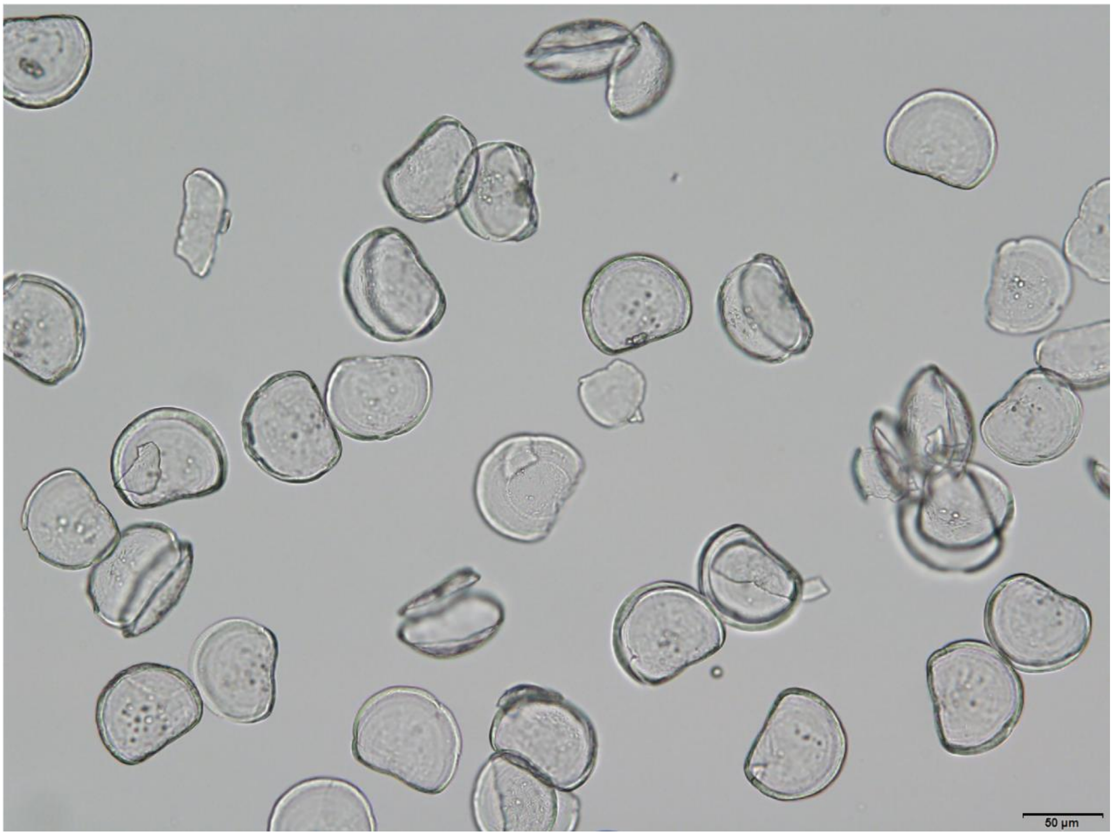


**Supplementary Fig 24. Larval shells after cleaning.**

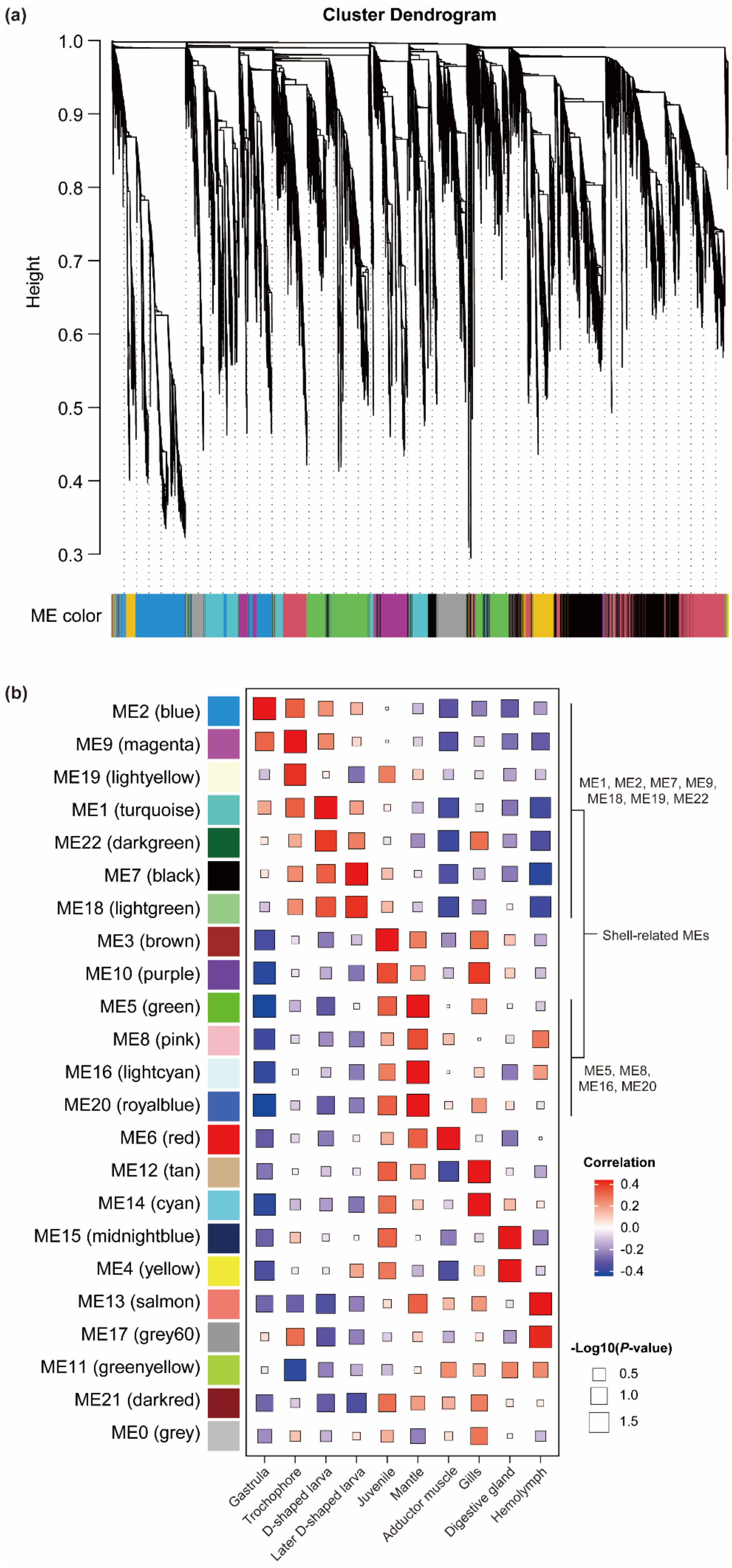
(a)
Cluster Dendrogram
Height
1.0
0.9
0.8
0.7
0.6
0.5
0.4
0.3
ME color
(b)
ME2 (blue)
ME9 (magenta)
ME19 (lightyellow)
ME1 (turquoise)
ME22 (darkgreen)
ME7 (black)
ME18 (lightgreen)
ME3 (brown)
ME10 (purple)
ME5 (green)
ME8 (pink)
ME16 (lightcyan)
ME20 (royalblue)
ME6 (red)
ME12 (tan)
ME14 (cyan)
ME15 (midnightblue)
ME4 (yellow)
ME13 (salmon)
ME17 (grey60)
ME11 (greenyellow)
ME21 (darkred)
ME0 (grey)
Gastrula
Trochophore
D-shaped larva
Later D-shaped larva
Juvenile
Mantle
Adductor muscle
Gills
Digestive gland
Hemolymph
ME1, ME2, ME7, ME9, ME18, ME19, ME22
Shell-related MEs
ME5, ME8, ME16, ME20
Correlation
0.4
0.2
0.0
-0.2
-0.4
-Log10(P-value)
0.5
1.0
1.5

**Supplementary Fig 25. WGCNA analysis of expressed genes (TPM > 1) across developmental stages and adult tissues in *C. nippona*. (a)** Module construction of gene-expression network for 30 samples. **(b)** Correlation matrix between MEs and developmental stages or adult organs. MEs associated with larval stages (ME1, ME2, ME7, ME9, ME18, ME19, and ME22) and the adult mantle (ME5, ME8, ME16, and ME20) contain biomineralization effector genes that are considered to be involved in shell formation (correlation valve > 0 and *P* value < 0.05).

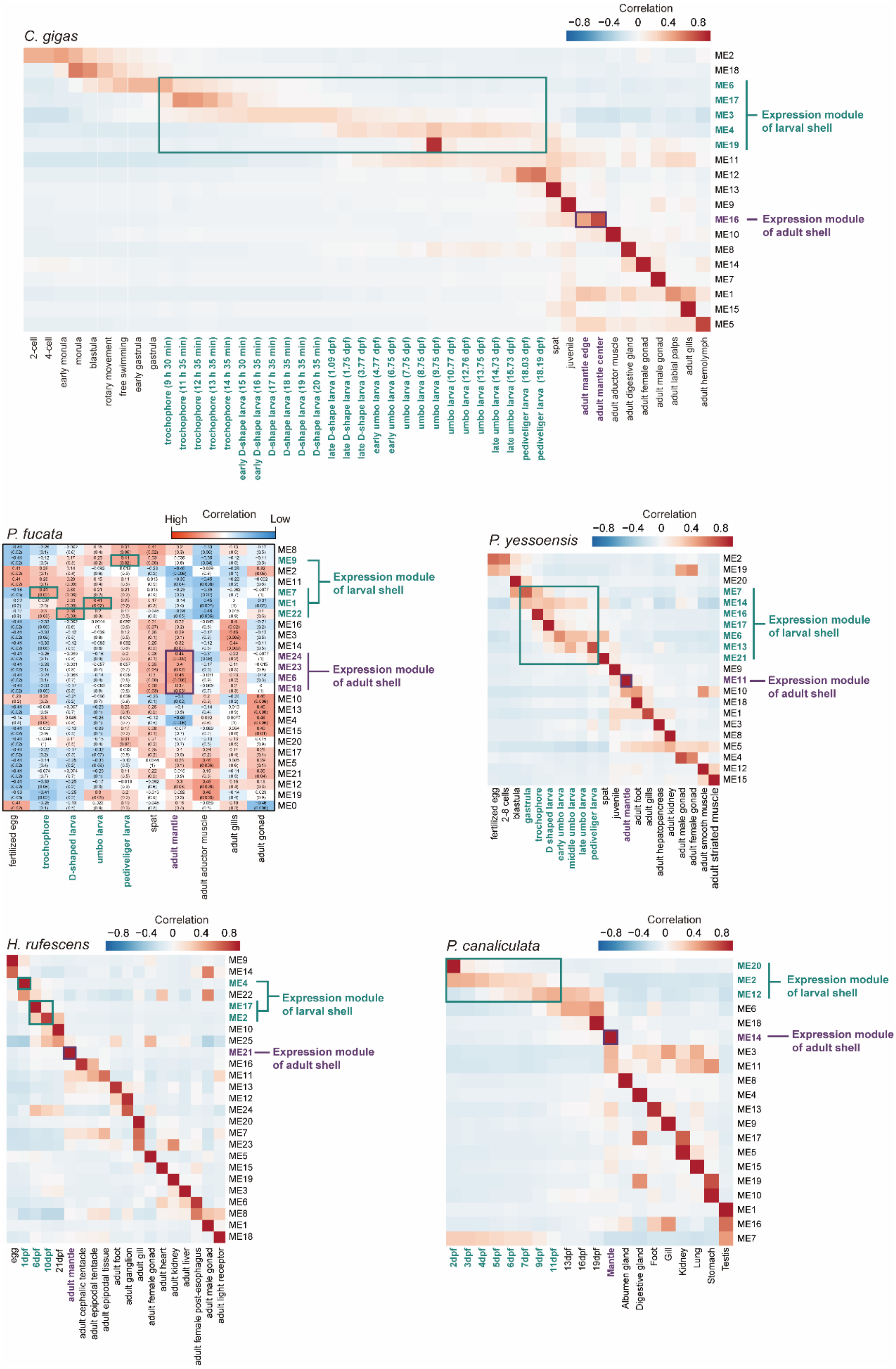


**Supplementary Fig 26. WGCNA analysis of expressed genes (TPM > 1) across developmental stages and adult tissues in five other molluscs.** MEs associated with larval stages (green) and the adult mantle (purple) contain biomineralization effector genes that are considered to be involved in shell formation.

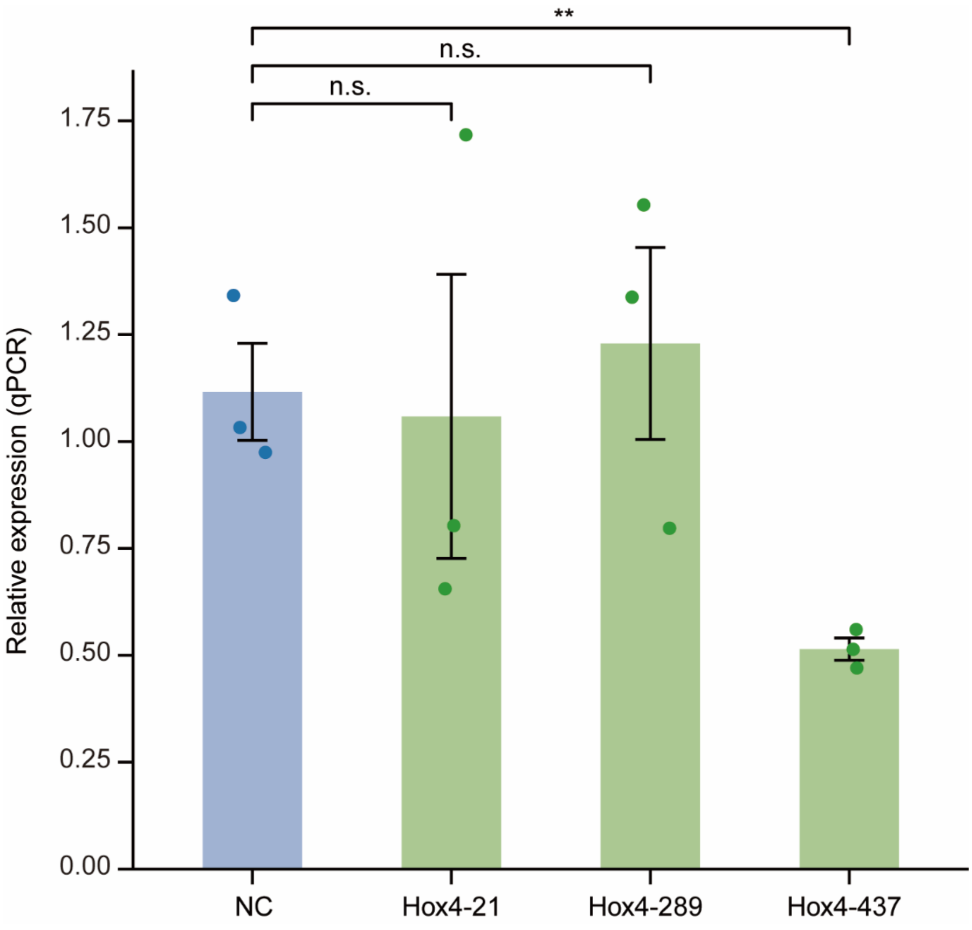


**Supplementary Fig 27. Results of the RNA interference (RNAi) pilot experiment.**

Three small interfering RNA (siRNA) strands targeting *Hox4* were tested by measuring the expression level of *Hox4* following the RNAi procedure described in the Methods section of the main text. The results indicated the siRNA strand Hox4-437 was the most effective one. Real-time quantitative polymerase chain reaction (RT-qPCR) was performed using cDNA from mantle tissues of *C. nippona* individuals injected with siRNA three times over a six-day period following shell-drilling ($n = 3$; mean ± SE; two-sided Student's t-test: $^{**}P < 0.01$; n.s., no significance).

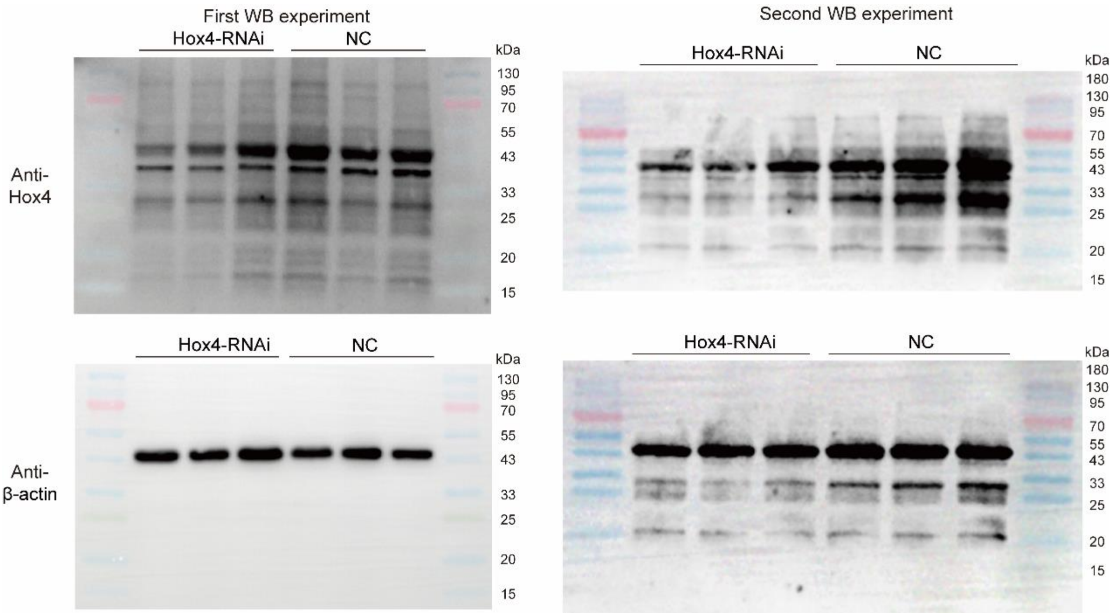


**Supplementary Fig 28. Western blot (WB) analysis of Hox4 protein abundance in the mantle tissues (n = 3) after RNAi experiment.** The experiment was repeated twice and yielded consistent results (Supplementary Data 26). For each replicate, the same membrane was sequentially incubated with anti-Hox4 and anti-β-actin antibodies after washing.
